# Cellular and molecular landscapes of human tendons across the lifespan revealed by spatial and single-cell transcriptomics

**DOI:** 10.1101/2025.06.19.660575

**Authors:** Alina Kurjan, Jolet Y. Mimpen, Lorenzo Ramos-Mucci, Ali C. Aksu, Christopher D. Buckley, Adam P. Cribbs, Mathew J. Baldwin, Sarah J. B. Snelling

## Abstract

Tendon injuries are common and often heal poorly. While developing tendons heal without scarring, this capacity declines with age, yet the underlying cellular transitions remain poorly defined. Here, we integrate histological, single-nucleus, single-cell, and spatial transcriptomic profiling of human Achilles and quadriceps tendons across embryonic, foetal, and adult stages, including ruptured adult tendons. We identify seven embryonic progenitor states that give rise to three distinct tendon-associated fibrillar, connective tissue, and chondrogenic lineages. These populations diversify during development and occupy distinct spatial niches, adopting specialised roles in matrix synthesis, tissue remodelling, and mechanical adaptation. While non-fibroblast populations remain transcriptionally stable with age, fibroblasts undergo marked reprogramming, shifting to homeostatic or injury-responsive states. In ruptured adult tendons, a subset of fibroblasts partially reactivates developmental programs but remains transcriptionally distinct from their regenerative counterparts. These findings define the cellular architecture of human tendon development and ageing and reveal lineage-specific targets for therapeutic repair.

## INTRODUCTION

Tendons are essential connective tissues that transmit force between muscles and bones, enabling movement and stabilising joints. Despite their remarkable strength and resilience, tendons are vulnerable to injuries and degenerative conditions (often termed tendinopathies), which cause pain, swelling, and impaired function, severely affecting quality of life.^1–4^

The capacity for tendon repair varies drastically across the lifespan. Whereas foetal and neonatal tendons can regenerate with minimal scarring, adult tendons typically heal through fibrosis or ectopic ossification, resulting in disorganised extracellular matrix (ECM) and compromised mechanical properties.^5–10^ These contrasting outcomes appear driven by intrinsic differences in tendon-resident cell behaviour, rather than external environmental factors.^6,8,11–17^

Recent spatial and single-cell/nucleus RNA-sequencing studies have begun to uncover the cellular heterogeneity underlying these differences, though findings vary across species and anatomical locations.^13,18–26^ Beyond the well-characterised *COL1A1*^+^*TNMD*^+^*MKX^+^*intrafascicular tenocytes responsible for ECM production and fibre alignment,^27–30^ tendons also harbour fibroblasts with elevated *COL3A1* expression,^20–26,31^ *PTPRC/CD45*^+^ immune cells,^13,19–22,24–26^ *PECAM1/CD31*^+^ endothelial cells,^13,19–26^ *MCAM*/CD146^+^ pericytes,^19,26^ *NOTCH3*^+^ mural or *ACTA2*^+^ smooth muscle cells,^20,21,24–26^ and, in some studies, muscle cells,^21,24,26^ neural cells,^19,24,26^ adipocytes,^24,26^ and fibro-adipogenic progenitors.^13,20^ Despite this emerging complexity, consistent identification of canonical tendon stem/progenitor cells (TSPCs) – marked by *STRO-1*, *MCAM*/*CD146*, *ENG*/*CD105*, *THY1/CD90*, *CD44*, *SCX*, *TNMD*, *COMP*, and *TNC*^32,33^ – remains elusive. Instead, recent studies describe injury-responsive, sheath-derived progenitors expressing *SCX*^-^ *TPPP3*^+^*PDGFRA*^+^ ^13,34^ or *AXIN2^+^* ^12,17^ signatures in adult tendons.

Evidence from murine models highlights that tendon healing is age- and context-dependent. Neonatal tendons exhibit robust regenerative capacity driven by intrinsic *Scx*^+^ cells,^9^ while adult tendons predominantly heal through fibrotic mechanisms, involving extrinsic *Scx*^-^*Acta2*^+9^ or *Sca1*^+35^ cells. Mechanical load and TGF-β signalling further modulate *Scx* expression and ECM organisation, shaping whether repair proceeds via regenerative or fibrotic pathways.^9,12,35–45^ Yet, major gaps remain regarding how tendon cell identity, plasticity, and niche-specific behaviours evolve over time from development into adulthood, particularly following injury. Moreover, whether developmental tendon lineages are retained, reactivated, or replaced after rupture in adult humans is not yet understood.

Here, we map the transcriptional landscapes of human tendons across development and ageing. Using single-nucleus, single-cell, and spatial RNA-sequencing, we profile cell populations within embryonic (6-9 post-conception weeks (pcw)), foetal (12-20 pcw), and adult (25-76 years) human tendons. We further characterise how rupture remodels the cellular composition and cell state in adult quadriceps tendons. This study identifies conserved and divergent fibroblast populations across life stages, revealing signatures of intrinsically regenerative fibroblasts and defining how ageing and injury reshape tendon cell fate and function.

## RESULTS

To characterise cell populations in intrinsically (re)generative human tendons, we performed single-nucleus RNA sequencing (snRNA-seq) on Achilles (*N*=8) and quadriceps tendons (*N*=7) from nine human foetal donors aged 12, 17, and 20 pcw (Figure 1A, Table S1). In parallel, spatial transcriptomics was conducted on Achilles (*N*=1) and quadriceps tendons (*N*=2) from a single 20 pcw donor (Figure 1B). Following data processing and quality control, scVI-integrated^46^ snRNA-seq data from 91,859 nuclei were clustered and annotated, identifying distinct fibroblasts, chondrocytes, immune, endothelial, muscle, and nervous system cells (Figure 1C) across the ages (Figure 1D). Cell2location^47^ analysis mapped these snRNA-seq cell type signatures to their likely spatial locations within the tissues (Figure 1F).

**Figure 1.**
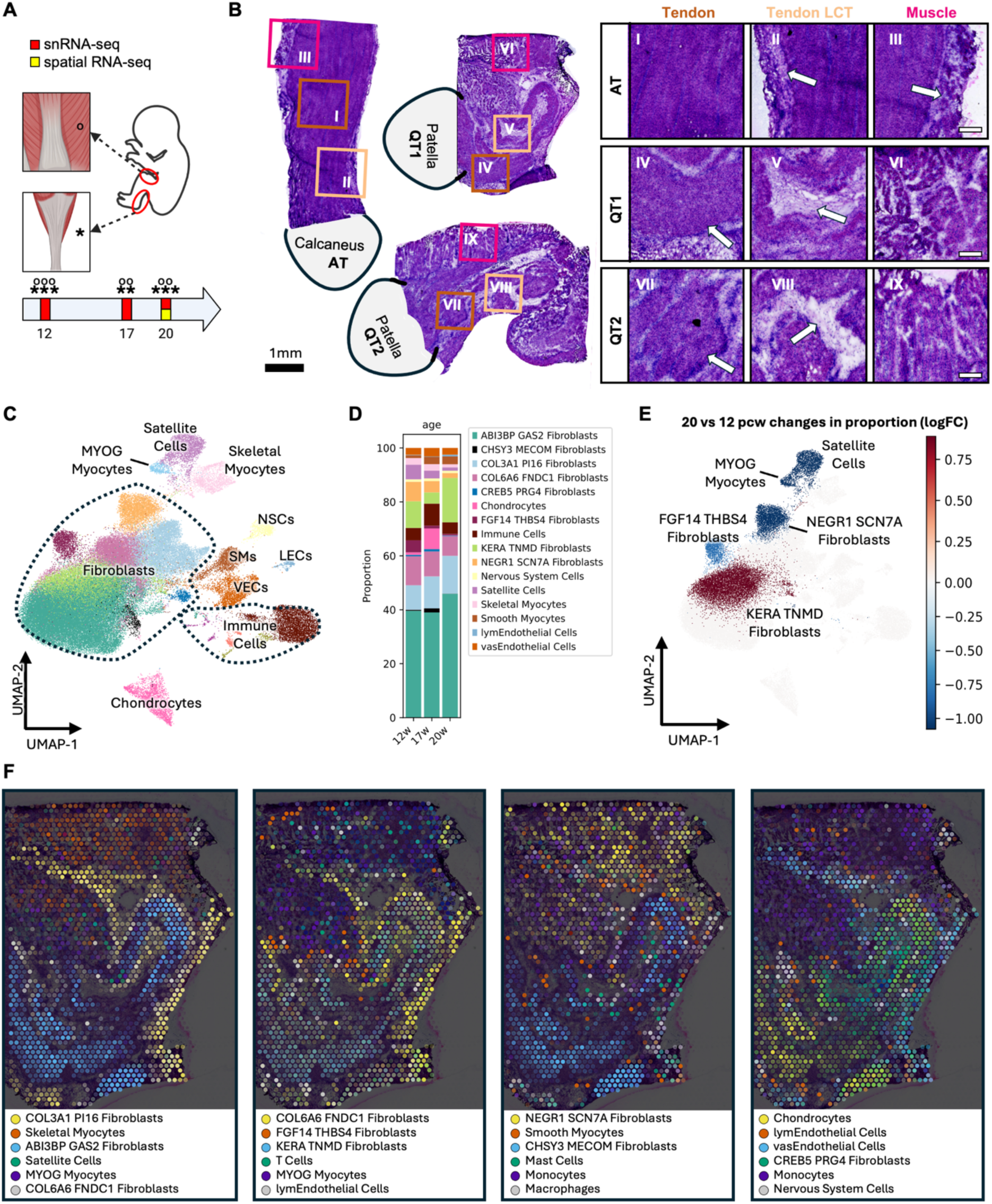
Experimental design, histological and spatial profiling, and compositional shifts in developing human tendons. (A) Overview of experimental design. Stars and circles indicate numbers of Achilles and quadriceps tendons analysed per timepoint, respectively. (B) H&E-stained cryosections (10 μm) of Achilles tendon (AT) and two quadriceps tendons (QT1, QT2) used for 10X Visium spatial RNA-seq. Tissue orientation is annotated by adjacent bone structures (calcaneus, patella). Black scale bar = 1 mm. Insets show zoomed regions (white boxes) with anatomical features annotated: tendon body, loose connective tissue (LCT), and muscle. White scale bars = 250 μm. (C) UMAP of scANVI-integrated 12-20 post-conception week (pcw) foetal tendon snRNA-seq data, showing annotated cell types. VECs: vascular endothelial cells; LECs: lymphatic endothelial cells; SMs: smooth myocytes; NSCs: nervous system cells. (D) Bar plot of cell type proportions by foetal age. (E) scCODA-inferred compositional changes from 12 to 20 pcw. UMAP highlights significant increases (red) and decreases (blue) in cell type proportions. (F) Cell2location spatial mapping of 20 pcw quadriceps tendon sections, showing relative abundance of each cell type.

### Single-nucleus and spatial RNA-sequencing reveal heterogenous compartmentalised fibroblasts within foetal regenerative tendons

Fibroblasts dominated the cellular landscape of foetal tendons, forming at least eight transcriptional cell states across a few closely related types (Figure 2A). Three populations – termed ABI3BP GAS2, KERA TNMD, and CHSY3 MECOM Fibroblasts – shared transcriptional similarities, indicating a common lineage with functional divergence marked by key gene expression profiles (Figure 2B). The largest, ABI3BP GAS2 Fibroblasts, were defined by high levels of *COL11A1*^48^ alongside ECM- and cytoskeletal-regulators *ABI3BP*, *GAS2, SOX5*, *EXT1* and *PLEKHH2,* consistent with roles in matrix assembly and cellular adhesion (Figure 2B,C). Transcriptionally similar to these, KERA TNMD Fibroblasts were enriched for collagens (*COL1A1*, *COL1A2*, *COL6A3*, *COL6A1*, *COL12A1*) and matrix-associated genes (*SPARC*, *POSTN*, *FMOD*, *KERA*), with low *MKX* and high *TNMD* expression (Figure 2B,C), suggesting a more specialised, mature fibroblast phenotype.^49–55^ CHSY3 MECOM Fibroblasts, in contrast, expressed ECM-hydration regulator *CHSY3*, transcriptional regulators *MECOM* and *FOXP2*, and mineralisation-associated *COL24A1,*^56^ *SMOC1*,^57^ and *ENPP1*,^58^ as well as chondrogenic and enthesis markers *SOX6*^30^ and *COL27A1*^31^ (Figure 2B), respectively, suggesting a role in tendon–bone interface remodelling and mechanoadaptation.

**Figure 2.**
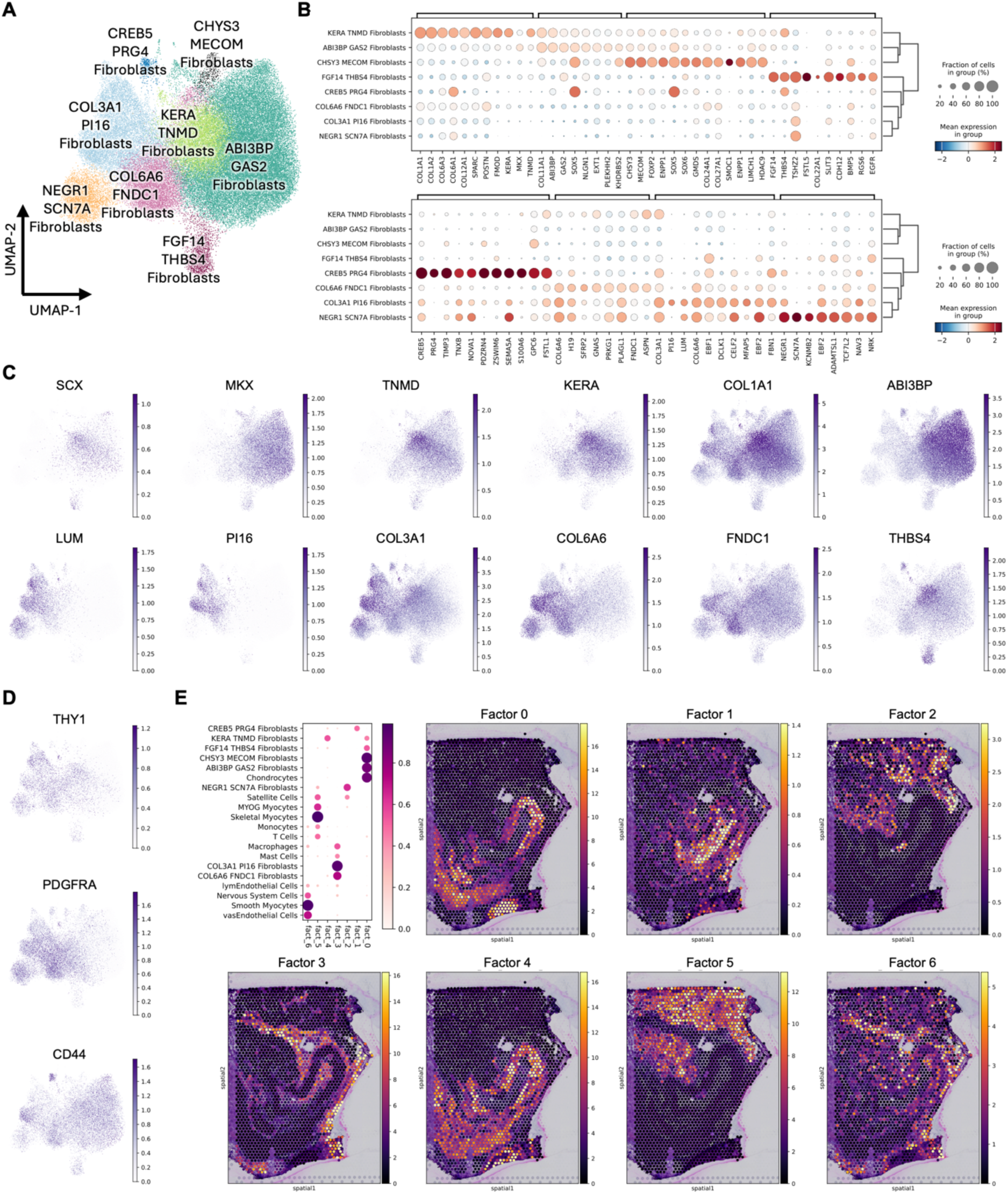
Transcriptional diversity, marker expression, and spatial localisation of fibroblast subtypes in foetal human tendons. (A) UMAP of annotated fibroblast subtypes from 12–20 post-conception week (pcw) foetal tendon snRNA-seq data. (B) Dotplots of log1pPF-normalised and scaled gene expression showing hierarchically clustered, differentially expressed genes across fibroblast subtypes. Dot size indicates cell type abundance. (C) UMAPs showing normalised expression of selected fibroblast marker genes across annotated subtypes. (D) UMAPs showing expression of selected tendon stem/progenitor cell (TSPC) markers: THY1 (CD90), PDGFRA, and CD44. (E) Unsupervised non-negative matrix factorisation (NMF) analysis of spatial transcriptomics cell2location output across three 20 pcw tendon samples. Dot plot displays NMF factors derived from mean normalised UMI counts per factor (dot size and colour reflect gene expression and loading strength). Gene loadings indicate the contribution of individual genes to each colocalised fibroblast factor. Spatial scatter plots show cell densities (mean normalised UMI counts) per NMF factor from one representative 20 pcw quadriceps tendon section.

Spatial transcriptomic analysis localised these three transcriptionally related fibroblast populations within the main fascicular bodies of 20 pcw Achilles and quadriceps tendons (Figures 1F, 2E-Factor 0). High expression of canonical TSPC markers *ENG*, *THY1*, *CD44*, and *NES*^32^ was specifically enriched within spatially-resolved KERA TNMD Fibroblasts (Figure S1).

Temporal pseudobulk differential gene expression (DGE) analysis between 12 and 20 pcw, followed by Gene Ontology Biological Process (GO:BP) enrichment, revealed that by 20 pcw, ABI3BP GAS2, KERA TNMD, and CHSY3 MECOM fibroblasts upregulate genes involved in cell growth, metabolism, pattern formation, Wnt signalling regulation, histone methylation, and stem cell differentiation (Table S2).

These populations also showed increased expression of genes linked to immune system modulation, while downregulating cell division and contractile programs, including those related to muscle contraction and myofibril assembly. These findings suggest that these foetal fibroblasts not only promote tendon matrix formation but also help modulate the local immune environment and suppress contractile differentiation during tendon development.

A second major group of transcriptionally related fibroblast states was characterised by high expression of *COL3A1*, *COL6A6*, *PDGFRA*, *DCLK1*, *TSHZ2*, *PLAGL1*, and *VCAN*, and comprised three dominant subtypes: COL3A1 PI16, COL6A6 FNDC1, and NEGR SCN7A Fibroblasts (Figure 2B). The most abundant, COL3A1 PI16 Fibroblasts, exhibited high expression of *COL3A1*, *PI16*, *EBF1, DCLK1*, *LUM*, *MFAP5* and *FBN1* (Figure 2B,C), and localised to the loose connective tissue (LCT) regions of Achilles and quadriceps tendons (Figures 1F, 2E). These cells also expressed sheath TSPC markers *TPPP3* and *PDGFRA*^13,34^ (Figure S1). From 12 to 20 pcw, they upregulated pathways related to Hippo signalling, chondrocyte and epithelial development, and suppression of vascular smooth muscle proliferation, while downregulating programs associated with muscle development, stress responses, apoptosis, and epigenetic modification (Table S2). These features suggest a role in early ECM organisation and structural maintenance within the tendon LCT.

A transcriptionally similar population, COL6A6 FNDC1 Fibroblasts, was distinguished by elevated expression of *COL6A6*, *FNDC1*, and regulators of chondrogenesis and mineralisation *SFRP2*,^61^ *GNAS*,^62^ and *ASPN*^63–65^ (Figure 2B). Like COL3A1 PI16 Fibroblasts, they resided within the LCT regions (Figures 1F, 2E). Between 12 and 20 pcw, they upregulated pathways involved in tissue repair while downregulating processes linked to cell adhesion, miRNA regulation, synaptic function, metabolism, and vascular development (Table S2). These signatures point to a role in maintaining tendon structural integrity and modulating immune responses during growth.

NEGR1 SCN7A Fibroblasts were characterised by high expression of *NEGR1*, *SCN7A*, *EBF2*, *NRK*, and *KCNMB2*, suggesting involvement in fibroblast mechanosensitivity. They also expressed *NAV3* and *ADAMTSL1*, linked to cytoskeletal and ECM organisation, respectively (Figure 2B). Spatial mapping located these cells between skeletal myocytes within tendon-adjacent muscle LCT regions (Figures 1F, 2E, 3E), implicating them in myotendinous junction (MTJ) organisation, mechanotransduction, and matrix remodelling.

Another distinct fibroblast population, FGF14 THBS4 Fibroblasts, occupied the tendon–muscle boundaries (Figure 1F) and expressed MTJ-associated genes *COL22A1*^66,67^ and *THBS4*, alongside high levels of *FGF14, SLIT3, TSHZ2, CDH12, BMP5, RGS6, EGFR,* and *FSTL5* (Figure 2B,C). From 12 to 20 pcw, these cells upregulated pathways involved in ECM synthesis, angiogenesis, immune interactions, and neurogenesis, while downregulating BMP signalling, ossification, and skeletal muscle proliferation pathways (Table S2), supporting a role in maintaining a specialised MTJ-supporting fibroblast identity.

Finally, a small CREB5 PRG4 Fibroblast population localised within LCT regions (Figures 1F, 2E) expressed synovium- and cartilage-associated *CREB5,*^68,69^ alongside lubricating *PRG4*.^70–75^ These cells also showed high levels of *TIMP3*, *TNXB*, and *NOVA1* (Figure 2B), suggesting roles in ECM maintenance, collagen organisation, and tendon lubrication, particularly in regions subject to mechanical stress.

### Vascular, immune, and neural niches emerge alongside fibroblasts to structure developing human tendons

Non-fibroblast populations identified in foetal tendons included vascular and lymphatic endothelial cells, smooth muscle cells, nervous system-associated cells, immune cells, and myocytes (Figure 3A). Vascular endothelial cells expressed *PECAM1*, *CD34*, and *VWF*, while lymphatic endothelial cells were marked by *LYVE1*, *PROX1*, and *FLT4* (Figure 3B,C). Smooth muscle cells showed high expression of *ACTA2*, *MYH11*, *CALD1*, *NOTCH3*, and *PDGFRB* (Figure 3B). Nervous system-associated cells were distinguished by *NRXN1*, *NCAM2*, and *SOX10* expression, displaying transcriptomic features consistent with glial cells (Figure 3B). All these cell types co-localised within the tendon LCT regions (Figures 1F, 2E, 3E).

**Figure 3.**
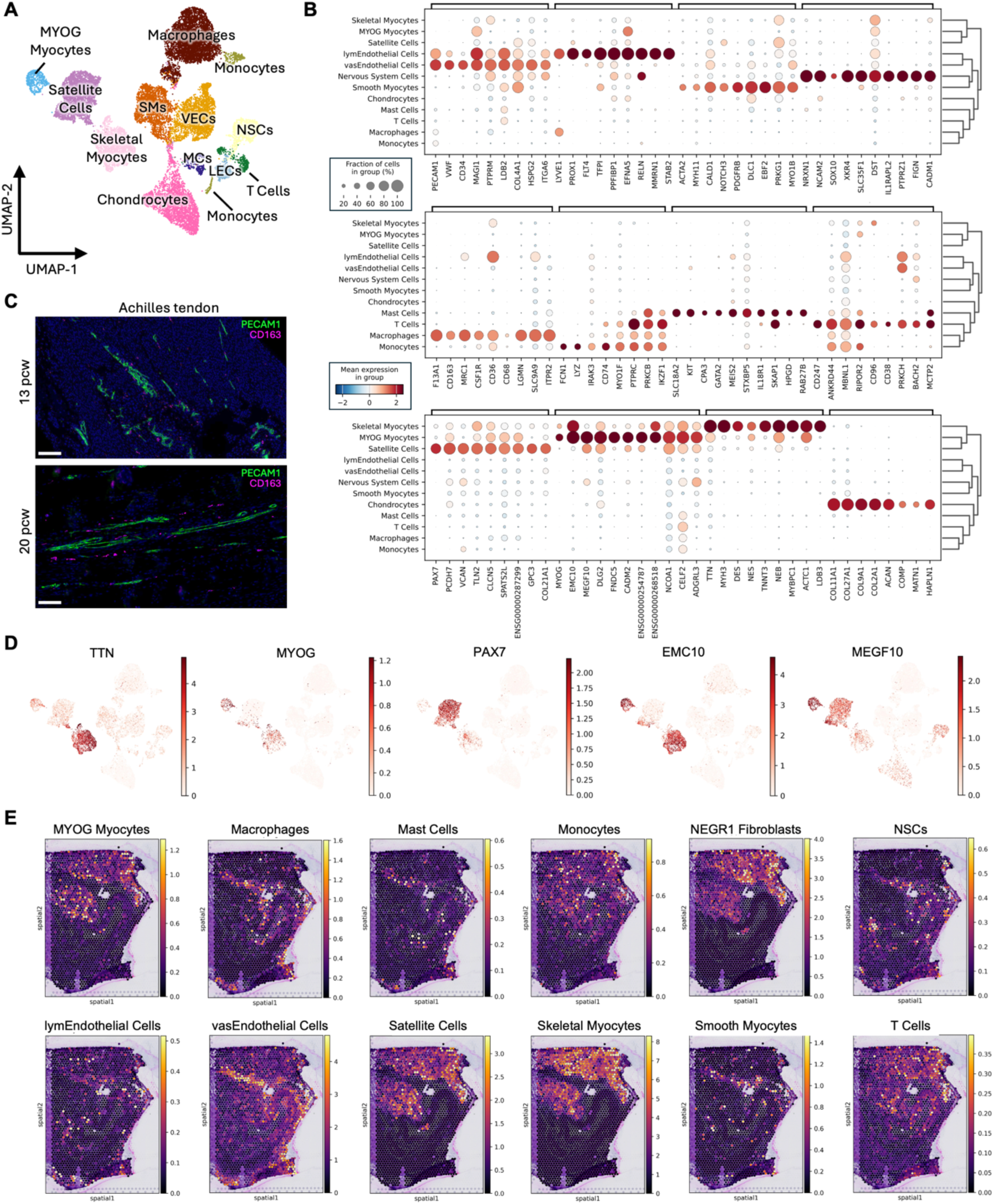
Characterisation and spatial mapping of non-fibroblast cell populations in developing human tendons. (A) UMAP of annotated non-fibroblast cell types from 12–20 post-conception week (pcw) foetal tendon snRNA-seq data. VECs: vascular endothelial cells; LECs: lymphatic endothelial cells; SMs: smooth myocytes; NSCs: nervous system cells; MCs: mural cells. (B) Dotplots of log1pPF-normalised and scaled gene expression showing hierarchically clustered, differentially expressed genes across non-fibroblast cell types. Dot size indicates cell type abundance. (C) Immunofluorescence images of 20 pcw Achilles tendon sections stained for PECAM1 (green, endothelial cells) and CD163 (magenta, macrophages). White scale bars = 100 µm. (D) UMAPs showing normalised expression of selected myocyte marker genes. (E) Cell2location spatial mapping of cell types in 20 pcw quadriceps tendon sections, with colour intensity indicating relative abundance at each location.

Immune cells broadly expressed *PTPRC*, *CD44*, *IKZF1*, *RUNX1*, *DOCK2*, and *INPP5D*. The predominant subset expressed *F13A1*, *CD163*, *MRC1* (*CD206*), *CSF1R*, *CD36*, and *LGMN*, consistent with macrophage identity (Figure 3B). Additional immune subsets included monocytes (*FCN1*, *LYZ*, *IRAK3*), mast cells (*KIT*, *CPA3*, *GATA2*) and T cells (*SKAP1*, *CD247*, *CD96*, *CD38*) (Figure 3B). While monocytes and T cells were predominantly co-localised with myocytes in muscle regions, mast cells and macrophages were enriched within the tendon LCT, co-localising with COL3A1 PI16 and COL6A6 FNDC1 fibroblasts (Figure 2E-Factors 3,5).

Three myocyte clusters were identified within muscle tissue adjoining the tendon MTJ (Figures 1F, 2E, 3E). One cluster expressed *PAX7*, marking satellite cells, while a second expressed markers of differentiated skeletal myocytes including *DES*, *NES*, *TNNT3*, *TTN*, *MYH3*, *COL22A1*, and *TNNC1* (Figure 3B,D). A third small cluster, expressing *MYOG*, *FNDC5*, *EMC10*, *MEGF10,* and overlapping markers from both satellite and skeletal myocytes, likely represented a transitional state (Figure 3B,D).

Finally, in one 17pcw quadriceps tendon sample, a discrete population expressing *COL2A1*, *COL9A1*, *ACAN*, *COMP*, *MATN1*, and *HAPLN1* was detected, consistent with a chondrocyte identity (Figure 3B). These cells co-localised with the fascicular fibroblasts within the main fibrillar tendon body (Figures 1F, 2E-Factor0), likely reflecting minor contamination from adjacent patellar cartilage.

### Second trimester foetal tendon populations shift from MTJ- and muscle-associated to tendon-building cells

To quantify changes in cell type proportions over time, we applied single-cell compositional data analysis (scCODA),^76^ a Bayesian modelling approach that accounts for sampling biases inherent to single-nucleus sequencing. Nervous System Cells were used as a reference group, assuming proportional stability between 12 and 20 pcw. At 20 pcw, compositional analysis revealed a significant decrease in MTJ- and muscle-associated populations, including FGF14 THBS4 Fibroblasts, MYOG Myocytes, Satellite Cells, and NEGR1 SCN7A Fibroblasts (Figure 1D). In contrast, KERA TNMD Fibroblasts – resembling differentiated tenocytes and expressing markers of fascicular TSPCs – showed a substantial proportional expansion during this period (Figure 1D).

### Single-cell and spatial transcriptomics of embryonic limbs reveal candidate progenitors for foetal tendon fibroblast lineages

The initiation of cell lineage decisions during early embryogenesis is crucial for establishing the cellular diversity required for tendon development. To investigate the origins of foetal tendon populations, we analysed published spatial and single-cell RNA-sequencing datasets from 6-9 pcw human embryonic limbs.^77^ Tendon regions were manually delineated within an 8 pcw H&E-stained spatial RNA-seq sample (Figure S2), and gene expression profiles were extracted as a reference for cell identification in the corresponding scRNA-seq dataset. A random forest classifier trained on these profiles identified 4,318 tendon-like cells out of 108,617 total cells (Figure S3). After filtering and batch correction, seven cell state clusters were annotated across 3,092 cells from six donors aged 6.5-9.3 pcw (Figure 4A).

**Figure 4.**
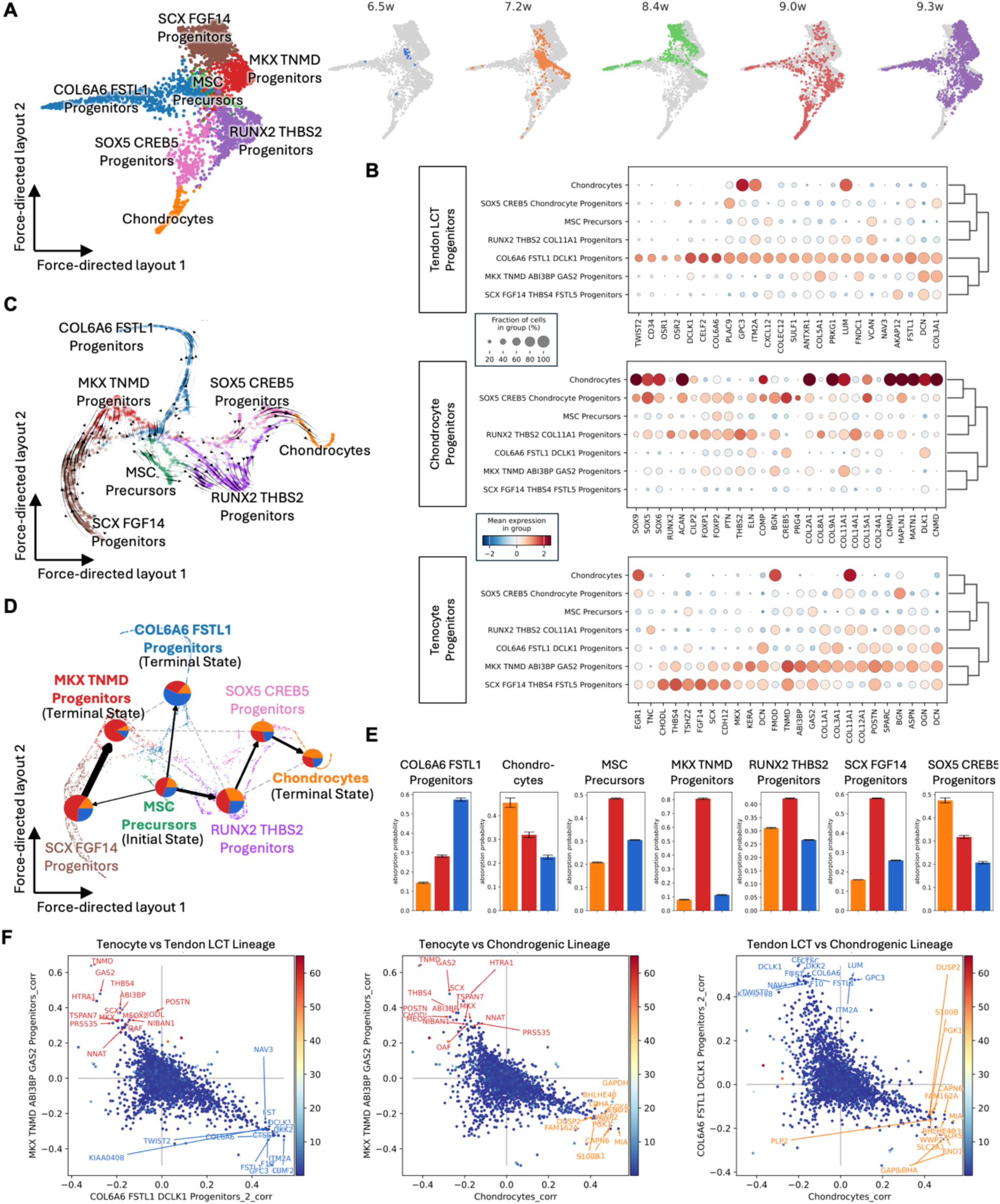
Embryonic progenitor states and lineage trajectories reveal early tendon cell fate specification. (A) Harmony-integrated force-directed layout showing the distribution of embryonic tendon cells annotated by cell type. Right: the same layout coloured by developmental stage (6.5–9.3 post-conception weeks), illustrating temporal dynamics of lineage emergence. (B) Dotplots of scaled, log1p-normalised gene expression showing differentially expressed genes across embryonic tendon cell types. Lineages are grouped by putative fate: fascicular tendon, loose connective tissue (LCT), or chondrocyte. Dot size reflects cell type abundance. (C) RNA velocity analysis overlaid on a force-directed layout derived from t-SNE embeddings calculated using multiscale diffusion components. Arrows indicate the predicted future transcriptional state and directionality of gene expression changes among progenitor populations. (D) CellRank-directed partition-based graph abstraction map showing lineage trajectories from MSC Precursors toward three terminal fates: fibrillar tendon (red), loose connective tissue fibroblast (blue), and chondrocyte (orange). Arrow thickness reflects transition probability; dashed lines indicate weaker transitions. Pie charts show fate likelihoods per cell type, and bar plots in (E) quantify absorption probabilities. (F) Correlation plot of CellRank-derived lineage driver genes. Each axis represents gene correlation with terminal cell fates. Top 15 drivers are annotated for each lineage.

Mesenchymal stem cells (MSCs) were first examined as the likely progenitors of tendon fibroblasts. The earliest state, MSC Precursors, was identified at 6.5-7.2 pcw and lacked classical MSC markers, but expressed a combination of non-myogenic (*HMGA2, FOXP2*, *RUNX1T1*)^78,79^ and myogenic (*SIX1)*^78,80,81^ connective tissue fate regulators. These cells also expressed genes involved in cell structure, adhesion, metabolism, growth, and signalling, alongside low levels of *TNMD*, *TSHZ2*, *GAS2*, and *VCAN* (Figure S4).

By 8.4 pcw, a distinct COL6A6 Progenitor population emerged, expressing canonical MSC markers (*CD73/NT5E*, *CD90/THY1*, *CD44*, *TWIST2*), along with tendon-associated matrisomal genes (*FSTL1*, *DCLK1*, *COL1A1, COL3A1, COL5A1, COL6A1, COL12A1, ELN*, *DCN*, *ASPN*, *OGN*, *FNDC1*, *LUM*, *VCAN*, *EMILIN2*, *MFAP5*, *TNXB*, *POSTN*) (Figure 4B, S5). This population also expressed LCT-associated transcription factors *OSR1* and *OSR2*,^82,83^ suggesting transition toward the *COL6A6-*expressing LCT fibroblast phenotypes observed in second-trimester foetal tendons.

In parallel, three chondrogenic clusters were identified at different stages of differentiation. A multipotent *RUNX2*-expressing population exhibited a hybrid transcriptomic profile, co-expressing collagens linked to tendon (*COL1A1*, *COL3A1*), cartilage (*COL2A1*, *COL9A1*–*COL9A3*), and endothelial basement membranes (*COL8A1*) (Figure 4B). Two more differentiated clusters expressed classic cartilage markers (*SOX5*, *SOX6*, *SOX9*, *ACAN*, *CILP2*, *HAPLN1*, *COMP*, *COL2A1*, *COL9A1*–*COL9A3*) (Figure 4B, S5). The less mature SOX5 Progenitors additionally expressed *CREB5* and *PRG4*, while the more differentiated Chondrocyte cluster was enriched for *MATN1*, *MATN3*, and *MATN4* (Figure 4B).

Finally, two distinct *SCX*-expressing progenitor clusters were identified. SCX Progenitors were characterised by high expression of *FGF14*, *THBS4*, *FSTL5*, *CHODL*, *TSHZ2*, *CDH12*, *KERA*, *POSTN*, *DCN*, *OGN*, and a broad array of fibrillar (*COL1A1*, *COL3A1*, *COL5A1*, *COL6A1*, *COL12A1*, *COL14A1*), basement membrane (*COL18A1*, *COL4A1*, *COL4A2*), and MTJ-associated (*COL22A1*) collagens (Figures 4B, S5). MKX Progenitors exhibited even higher expression of matrisomal genes, along with *LOX*, *MKX*, *TNMD*, *ABI3BP*, and *GAS2*, marking a more differentiated tenogenic state (Figure 4B, S5).

### Trajectory and regulatory network analyses reveal emergence of three distinct tendon-associated lineages during human development

To reconstruct the differentiation pathways underpinning tendon development, we applied RNA velocity^84^ and pseudotime analysis using Palantir^85^ and CellRank^86–88^ across embryonic progenitor populations. To link embryonic trajectories to foetal tendon states, we integrated datasets using scGen^89^ with batch correction by moscot,^90^ enabling continuous reconstruction of lineage emergence from early embryogenesis through the second trimester. Gene regulatory networks were then inferred using SCENIC to identify transcription factors and regulatory programs associated with lineage specification.

RNA velocity analysis indicated that MSC Precursors diverged toward four distinct fates: MKX Progenitors, RUNX2 Progenitors, SOX5 Progenitors, and MSC-like COL6A6 Progenitors (Figure 4C). SCX Progenitors also displayed a bias toward MKX Progenitors, consistent with their early tenogenic identity characterised by high *SCX* but lower *MKX* and *TNMD* expression levels.

To further resolve these differentiation pathways, Palantir pseudotime analysis was applied. Using extreme values of multiscale diffusion components, MSC Precursors were defined as the initiation state, while Chondrocytes (chondrogenic lineage), COL6A6 Progenitors (LCT lineage), and MKX Progenitors (tenocyte lineage) served as terminal states (Figure S6A). Consistent with RNA velocity, Palantir-CellRank analysis mapped a trajectory from MSC Precursors to MKX Progenitors via an intermediate SCX Progenitor stage, recapitulating known tenocyte differentiation hierarchies (Figures 4D, S6B-D). Chondrogenic differentiation proceeded from RUNX2 Progenitors to SOX5 Progenitors, ultimately forming embryonic Chondrocytes. Fate probability visualisation confirmed these transitions across donor ages, while also indicating that most progenitor populations retained some degree of plasticity at later stages (Figure 4D,E).

Analysis of lineage-driving genes revealed strong anti-correlation patterns between competing lineages (Figure 4F). In the tenocyte lineage, genes such as *SCX*, *MKX*, *TNMD*, *ABI3BP*, and *GAS2* were negatively correlated with LCT- and chondrocyte-associated markers, suggesting roles in reinforcing tenogenic identity. Conversely, *TWIST2*, *DCLK1*, and *CELF2* were anti-correlated with tenogenic markers, supporting maintenance of LCT fate. The anti-correlation between LCT and chondrogenic lineages was less pronounced, indicating closer transcriptional relationships between these cell types. Genes like *LUM* and *GPC3* selectively anti-correlated with tenocyte, but not chondrocyte, differentiation, highlighting lineage-specific regulatory mechanisms.

Integration of embryonic and foetal datasets supported these inferred trajectories (Figure 5A). MSC Precursors first differentiated into SCX and COL6A6 Progenitors (6.5-8.4 pcw), followed by RUNX2 and SOX5 Progenitors (8.4-9.0 pcw). By 9.3-12 pcw, SCX Progenitors gave rise to FGF14 THBS4 and ABI3BP GAS2 Fibroblasts, while MKX Progenitors contributed specifically to ABI3BP GAS2 Fibroblasts (Figure 5B). In parallel, COL6A6 Progenitors differentiated into COL3A1 PI16, COL6A6 FNDC1, and NEGR1 SCN7A Fibroblasts, reflecting their broader differentiation potential.

**Figure 5.**
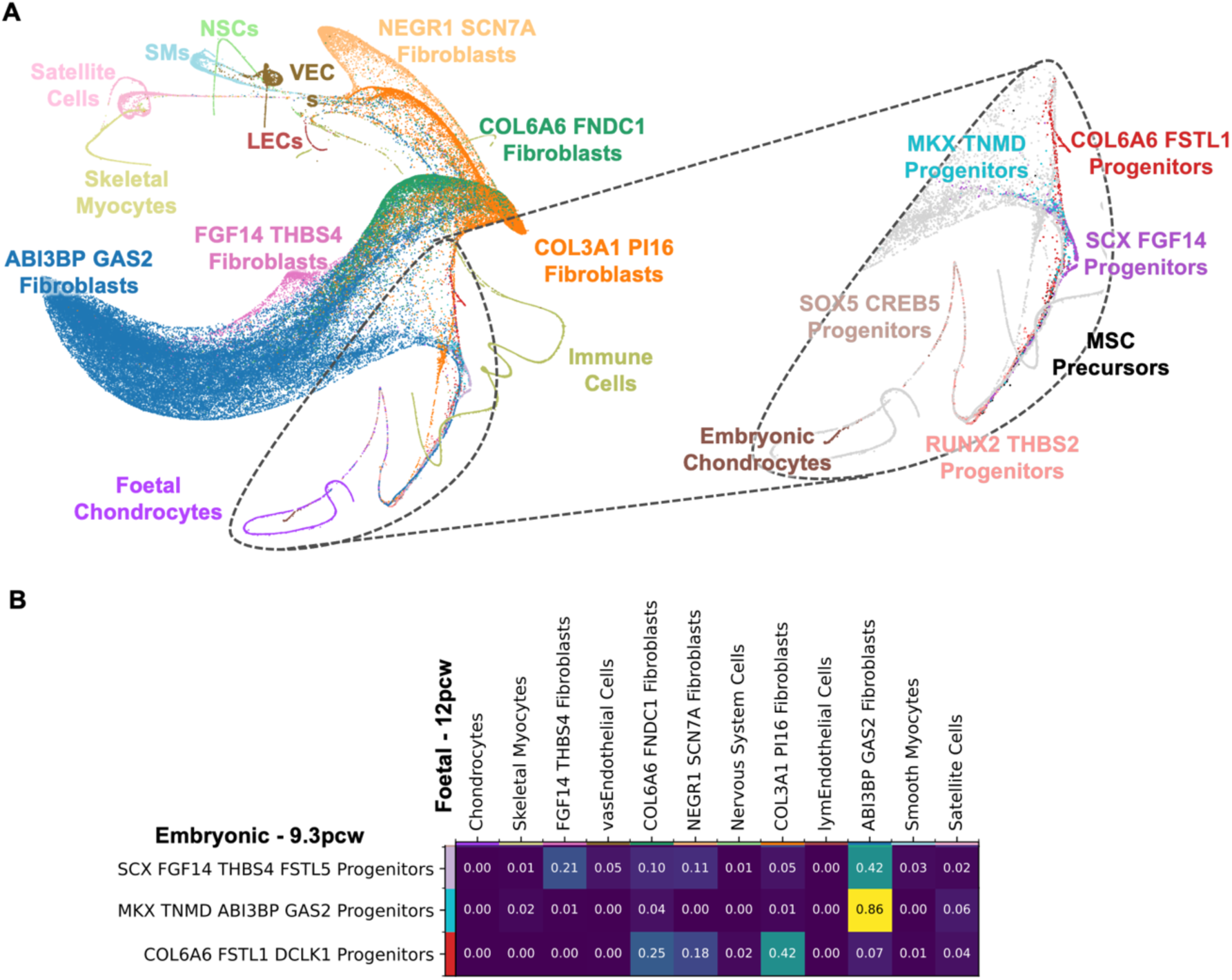
Integration of embryonic and foetal datasets reveals continuity and divergence in fibroblast lineage trajectories. (A) Force-directed layout of scGen-integrated embryonic (6-9 pcw) single-cell RNA-seq and foetal (12-20 pcw) single-nucleus RNA-seq datasets. Inset highlights embryonic tendon cells, which remain largely segregated, reflecting transcriptional divergence likely due to developmental stage differences and platform-specific biases. (B) Transition matrix from moscot analysis showing the inferred descendants of 9.2 pcw fibroblast-lineage progenitor populations within 12 pcw foetal samples. Values represent the proportion of cells predicted to transition into each differentiated fibroblast subtype.

Finally, SCENIC regulatory network analysis of 20 pcw and earlier foetal tendon populations revealed an absence of definitive fibroblast-specific regulons (Figure S7), indicating that tendon fibroblast transcriptional identities remained heterogeneous and incompletely matured at this developmental stage.

### Developing tendons exhibit histological features typically associated with adult tendon pathology

To initiate comparisons between developing and aged tissues, histological and histochemical analyses were performed on foetal and adult Achilles and quadriceps tendons. Foetal tendons exhibited distinct structural characteristics compared to adult tendons, including high cellularity, increased vascularity, and abundant ground substance rich in acidic polysaccharides – features typically associated with tendon pathology in adulthood.^91–93^

Although foetal collagen fibres appeared well-organised, their arrangement was less uniform than in mature tendons. Picrosirius red staining under polarised light revealed progressive collagen maturation: early foetal tendons (11-12 pcw) displayed loosely arranged, thin fibres with low birefringence (consistent with type III collagen), while 16-20 pcw tendons showed thicker fibres with a transition from blue/green to red birefringence, indicative of type I collagen (Figure S8). Collagen crimp patterns also evolved over time, becoming more elongated and approaching the adult tendon architecture by 20 pcw.

Quantitative image analysis confirmed a progressive decline in nuclear density from 11 pcw through adulthood (Figure S9). Nuclear areas decreased between 11 and 15 pcw before returning to 11 pcw baseline levels by 20 pcw. In adult samples, nuclear size was variable, with some nuclei smaller and others comparable to those in late-stage foetal tendons. Nuclear morphology transitioned from elongated to rounded and back to elongated during foetal development, whereas adult tendons exhibited regional variability in nuclear shape, associated more with microanatomical site differences than chronological age (Figure S10).

### Adult tendon fibroblasts diverge transcriptionally from developmental lineages, unlike conserved non-fibroblast populations

Integration of embryonic (6-9 pcw), foetal (12-20 pcw), and adult (25-76 years) Achilles and quadriceps tendon datasets using scANVI showed that developing and adult fibroblast populations remained largely distinct (Figures 6A, S11). However, fibroblasts from ruptured adult quadriceps tendons (referred here as COL3A1hi Fibroblasts, named ADAM12hi Fibroblasts in recent publication^26^) showed higher expression of top differentially expressed genes (DEGs) from foetal ABI3BP GAS2 and COL6A6 FNDC1 Fibroblasts compared to healthy adult tendons, suggesting a shift toward a developmental-like state potentially linked to repair processes (Figures 6B, S12). In contrast, foetal DEGs overlapped significantly with non-fibroblast adult populations, including immune, endothelial, muscle, and nervous system cells (Figure 6A,B).

**Figure 6.**
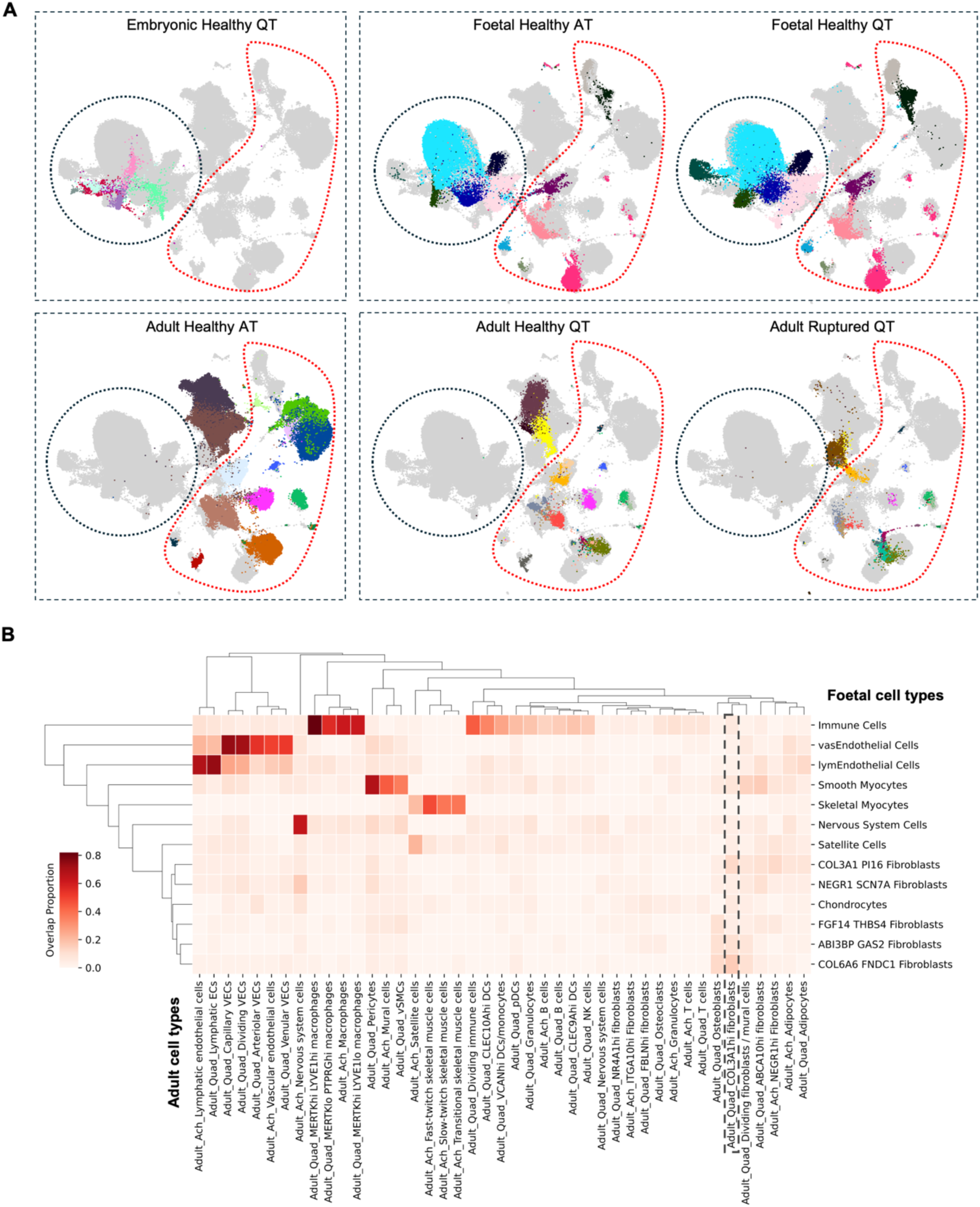
Cross-stage and cross-dataset harmonisation reveals conserved and divergent tendon cell identities. (A) ScANVI-integrated embryonic, foetal, and adult tendon datasets. All embryonic and foetal fibroblast populations are outlined in blue. All non-fibroblast populations are outlined in red. The populations that are not outlined are exclusively adult fibroblasts. Annotated adult datasets were provided by Dr Carla Cohen (Achilles tendon) and Dr Jolet Mimpen (quadriceps tendon); annotations can be found, in full, in Figure S11. (B) Clustermap shows normalised overlap of the top 50 differentially expressed genes in foetal cell types (rows) with those identified in adult cell types (columns), computed using the Wilcoxon rank-sum test. Overlap is expressed as a proportion of the foetal marker gene set, with darker shades indicating higher marker overlap. Adult quadriceps COL3A1hi population (outlined in black) is the unique fibroblast population specific to ruptured adult tendon. Ach/AT: Achilles tendon; Quad/QT: quadriceps tendon.

Cell type harmonisation using CellHint refined annotations across datasets, confirming cross-stage consistency and revealing developmental relationships between foetal and adult tendon populations (Figure S13). For instance, foetal immune cells aligned to adult *MERTK*-expressing Macrophages, and foetal Skeletal Myocytes mapped to Transitional and Fast-twitch Skeletal Muscle Cells in adult Achilles tendons. Adult tendons also exhibited increased cellular diversity, including Adipocytes and NR4A1hi Fibroblasts. While foetal and adult Achilles tendon Nervous System Cells showed continuity, adult quadriceps tendon Nervous System Cells appeared developmentally distinct, lacking a clear foetal counterpart.

This analysis also supported previous trajectory findings. Embryonic MKX Progenitors were linked to foetal ABI3BP GAS2 Fibroblasts; SCX Progenitors to foetal FGF14 THBS4 Fibroblasts; COL6A6 Progenitors to foetal COL3A1 PI16 Fibroblasts; and SOX5 Progenitors, RUNX2 Progenitors, and embryonic Chondrocytes to foetal Chondrocytes (Figure S13). While foetal ABI3BP GAS2 Fibroblasts were traced to adult ITGA10hi and FBLN1hi Fibroblasts in healthy Achilles and quadriceps tendons, respectively, foetal COL3A1 PI16 and NEGR1 SCN7A Fibroblasts were linked to NEGR1hi and ABCA10hi Fibroblasts in adult tendons. Notably, embryonic RUNX2 Progenitors, foetal Chondrocytes, and injury-responsive TSPC marker-expressing COL6A6 FNDC1 Fibroblasts were traced to ruptured quadriceps COL3A1hi Fibroblasts. In contrast, MTJ-associated foetal FGF14 THBS4 Fibroblasts had no clear adult counterpart, indicating a likely transient developmental role.

DGE analysis between foetal and adult quadriceps tendon fibroblasts revealed substantial transcriptional shifts. In adults, 262 pathways were downregulated (Figure 7A), encompassing energy metabolism, nucleotide and protein biosynthesis, apoptosis, cell cycle regulation, telomere maintenance, and stress response processes (Table S3). In contrast, 35 pathways were upregulated (Figure 7A; Table S3), including cellular signalling, structural organisation, cell migration, and nervous system development. Adult fibroblasts significantly upregulated *ECM2* while downregulating genes associated with cartilage (*COL2A1, COL9A1, COL9A3*), early fibrillogenesis (*LUM*), growth factor signalling (*IGFBP2, IGFBP4, IGFBP5*), elastic fibre formation (*EFEMP1, EFEMP2, EMILIN1, FBLN1, MFAP2, MFAP4*), cell adhesion (T*GFBI, VWA1*), collagen deposition (*CTHRC1, PCOLCE*), and calcification inhibition (*MGP*) (Figures S14-17). These findings suggest a transition from the highly metabolic and biosynthetically active foetal state to a more maintenance-oriented state adult fibroblast phenotype.

**Figure 7.**
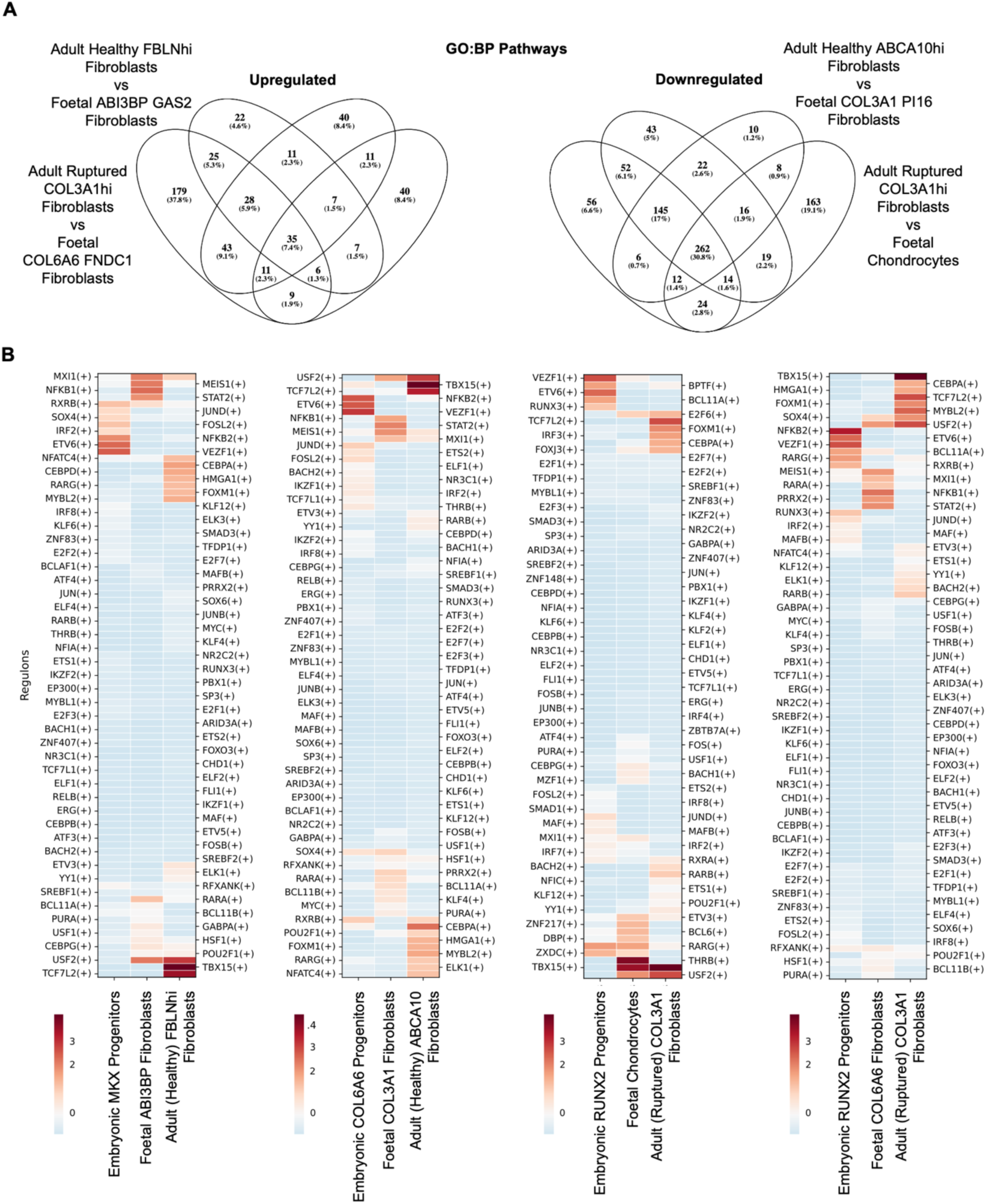
Conserved and divergent pathways and gene regulatory programs in fibroblasts across tendon development and ageing. (A) Venn diagrams of intersecting upregulated or downregulated gene ontology biological process (GO:BP) pathways in CellHint-paired foetal and adult cell types. Only significant (<0.01 BH-FDR) pathways with term sizes of 20-500 were used. (B) Hierarchical clustermaps showing z-score normalised SCENIC regulon activity in selected fibroblast subtypes from embryonic, foetal, and adult tendon datasets. Regulons are clustered using average linkage and Euclidean distance; cell types remain in original order. Colour scale indicates relative regulon activity (blue: below mean, white: mean, red: above mean). Only regulons shared across all datasets are shown. Regulon names are alternately labelled on the left and right axes for clarity.

Finally, SCENIC analysis of aged human tendons revealed strong cell type-specific regulons, with limited regulon conservation across development and adulthood. Notably, the RARG regulon remained active in embryonic RUNX2 Progenitors, foetal Chondrocytes, and adult ruptured tendon COL3A1hi Fibroblasts (Figure 7B), regulating 19 genes (e.g., *ARHGEF10L*, *PIK3R1*, *BOC*, *PCDH9*, *ZFHX3*, *PDGFD*, *DCLK1*) linked to epithelial-to-mesenchymal transition (EMT) and wound healing.^94–97^ The EMT regulators controlled by SCX – *TWIST1* and *SNAI1*^98^ *–* were highly activated in ruptured COL3A1hi Fibroblasts compared to healthy fibroblasts. While TWIST1 regulon was associated with genes regulating ECM remodelling and fibroblast function (e.g., *BCL7A, BICC1, CACNB2, COL1A2, HMCN1, HTRA1, PLEKHA5*), SNAI1 regulon was linked to genes involved in angiogenesis, cytoskeletal organisation, and fibrosis (e.g., *COL4A1, EFNB2, KDR, MIR31HG, PALLD, SLC45A4*). Additionally, both ruptured and healthy adult fibroblasts exhibited high activation of the TBX15 regulon, which encompassed 615 associated genes linked to growth, differentiation, ECM organisation, and tissue-specific developmental pathways, including Wnt and Notch signalling.

## DISCUSSION

Tendon development and homeostasis rely on complex transcriptional programs that guide progenitor differentiation, ECM organisation, and tissue specialisation. By integrating single-cell, single-nucleus, and spatial transcriptomic datasets spanning embryonic, foetal, and adult human tendons, we mapped the emergence and diversification of fibroblast populations across developmental stages and following tendon injury. Our findings reveal substantial transcriptional and functional reprogramming of tendon fibroblasts from early development to adulthood, with the preservation of certain regulatory features – particularly in the context of repair.

### Differentiation trajectories and lineage commitments in embryonic and foetal tendon development

We show that during early development (6.5-8.4 pcw), multipotent MSC Precursors diverge into distinct tendon, LCT, and chondrocyte lineages. *SCX*-expressing tendon progenitors transition into MKX Progenitors, committing to fibrillar ECM production and forming the core tendon fibroblast lineage. In parallel, MSC-like COL6A6 Progenitors give rise to LCT fibroblasts, while RUNX2 and SOX5 Progenitors mark early chondrogenic differentiation, progressing toward mature embryonic Chondrocytes through coordinated activity of *SOX5*, *SOX6*, *SOX9*, and *RUNX3*.

By the second trimester (12-20 pcw), tendon fibroblasts diversify into spatially and transcriptionally distinct subpopulations. Within the fibrillar core, ABI3BP GAS2 Fibroblasts and more differentiated KERA TNMD Fibroblasts contribute to matrix organisation, remodelling, and tenocyte differentiation, with the latter population becoming more prevalent over time. A small CHSY3 MECOM Fibroblast population may contribute to ECM hydration, mineralisation, and tendon–bone interface remodelling. Within the LCT, CREB5 PRG4 Fibroblasts likely facilitate lubrication and ECM maintenance under mechanical load, COL3A1 PI16 Fibroblasts support tissue remodelling and matrix adaptation through cell–ECM interactions, and COL6A6 FNDC1 Fibroblasts show early injury-responsive and fibrillogenesis-associated features, suggesting a developmental link to adult repair processes. At the tendon– muscle junction, FGF14 THBS4 Fibroblasts contribute to ECM synthesis, vascularisation, and immune interactions, but decline in abundance by 20 pcw, indicating a transient role in MTJ development. Similarly, NEGR1 SCN7A Fibroblasts – localised to the muscle endo- and perimysium – appear to support tendon–muscle integration but diminish with maturation.

A key insight from this study is the identification of a previously uncharacterised *SCX*-negative progenitor population expressing *COL6A6* and *FSTL1*, which branches from MSC Precursors into both LCT and muscle-associated fibroblasts. These progenitors are defined by regulatory programs involving *TWIST2*, *OSR1*, *OSR2*, *RUNX1T1*, *DCLK1*, and *CELF2*, and exhibit strong anti-correlation with the SCX–MKX–TNMD– ABI3BP–GAS2 tenogenic program. Their foetal descendants express injury-responsive TSPC markers *TPPP3* and *PDGFRA*, implicating this axis as a potential contributor to both tendon development and regeneration.^13,34^

While our data confirm the presence of *SCX*-expressing progenitors from early embryogenesis through the second trimester, further work is needed to clarify the relationship between the *SCX*-negative COL6A6 lineage at 8.4 pcw and earlier *SCX*-expressing FGF14 Progenitors identified at 7.2 pcw. Existing studies have primarily focused on *SCX*-expressing lineages, with limited exploration of LCT origins. This gap likely reflects both a historical lack of definitive LCT markers and technical challenges in imaging early-stage tendons, where *SCX*-negative domains may be difficult to resolve. For example, murine studies using Scx-GFP reporters suggest uniform Scx expression across tendons until late foetal development,^99,100^ potentially overlooking early LCT heterogeneity.

### Fibroblast plasticity contrasts with conserved non-fibroblast programs across tendon development and ageing

Beyond fibroblasts, several non-fibroblast populations play key roles in tendon development and homeostasis. *PECAM1*+ vascular endothelial cells, *PROX1*+ lymphatic endothelial cells, and *ACTA2*+ smooth myocytes are present during the development. Cells expressing neural markers such as *NRXN1*, *NCAM2*, and *SOX10* suggest a role in neural development and tendon innervation. Furthermore, *PAX7*-expressing satellite cells and myocytes demonstrate gene expression patterns linked to muscle differentiation and function, underscoring the interconnectedness of tendon and muscle development. Immune cells, predominantly macrophages expressing *F13A1*, *MRC1*, and *CD163*, are also abundant, consistent with their known roles in tissue patterning and immune regulation.^40,101–104^

Notably, these populations persist into adult tendons, maintaining conserved transcriptional programs across developmental stages. This suggests they serve fundamental, stable roles in tendon physiology, extending beyond development to support tissue maintenance, innervation, and vascularisation across the lifespan. In contrast, fibroblasts exhibit significant age- and health-specific transcriptional divergence. Dimensional reduction embeddings show clear separation between developmental and adult fibroblast populations, reflecting adaptive transcriptional programs tuned to tissue state and biomechanical demands. While certain populations – such as ABI3BP GAS2 Fibroblasts – persist across time and align with adult ITGA10hi and FBLN1hi subsets, others, like FGF14 THBS4 Fibroblasts at the MTJ, appear transient and lack clear adult counterparts. Furthermore, some cell populations, including adipocytes and neural cell types, are exclusive to adult tendons.

A particularly striking observation is the link between adult rupture-specific COL3A1hi Fibroblasts and embryonic RUNX2 Progenitors, foetal LCT COL6A6 FNDC1 Fibroblasts, and Chondrocytes. These adult fibroblasts express genes typically active during foetal development, suggesting a partial reactivation of foetal programs following injury – a mechanism potentially beneficial for initiating repair.^105–107^ However, these reactivated cells lack TSPC markers and do not recapitulate full regenerative programs observed in foetal tendons.^6,8^ Their precise origin – whether reprogrammed fascicular or LCT fibroblasts – remains unclear.

As tendons age, the synthetic and metabolic activity of fibroblasts declines, correlating with increased matrix stiffness and reduced elasticity.^108–113^ Our DGE analysis supports this shift, revealing a global transition from biosynthetic and proliferative programs to maintenance-focused gene expression in adult fibroblasts. Upregulated pathways include cell signalling, structural regulation, and nervous system development, while genes involved in cartilage and elastic fibre formation are downregulated. Notably, adult fibroblasts also exhibit reduced expression of proteoglycans and early fibrillogenic collagens, aligning with diminished ECM turnover after tendon maturation.^114–117^

Regulatory network analysis further underscores this divergence. Few regulons are shared between developmental and adult tendon fibroblasts, highlighting transcriptional reprogramming across maturation. For example, adult rupture-specific fibroblasts activate EMT-related regulons associated with migratory and invasive phenotypes commonly seen in wound healing and fibrosis,^118^ suggesting that this process may be involved in adult tendon injury repair.

These findings diverge from murine studies of tendon repair. Our human quadriceps tendon samples were collected 8–9 days post-rupture, corresponding to the fibroblastic proliferative phase characterised by collagen III-rich granulation tissue, prior to collagen I-mediated remodelling.^14,119–121^ In contrast, mouse models at this stage show active infiltration of migrating LCT cells expressing *Acta2*, *Sca1*, *Axin2*, or *Glast.*^9,12,35,36,39,42,43^ We did not observe expression of these markers in rupture-specific COL3A1hi or other adult fibroblasts in our dataset. A likely factor is the advanced age of the human donors (67–75 years), in contrast to young adult mice (2.5–6 months) used in tendon injury experimental models.^9,12,35,36,39,42,43^ To bridge this gap, future work should directly compare human tendon populations across developmental and post-injury stages with murine and zebrafish models, enabling more accurate mapping of tendon repair processes across ages and species.

Finally, research to date suggests that tendon healing is most effective before significant mechanical loading and fibril expansion occur, with regenerated tendons typically displaying smaller, more immature collagen fibrils compared to uninjured tendons.^5,9^ In this study, ruptured adult COL3A1hi Fibroblasts downregulated key genes involved in elastic fibre production (*EMILIN1*, *FBLN1*, *MFAP2*) and collagen fibril initiation, deposition, and processing (*COL11A1*, *CTHRC1*, *PCOLCE*, *MGP*), potentially impairing structural repair. Alongside reduced metabolic and biosynthetic activity, these observations align with previous findings that foetal tendon fibroblasts are more proliferative and metabolically active,^38,122^ and that ECM turnover in adult tendons is minimal after late adolescence.^116,117^ Together, our findings suggest that adult tendon fibroblasts primarily maintain tissue homeostasis rather than help regenerate it, a limitation that may underlie the poor repair outcomes observed clinically.

In summary, by integrating single-nucleus and spatial RNA-sequencing approaches, we mapped the cellular and transcriptional architecture of developing and adult human tendons. We defined distinct cell types within foetal Achilles and quadriceps tendons and reconstructed the developmental trajectories of seven embryonic progenitors contributing to the formation of fibrillar core fibroblasts, loose connective tissue fibroblasts, and chondrocytes. Comparative analyses between embryonic, foetal, and aged adult tendons revealed profound shifts in cell composition, density, and ECM organisation: while vascular, immune, neural, and muscle-associated non-fibroblast populations remained transcriptionally conserved across the lifespan, fibroblasts showed striking age- and health-specific divergence, transitioning from metabolically active, matrix-synthesising phenotypes in development to homeostatic maintenance states in maturity. Notably, rupture-associated adult fibroblasts partially reactivated developmental gene programs but failed to express canonical TSPC markers or fully recapitulate regenerative trajectories, highlighting limitations in the adult tendon’s capacity for repair.

These findings provide a comprehensive atlas of tendon cell diversity across the human lifespan. They underscore the need to characterise fibroblast plasticity and immune–stromal interactions in greater detail and point to specific cellular targets and regulatory pathways that may be leveraged to enhance repair. Future work translating these insights into therapeutic strategies – particularly those that emulate or reintroduce regenerative developmental programs – could help address the persistent clinical challenges of tendon injury, degeneration, and ageing.

## RESOURCE AVAILABILITY

### Lead Contact

Further information and requests for resources and reagents should be directed to and will be fulfilled by the Lead Contact, Sarah Snelling.

### Materials Availability

This study did not generate new unique reagents.

### Data and Code Availability

Processed and annotated foetal tendon single-nucleus RNA-sequencing data is available through CellxGene (https://cellxgene.cziscience.com/collections/7b9ae565-a781-433d-98d4-430394e7802a). Raw data is stored within an AWS bucket and is available upon request from the lead contact and the CZI Lattice. Foetal tendon spatial RNA sequencing data is available upon request from the lead contact. All code used for the analysis is available at https://github.com/AlinaKurjan/DPhilCode.

## ACKNOWLEDGMENTS

We are grateful to the staff of the Human Developmental Biology Resource (UCL Institute of Child Health, London) for their efforts in obtaining the human foetal material, as well as the donors that have consented for the use of the material. In addition, we thank the staff at the Histology Service (Kennedy Institute of Rheumatology, Oxford) for their help with histology and histochemistry. We also thank Dr Peng He, Dr John E. G. Lawrence, and Prof Sarah A. Teichmann for processing the embryonic single-cell and spatial RNA-sequencing data of whole embryonic limbs to meet the requirements of this work and for aid with analysis and interpretation of results; Dr. Carla Cohen for providing annotated human Achilles tendon data for comparisons, Dr. Claudia Paul for providing stained sections of adult supraspinatus tendons for comparisons, Dr Mate Naszai for assistance with quantitative image analysis, Chinemerem Ikwuanusi for assistance with tissue processing and immunofluorescence stainings, and Naomi Gray for assistance with logistics.

This work was supported by funding from the Oxford-Medical Research Council Doctoral Training Partnership, https://www.medsci.ox.ac.uk/study/graduateschool/mrcdtp/; the National Institute for Health Research (NIHR), https://www.nihr.ac.uk/ and the NIHR Oxford Biomedical Research Centre (BRC), https://oxfordhealthbrc.nihr.ac.uk/ (NIHR203311; M.J.B. and S.J.B.S.); the Chan Zuckerberg Initiative (2019-002426; J.Y.M., L.R.M., M.J.B., A.P.C., S.J.B.S.), https://chanzuckerberg.com/. The views expressed are those of the authors and not necessarily those of the NHS, the NIHR or the Department of Health. The funders had no role in study design, data collection and analysis, decision to publish, or preparation of the manuscript.

## Author Contributions

S.J.B.S., M. J. B., C.D.B., A.P.C., and A.K. conceived and designed the study. A.K., M. J. B., J.Y.M., L.R.M and A.C.A. collected samples. A.K., J.Y.M., L.R.M and A.C.A carried out the experiments. A.K. performed the bioinformatics analyses, analysed, and interpreted data. J.Y.M. provided processed and annotated data for adult quadriceps tendons. A.K., S.J.B.S., and M. J. B. wrote the manuscript. All authors provided critical feedback and approved the final version of the manuscript.

## STAR METHODS

### EXPERIMENTAL MODEL AND STUDY PARTICIPANT DETAILS

#### Human foetal tissue samples

Human foetal material was supplied by the Joint MRC/Wellcome Trust (grant #MR/R006237/1) Human Developmental Biology Resource (HDBR, www.hdbr.org) under the MTA license (R60786/CN008) with REC 18/LO/0822. Fresh foetal lower limbs aged 9-20 post-conception weeks (pcw) were provided by the HDBR London UCL Institute of Child Health. Foetal age was estimated using the independent measurement of the crown rump length (CRL), using the formula PCW (days) = 0.9022 3 CRL (mm) + 27.372 and then rounding up to the full decimal. All samples used in this work were sourced from elective terminations with no abnormalities recorded. Foetal sample ages in post-conception weeks were as follows: DEV16134 – 12, DEV16135 – 12, DEV16171 – 12, DEV16136 – 12, DEV16127 – 17, DEV16569 – 17, DEV15983 – 20, DEV15984 – 20, DEV15985 – 20, DEV16126 – 20. DEV16126 was used for Visium spatial RNA-sequencing, while the remaining samples were used for single-nucleus RNA-sequencing. Sample metadata is summarised in Table S1.

#### Human adult tissue samples

Ethical approval for the Oxford Musculoskeletal Biobank (OMB, 19/SC/0134) was granted by the Oxford Research Ethics Committee B for all work on human Achilles and quadriceps tendons. Written informed consent according to the Declaration of Helsinki was obtained from all patients. Healthy Achilles and quadriceps tendon tissues were collected from patients undergoing above- or below-the-knee amputations (e.g., OMB0785) or suprapatellar nailing of tibial shaft fracture (e.g., OMB1266). Ruptured quadriceps tendon samples were obtained from patients with acute full quadriceps tendon ruptures, with surgeries performed 8 to 9 days post-rupture. Patient ages in years were as follows: healthy Achilles tendon (OMB0785 – 74, OMB1556 – 51, OMB1250 – 45, OMB1691 – 58, OMB1687 – 76); healthy quadriceps tendon (OMB0792 – 29, OMB1266 – 25, OMB1248 – 44); ruptured quadriceps tendon (OMB0778 – 67, OMB0793 – 69, OMB0779 – 75). Sample metadata is summarised in Table S1.

### METHOD DETAILS

#### Foetal and adult tendon tissue processing

The tissues were washed in PBS (phosphate-buffered saline) and dissected to retain the regions of interest – whole tendons for foetal tissues, including muscle and bone attachment sites; enthesis, midbody, and MTJ regions (∼1cm pieces) for adult tissues – using anatomical landmarks. Adult tissue cuts were photographed to retain topographical reference.

The tissues were then either formalin-fixed for staining or snap-frozen in liquid nitrogen for sequencing as soon as possible after collection. Following formalin-fixation (with different lengths of time depending on the size of the samples), the samples were resuspended in 70% IMS (industrial methylated spirit) and stored at room temperature. After dehydration, formalin-fixed tissues were paraffin-embedded and sectioned. Snap-frozen tissues were stored at -70°C to -80°C before being used for single-nuclei or spatial RNA-sequencing.

#### Foetal and adult tendon nuclei isolation

Nuclei isolation for foetal and adult tendon samples outlined in Table S1 was carried out following protocols for large (adult) and small (foetal) tissues developed and published by Mimpen and colleagues from the Tendon Seed Network.^123^ Briefly, tendons were cut into ∼1 mm pieces on dry ice and dissociated in CST buffer (292 mM NaCl, 20 mM Tris-HCL 7.5 pH, 2 mM CaCl2, 42 mM MgCl2, 0.5% CHAPS, 0.01% BSA, RNase and protease inhibitors) on a rotor for 2 or 10 min at 4°C. After addition of PBS with 2% or 1% BSA (bovine serum albumin) the suspensions were strained through 20 or 40 µm strainers (Greiner Bio-one), with former numbers used for foetal and latter for adult tissues. The nuclei-containing suspensions were then centrifuged at 500 g for 5 min at 4°C. Following centrifugation, the supernatant was discarded and the nuclei within the pellet were stained with DAPI and counted manually using a haemocytometer (NanoEntek DHC-N01) and fluorescence microscopy.

#### Foetal and adult tendon single-nucleus RNA-sequencing

Nuclear suspensions, diluted in PBS with 1% BSA to a concentration of 200-1000 nuclei/μl, were loaded onto a Chromium Next GEM Chip G (10x Genomics) with the aim of recovering 1,000-10,000 nuclei per sample. The samples were then processed using the Chromium Controller (10x Genomics) and prepared into libraries using the Chromium Next GEM Single Cell 3’ Reagent Kits v3.1 (10x Genomics) following the manufacturer’s instructions. Libraries were indexed with the Single Index Kit T Set A (10x Genomics). Quality control assessments for cDNA and final libraries were conducted using D1000 or D5000 High Sensitivity ScreenTape (Agilent) assays on a 4150 TapeStation System (Agilent). The final libraries were pooled together and sequenced using a NovaSeq 6000 (Illumina) by Genewiz (UK) with a minimum sequencing depth of approximately 20,000 read pairs per expected nucleus.

#### Adult tendon snRNA-seq data processing

Human adult tendon single-nuclei RNA-sequencing datasets (*N*=26 from 12 donors) were processed, integrated and annotated by Dr Carla Cohen (Achilles tendon) and Dr Jolet Y. Mimpen (quadriceps tendon; published in ^26^) within the framework of the Tendon Seed Network^124,125^ (Oxford, UK). Briefly, raw sequencing files were processed using scflow^126^ (custom development version; pipeline *scflow quantnuclei*), with reads mapped to the human Ensembl GRCh38 transcriptome (release 106) using kallisto bustools^127^ (v0.27.3). Single-nucleus RNA-seq analysis and annotation was performed in R (v4.3.1) and RStudio Server (v2023.03.1, build 446) using SingleCellExperiment^128^ (v1.22.0) and Seurat^129^ (v4.3.0.1) packages. The counts were log-normalised using Seurat’s default functions. QC metrics were calculated with scuttle^130^ (v1.10.1). Filtering thresholds for number of cells, number of features and mitochondrial ratios were set manually for each sample to remove poor-quality cells. Doublets were detected and removed with scDblFinder^131^ (v1.12.0) using default settings. Ambient RNA was detected using *decontX()* from celda^132^ (v1.14.0). Further ambient RNA detection was performed using SoupX^133^ (v1.6.2), and the soupX-adjusted count matrix was used for downstream analysis. Integration of samples was performed using Harmony^134^ (v0.1), with a combined sample donor and tissue type variable specified for batch correction. Clusters were defined with Seurat’s *FindClusters()*, and a cluster comprising nuclei with high decontX scores was removed. Annotation was performed by assessing cluster-specific expression of manually curated gene sets.

#### Foetal tendon snRNA-seq data processing

Foetal tendon raw sequencing files were aligned with CellRanger^135^ (v7.0) using 10x Genomics’ pre-built GRCh38-2020-A reference (compiled from ENSEMBL’s 98th release of the human reference genome) with default settings. Ambient RNA was removed using CellBender^136^ (v0.2.2) with custom values for expected-cells, total-droplets-included, epochs and low-count-threshold parameters, which were selected based on the properties of individual samples and what was expected. The command was rerun until optimal conditions (training and test loss converging, expected number of cells selected, no error or warning messages in log files) were achieved for each sample. To obtain spliced and unspliced count matrices, velocyto^84^ (v0.17.17) command line function *velocyto run10x* was used with the aforementioned reference genome file and a repeat sequences masked gtf file (h38_repeat_mask.gtf) downloaded from UCSC Genome Browser. The resulting loom output files were compared with CellBender output files, then merged using scVelo^137^ (v0.2.5) function *scvelo.utils.merge()* to retain overlapping barcodes.

Data processing and analysis were carried out using Python and R packages. Briefly, merged files were filtered to remove: 1) genes that were detected in fewer than 20 cells (using Scanpy^138^ (v1.7.2) function *scanpy.pp.filter_genes(adata, min_cell=20))*, 2) genes with 0 UMI counts, and 3) cells with fewer than 200 UMI counts. Following this basic filtering, low quality reads were removed using permissive automatic thresholding based on median absolute deviations (MAD), as described in single-cell best practices guidelines.^139^ Cells were marked as outliers and filtered out if they differed by 5 MADs in their logarithmical total counts, genes-by-counts, and percentage counts in top 20 genes. Additionally, cells with more than 10% mitochondrial counts and 3 MADs were also removed. Doublets were removed individually for each sample using scDblFinder^131^ (v1.4.0) with default settings. Finally, QC plots were analysed to determine the necessity of additional filtering thresholds for each individual sample. The cells were then filtered by manually defined minimal genes-by-counts thresholds that ranged from 200 to 500.

For normalisation, the filtered and concatenated anndata object was split into separate adult-only and developmental-only sample objects. Each was normalised using a shifted logarithm approach based on the delta method (referred to as log1pPF) involving *scanpy.pp.normalize_total(target_sum=None)*followed by *scanpy.pp.log1p()*, as recommended by the recent comprehensive data transformation benchmark study.^140^ Top 3,500 highly variable genes were selected for adult-only and developmental-only objects using scIB^141^ (v1.1.3) package’s *scib.preprocessing.hvg_batch()* function, with *flavor* and *batch*_*key* parameters set to ’cell_ranger’ and ’sampletype’, respectively. Additional filtering was done to remove 51 genes detected in less than 5 counts and remove counts with fewer than 200 genes expressed. Cell cycle phase was determined using *scib.preprocessing.score_cell_cycle()*. The data were then scaled using a custom function *split_and_scale()* that split the concatenated objects by ‘sampletype’ and applied *scanpy.pp.scale()* individually for each sample. Dimensionality reduction in the form of principal component analysis (PCA) was then applied to scaled highly variable genes, and neighbours (*n_neighbors=30, npcs=15*) were calculated to produce uniform manifold approximation (UMAP) plots to assess the quality control steps.

Foetal sample data were integrated and batch-corrected using scvi-tools^142^ (v0.16.1) package’s single-cell variational inference (scVI)^46^ modelling on all unnormalised gene counts, with the main batch effects of interest specified to be ’sampletype’ and ’libbatch’, corresponding to different donor and tissue type (e.g., DonorID1_AchillesTendon and DonorID1_QuadTendon) and library preparation batches, respectively. Model hyperparameters were optimised using manual runs with different parameters as well as using scVI’s autotune functionality, which determined the best fitting parameters to be ’n_latent’: 30, ’n_layers’: 2, ’dropout_rate’: 0.1, ’gene_likelihood’: ’zinb’, ’dispersion’: ’gene-batch’. The model was trained using 398 epochs until model training and validation sets were stably converged.

Data clustering was carried out using Scanpy’s Leiden algorithm, identifying a total of 19 clusters (0-18) at 0.6 resolution. Normalised and log-transformed cluster gene counts were ranked using a Wilcoxon rank-sum test with *scanpy.tl.rank_genes_groups().* The clusters were manually annotated by checking known cell type markers and by querying top 350-550 Leiden cluster DEGs with CellMESH^143^ and gProfiler’s g:GOSt functional profiling^144^ tools. Heatmaps and hierarchical clustering dendrograms were also consulted in the process to make the best guesses for the previously undefined cell types. Following scVI integration and cell type labelling, scANVI^145^ was run with the scVI model as basis for 25 epochs. The resulting latent representation embeddings were used for the computation of a neighbourhood graph of observations, producing annotated data UMAPs.

#### 10X Visium spatial RNA-sequencing of foetal samples

Foetal Achilles (*N=1*) and quadriceps tendons (*N=2*) (Table S1) were dissected from both legs of a single 20 pcw foetus and flash frozen in liquid nitrogen. In preparation for spatial transcriptomics (ST), the samples were cut to ≤0.65 cm² to fit the 10x Genomics Visium ST slide regions. We were able to retain enthesis-to-MTJ as well as adjacent muscle tissue regions for both types of tendons. The samples were embedded in cold OCT mounting medium (VWR) on dry ice and cut longitudinally into 10 μm sections, which were then fixed and stained with H&E to verify tissue morphology and suitability. The sections were prepared for sequencing according to the 10X Genomics recommended protocols using the Visium Gene Expression Slide and Reagent Kit alongside a Dual Index Kit TT Set A. Libraries were sequenced using Illumina NextSeq500 (paired-end) at a depth of 54,000 (Quad2 tendon), 74,000 (Ach) and 119,000 (Quad1) mean reads per spot. The data and images were processed with SpaceRanger (v1.3.1; 10X Genomics) using default settings and mapped to the GRCh38 reference genome.

#### Foetal spatial RNA-seq data processing

The data were processed using Scanpy^138^ (v1.9.5) and Squidpy^146^ (v1.2.3). Briefly, tissue objects were manually filtered to remove: 1) cells with fewer than 500-1,000 counts or more than 10,000-20,000 counts (with exact numbers depending on individual sample properties), 2) genes that were detected in fewer than 10 cells, and 3) any ribosomal and mitochondrial reads. The counts were then normalised to log1pPF, and 2,000 highly variable genes were selected with the “cell_ranger” flavor using *scanpy.pp.highly_variable_genes()*. After normalised count scaling with *scanpy.pp.scale()*, PCA was carried out, and neighbours and UMAPs were calculated for the data using default Scanpy functions. Finally, all tissue objects were subjected to Leiden clustering at 1.0 resolution, identifying 7 clusters for each of the samples. The clusters were examined by analysing the outputs of a Wilcoxon rank-sum test run using *scanpy.tl.rank_genes_groups()*

#### Foetal cell type mapping to spatial coordinates

To infer the spatial distribution of cell types within the tissue, processed and annotated foetal snRNA-seq data were integrated with spatial RNA-seq information using cell2location^47^ (v0.1.4). Cell2location hyperparameters were specified based on the tissue and experiment considerations in mind as having a) expected cell abundance per Visium spot set to 17 (average from a range of 10-32 nuclei in different regions), and b) regularisation of within-experiment variation in RNA detection sensitivity set to 20. The model was trained until convergence for a total of 16,000 iterations.

H&E images were used as basis for microanatomical tissue region identification and labelling. To further analyse potential tissue microenvironments in an unsupervised manner, non-negative matrix factorization (NMF) using cell2location’s scikit-learn NMF wrapper function was also applied to estimated cell abundances. The model was trained using a range of factors (5-30) for decomposition of cell abundance data, with smaller factors assuming lower numbers of distinct patterns and higher likelihood of cell co-location.

#### Immunofluorescence staining and imaging of foetal tendons

Snap-frozen tendon samples were embedded in OCT (VWR) and sectioned at 7 μm thickness. All staining procedures followed the Cell DIVE Platform protocol (GE Research, Niskayuna, NY, USA). Tissue sections were post-fixed for 1 minute at 4 °C in a 1:1 ethanol-acetone solution, then blocked overnight at 4 °C in PBS containing 3% BSA and 10% donkey serum (Bio-Rad).

Slides were stained with DAPI (ThermoFisher) and mounted using an antifade medium containing 4% propyl gallate and 50% glycerol (Sigma-Aldrich). Initial imaging at 20X magnification was performed to capture background autofluorescence, which was subtracted from subsequent staining rounds. Following this, coverslips were removed in PBS, and slides were incubated with antibodies overnight at 4 °C. After incubation, slides were washed three times in PBS (5 minutes each with gentle agitation), re-coverslipped, and imaged.

A bleaching step was then performed by decoverslipping and incubating the slides three times for 15 minutes in 0.5 M NaHCO₃ (pH 11.2) containing 3% H₂O₂, with 1-minute PBS washes between each bleach. This was followed by three additional PBS washes and a 2-minute DAPI recharge. Slides were then re-coverslipped, and a bleached image was acquired for subtraction from the next staining round. Image analysis was conducted using QuPath (v0.5.1).

#### Histochemistry of foetal and adult tendon tissues

Histochemical staining of foetal and adult sectioned formalin-fixed tissues was performed by the Histology Team at the Kennedy Institute of Rheumatology. The slides were stained with haematoxylin and eosin (H&E), alcian blue, masson’s trichrome, and picrosirius red. Slides were scanned with Motic EasyScan One. PSR-stained slides were additionally imaged under polarized light using an Olympus BX40.

#### Embryonic whole limb data processing

Single-cell RNA-sequencing (*N*=25 from different hind limb regions of 11 donors) and 10X Visium spatial RNA-sequencing (*N*=8 from different hind limb regions of two 6pcw and one 8pcw donors) files were provided by the Teichmann group at the Wellcome Sanger Institute (Cambridge, UK) (Table S1). The tissues were collected, processed and sequenced by the group in accordance with the methods published by Zhang et al.^77^. For this analysis, the raw sequencing files were processed and aligned in the same way as was done for the foetal samples to minimise bias, using same versions of the CellRanger and CellBender packages for the single-cell data and the SpaceRanger package for the spatial data.

Human embryonic spatial RNA-seq data were analysed using Loupe Browser (v6.4.1; 10X Genomics). Expression of early tenocyte markers (e.g., *SCX*, *MKX*, *FMOD*, *TNMD*, *EGR1* etc.) and H&E-stained sections guided tendon tissue annotation. While tendon regions could not be confidently defined in 6 pcw samples, a merged 8 pcw hindlimb sample enabled identification of developing patellar and quadriceps tendon areas (Figure S2). Of 2,439 Visium spots, 28 were annotated as tendon: 18 patellar and 10 quadriceps (Figure S2).

Single-cell embryonic data samples from 6 to 9 pcw (*N*=25, total of 108,617 cells) were processed and integrated using scvi-tools’ scVI modelling on all unnormalised gene counts, with the main batch effects of interest specified to be ‘samplename’ (consisting of Sample ID), ‘kit’, ‘seq_protocol’, and ‘sex’ (Figure S3A). As before, model hyperparameters were optimised with scVI autotune and manual runs to ‘n_hidden’: 256, ’n_latent’: 14, ’n_layers’: 3, ’dropout_rate’: 0.1, ’gene_likelihood’: ’nb’, and ‘dispersion’: ‘gene-batch’. The model was trained using 394 epochs until training and validation sets were stably converged. The resulting latent representation embeddings were then used as basis for the computation of a neighbourhood graph of observations and dimensionality reduction with UMAP.

To identify tendon cell subsets in embryonic whole-limb scRNA-seq data, a random forest classifier was trained using spatial transcriptomics as a reference. Spatial Visium data were first divided into 28 tendon and 2,411 non-tendon spots. To address the small sample size, “pseudodonors” were created by assigning each tendon and an equal number of random non-tendon spots unique identifiers, while the remaining non-tendon spots were pooled into a single 29th pseudodonor. The *aggregate_and_filter()* function from Heumos et al.^147^ was adapted to generate 25 pseudoreplicates per donor based on estimated cell counts per spot. These were aggregated into a spatial reference AnnData object containing 1,425 observations and 14,208 genes. Both spatial and scRNA-seq data were log1pPF-normalised and concatenated to identify 5,000 highly variable genes (flavor=“cell_ranger”, batch_key=“modality” (“spatial” vs “single-cell”)).

To train the random forest classifier, class imbalances of tendon vs non-tendon data were addressed using Synthetic Minority Over-sampling Technique (SMOTE) from imbalanced-learn^148^ (v0.12.0) package (Lemaître et al., 2017). Spatial data were then split into train and test datasets at a ratio of 80:20. Scikit-learn’s^149^ (v1.3.0) *GridSearchCV()* was applied to select the best model hyperparameters for classifier training, and *RandomForestClassifier()* was trained with 2,000 decision trees, using parameters such as bootstrap sampling and square root feature selection at each split, achieving high (>0.96) out-of-bag, accuracy, precision, recall, and F1 scores. Trained classifier was then applied to the scRNA-seq dataset, predicting 4,318 tendon cells out of 108,617 total. Predictions were validated against spatial gene score-based annotations using the top 20 tendon-enriched genes (including *SCX*, *MKX*, *TNMD*, *ABI3BP*, *GAS2* etc), computed with *sc.tl.score_genes()* (Figure S3A,B).

Embryonic single-cell data were subset to the random forest classifier predictions. Only samples with at least 20 cells in each of the batch effect categories (‘samplename’, ‘kit’, ‘seq_protocol’, ’sex’) were retained. Samples sequenced with 5’ v1 kit as well as those sequenced using NovaSeq 6000 were removed due to significant batch effect confounding. Finally, genes detected in fewer than 5 cells were removed, leaving a total of 3,092 cells and 18,119 genes from 6 donors aged 6.5-9.3pcw. The data were re-normalised to log1pPF, and 4,000 highly variable genes were selected with flavor=“cell_ranger”. The effects of the cell cycle scores were regressed out to remove those significant sources of uninteresting variation (sc.pp.regress_out(adata, [‘S_score’, ‘G2M_score’])). The regressed counts were scaled, and the PCA was carried out with previously identified highly variable genes. The data were then clustered using Scanpy’s Leiden algorithm at 0.3 resolution, identifying 7 gene expression clusters. Those were manually annotated through marker gene exploration using the results from the between-cluster Wilcoxon rank-sum test. Marker sets from CZ CellxGene’s Cell Guide library for mesenchymal stem cells (CL:0000134) and chondrocytes (CL:0000138)^150^ were used for gene scoring to aid annotation.

#### Embryonic tendon RNA velocity analysis

scVelo (v0.2.5) preprocessing functions were applied to the combined, spliced, and unspliced counts matrices to re-normalise and filter them using the default settings. The neighbours and moments were calculated using regressed PCA embeddings. RNA Velocity analysis was carried out using a dynamical gene expression model by recovering dynamics, calculating velocities, and constructing a velocity graph with default settings. Differential kinetics test was then run using the top 100 dynamical genes, and the velocity was recalculated with those in mind.

#### Embryonic and foetal tendon trajectory and fate analysis

Embryonic and foetal tendon sample counts matrices were reorganised by sample age in ascending order. Harmony^151^ (v0.1.4) function *harmony.core.augmented_affinity_matrix()* was used with top 40 PC loadings (containing over 90% of total variation) and 20 k-nearest neighbours (knn) to construct an augmented affinity matrix that incorporated the developmental age information into the similarity measures between cells. Force-directed layouts based on this matrix are shown in Figure 4A. This matrix was then used to construct diffusion maps with 20 knn using Palantir^85^ (v1.3.1). The diffusion map embeddings were used to determine the multi-scale space of the data through 9 eigenvectors identified at the first eigengap. These multiscale space embeddings served as basis for tSNE dimensionality reduction, recalculation of the nearest neighbours (knn=15), and creation of the force-directed graphs shown in Figure 4C.

Palantir trajectory analysis was run using multiscale space embeddings (knn=20, num_waypoints=2000), with differentiation initiation and termination cells selected based on the most extreme values in multiscale diffusion components. CellRank^86^ (v1.5.1) Palantir pseudotime kernel was initiated, and the transition matrices were computed using a Generalized Perron Cluster Cluster Analysis (GPCCA) estimator with default settings. By analysing real eigenvalue plots, 3 distinct, stable macrostates representing terminal states were identified. Absorption probabilities were then computed using default settings, calculating lineage drivers for each terminal state. A single initial state was determined using a backward kernel based on the same process. Log1pPF-normalised, non-imputed counts were used to calculate correlations between lineage drivers shown in Figure 4F.

To bridge the gap between initial and terminal states, scVelo’s *scvelo.tl.recover_latent_time()* was used to recover latent time, and *scvelo.tl.paga(adata, groups=“cell_type”, root_key=“initial_states_probabilities”, end_key=“terminal_-states_probabilities”, use_time_prior=“palantir_pseudotime”)* was applied to calculate a directed PAGA incorporating Palantir pseudotime as prior. This was used as basis for plotting cell fate probabilities shown in Figure 4D,E.

#### Embryonic and foetal data integration and trajectory inference

Embryonic scRNA-seq and foetal snRNA-seq datasets were integrated using scGen^89^ (v2.1.1), with different sequencing runs specified as batches. The model was trained until convergence using default settings for a total of 28 epochs. Corrected latent space embeddings were used for the generation of a Harmony augmented affinity matrix. This was followed by Palantir multiscale diffusion map calculation and force-directed graph plotting, as described earlier. Following the generation of PAGA graphs, the Immune Cells were removed from further trajectory analysis due to them being disconnected from the rest of the cells even at low thresholds.

To reconstruct developmental trajectories, moscot^90^ (v0.3.3) was applied. The genes were re-filtered to retain only those with more than 20 counts. Next, a TemporalProblem object was initiated with scGen-corrected latent embeddings, and the proliferation and apoptosis scores were obtained. The temporal problem was solved using manually optimised parameters (epsilon=1e-3, tau_a=0.99, tau_b=0.999, scale_cost=“mean”, batch_size=1200). Cell transition scores were calculated to determine putative cell ancestors and descendants.

CellRank^88^ (v2.0.2) RealTimeKernel was initiated from the TemporalProblem object, and the transition matrices were computed with self_transitions=“all”, conn_weight=0.2, and threshold=“auto_local”. As before, GPCCA estimator was used. Overall, 7 macrostates were identified at the largest eigengap, out of which 4 corresponded to the major foetal tendon fibroblast cell types and were manually set to be terminal. The initial states were set manually to the putative tendon fibroblast precursor populations identified at 7.2 and 8.4pcw.

#### Embryonic, foetal and adult data integration

Processed, filtered, and annotated adult Achilles tendon (*N*=6, split by microanatomical regions; Table S1) and quadriceps tendon (*N*=7, healthy or torn midbodies; Table S1) datasets provided by the Tendon Seed Network^124,125^ were concatenated with processed and annotated embryonic and foetal tendon datasets. Cells with fewer than 200 genes and genes in fewer than 30 counts were filtered out, yielding 176,691 cells and 32,869 genes. Top 7,000 highly variable genes were selected in a batch-aware manner using *scanpy.pp.highly_variable-_genes(batch_key=”sampletype”)*. Data integration was performed using scVI modelling on unnormalised, highly variable counts, with library preparation batches (“libbatch”), “sampletype”, and the cell cycle phase scores (“G2M_score” and “S_score”) specified as main batch effects. The model’s hyperparameters were optimised with scVI’s autotune to ‘n_hidden’: 256, ’n_latent’: 50, ’n_layers’: 1, ’dropout_rate’: 0.1, ’gene_likelihood’: ’zinb’, and ‘dispersion’: ‘gene-batch’. The model was trained for 80 epochs until training and validation set convergence. Next, cell type labels were harmonised using the scANVI model. The resulting embeddings were used for the computation of neighbourhood graphs and dimensionality reduction with UMAP.

The data were normalised to log1pPF and subsequently subjected to Wilcoxon rank-sum test differential gene expression analysis (*scanpy.tl.rank_genes_groups()*), which was performed separately for foetal and adult cell populations. For each specified cell type within foetal tendons, the top 50 differentially expressed genes were extracted. Marker gene overlap analysis using *scanpy.tl.marker_gene_overlap()* was conducted to quantify and visualise transcriptional similarities between the foetal top 50 DEGs and the marker genes derived from the adult differential expression analysis. Normalisation of overlap scores was performed relative to the foetal DEGs, providing a proportion of foetal marker genes detected in each adult cell type (Figure S4B).

To achieve more robust cell type label harmonisation, CellHint^152^ (v1.0.0) was applied to the different tendon types across the embryonic, foetal, and adult age groups. Specifically, *cellhint.harmonize()* function was used to calculate euclidean distances between cells within batch-corrected scVI embeddings, constructing a harmonisation graph of cell type annotations. The datasets were specified to be harmonised sequentially, from embryonic to adult stages.

#### Gene regulatory network analysis

Single-cell regulatory network inference and clustering (SCENIC) analysis was performed for embryonic, foetal, and adult quadriceps tendon datasets using pyscenic^153^ (v0.12.1). Gene regulatory networks were inferred using the GRNBoost2 algorithm with default settings. The input data were raw, unnormalised counts. Human transcription factors were predefined using the ’allTFs_hg38.txt’ list from the Aertslab GitHub repository. The resulting TF-gene interactions were utilised to infer co-expression modules, identify enriched motifs, and predict regulons with the *pyscenic ctx* command. Genome rankings (.feather) and motif annotation (.tbl) v10 files were obtained from the Aertslab cistarget resources webpage. A total of 335 embryonic regulons, 285 regulons from 12 pcw foetal samples, 237 regulons from 17 pcw foetal samples, 286 regulons from 20 pcw foetal samples, and 193 adult regulons were identified.

Regulon activity within individual cells was quantified using the *pyscenic aucell* command, with an AUC threshold of 0.1 applied to the embryonic dataset and the default threshold of 0.05 applied to the foetal and adult datasets. Binarization of regulon activity was achieved using a Gaussian mixture model, resulting in matrices indicating active and inactive regulons per cell.

The binarized regulon matrices underwent several transformations to extract and normalise regulon activation patterns across different cell populations. Initially, cells were grouped according to their cell type annotations, allowing the aggregation of active regulons by calculating the sum of active cells per regulon within each group. Subsequently, these summaries were normalised based on the total cell count per group to account for variations in cell numbers. Next, normalised regulon activation frequencies within each group were further z-score normalised, facilitating comparative analysis across developmental stages and conditions. The z-scored regulon activity patterns were visualised using custom clustermaps. Individual regulons were explored and summarised using the STRING database (*stringdb,*^154^ v12.0). Large regulons were subjected to pathway analysis with gProfiler g:GOSt^144^ as previously described.

## QUANTIFICATION AND STATISTICAL ANALYSIS

### Quantification of foetal cell type compositional changes

Cell type compositional changes between 12 and 20 pcw foetal tendons were assessed using the scCODA^155^ (v0.1.9) package. A MuData object was generated from the foetal tendon snRNA-seq dataset, and scCODA was run with modality_key=’coda’. The Nervous System Cells cluster was automatically selected as the reference. The model was trained using 11,000 NUTS (No-U-Turn Sampler) iterations, and statistically credible shifts in cell type proportions were identified. Results are summarised in Figure 1E.

### Quantifying metrics from tendon histology

QuPath^156^ (v0.4.3) classification functions were used to manually delineate tendon regions within H&E-stained tissue samples. From these regions, 3-6 random 250×250 μm tiles were extracted and processed using ImageJ (v1.53t). First, haematoxylin and eosin channels were separated using colour deconvolution. Subsequently, the haematoxylin channel was used for nuclei segmentation using the StarDist^157^ ImageJ plugin (v0.3.0) (see Figure S10).

Following this segmentation, quantitative analyses were performed to determine the number of nuclei, the mean area of the nuclei, and the maximum distance to the centroid of each nucleus (Figure S9). Statistical analysis was performed using scipy^158^ (v1.11.1), statsmodels^159^ (v0.14.0), and scikit-posthocs^160^ (v0.9.0). The normality of the data distributions for the number of nuclei, their areas, and maximum distances to their centres was assessed using the Shapiro-Wilk test as well as visual inspections of histograms and Q-Q plots. For data that approximated normal distributions – the nucleus area and maximum distance to nucleus centroid variables – analysis of variance (ANOVA) was performed to test for significant differences across different age groups. For the number of nuclei, which exhibited an almost bimodal distribution, the Kruskal-Wallis H-test was used to assess significant differences across age groups. Post-hoc analysis was conducted to identify specific group differences. Tukey’s Honest Significant Difference test was used following the ANOVA, and Dunn’s test with Bonferroni correction was employed after the Kruskal-Wallis H-test.

### Foetal tendon pseudobulk differential gene expression analysis

For differential gene expression (DGE) analysis of foetal cell types across different ages, the raw single-nuclei sequencing counts were converted into “pseudobulk” expression profiles using custom functions inspired by Heumos et al.^147^ Briefly, the data were partitioned by distinct cell types, and a donor-filtering criterion was applied to retain only tissue donors with a minimum cell count threshold of 30. Three pseudoreplicates were then generated by randomly subsampling cells within retained donors, and these pseudoreplicates were subsequently transformed into pseudobulk samples by aggregating expression counts. Metadata associated with donors, experimental conditions, and cell identity were preserved.

DGE analysis was carried out individually for each of the cell type pseudobulked counts using DESeq2^161^ (v1.40.2). Both LRT and Wald tests were done to assess the differences between different developmental timepoints and tissue types. LRT results with p-adjusted values of <0.01 were clustered to investigate common patterns in gene expression changes over time. The resulting cluster gene lists were subjected to pathway analysis against all expressed genes using the gProfiler g:GOSt tool^144^ with Benjamini-Hochberg FDR set to <0.05 and pathway term sizes restricted to between 20 and 500 terms. Shared or cell type-specific terms for each cluster were then isolated and manually grouped into common functional categories to reduce data dimensionality and enable comparisons.

### Foetal and adult tendon pseudobulk differential gene expression analysis

For DGE analysis across different foetal and adult cell types, the raw single-nuclei sequencing counts were converted into grouped “donor + tissue type + cell type” pseudobulk expression profiles using custom functions inspired by Heumos et al.^147^ Differential gene expression analysis was carried out separately for the specified groups containing specific CellHint-paired foetal and adult cell types using DESeq2^161^ (v1.40.2). Wald tests were used to identify all genes that were significantly up- or downregulated in adult cell types compared to the foetal cell types (with *p-adjusted* < 0.01, *log2FC* > +/-1). These genes were used for pathway analysis against all expressed genes using the gProfiler g:GOSt tool^144^ with Benjamini-Hochberg FDR set to <0.01 and term sizes restricted to between 20 and 500 terms. The identified up- or downregulated GO:BP terms for each CellHint-aligned pair of foetal and adult fibroblasts were compared against each other using Venn diagrams made with Venny^162^ (v2.1.0; Figure S12). The pair-specific pathways were then manually summarised by common functions into broad categories to enable comparisons (Figures S13-16).

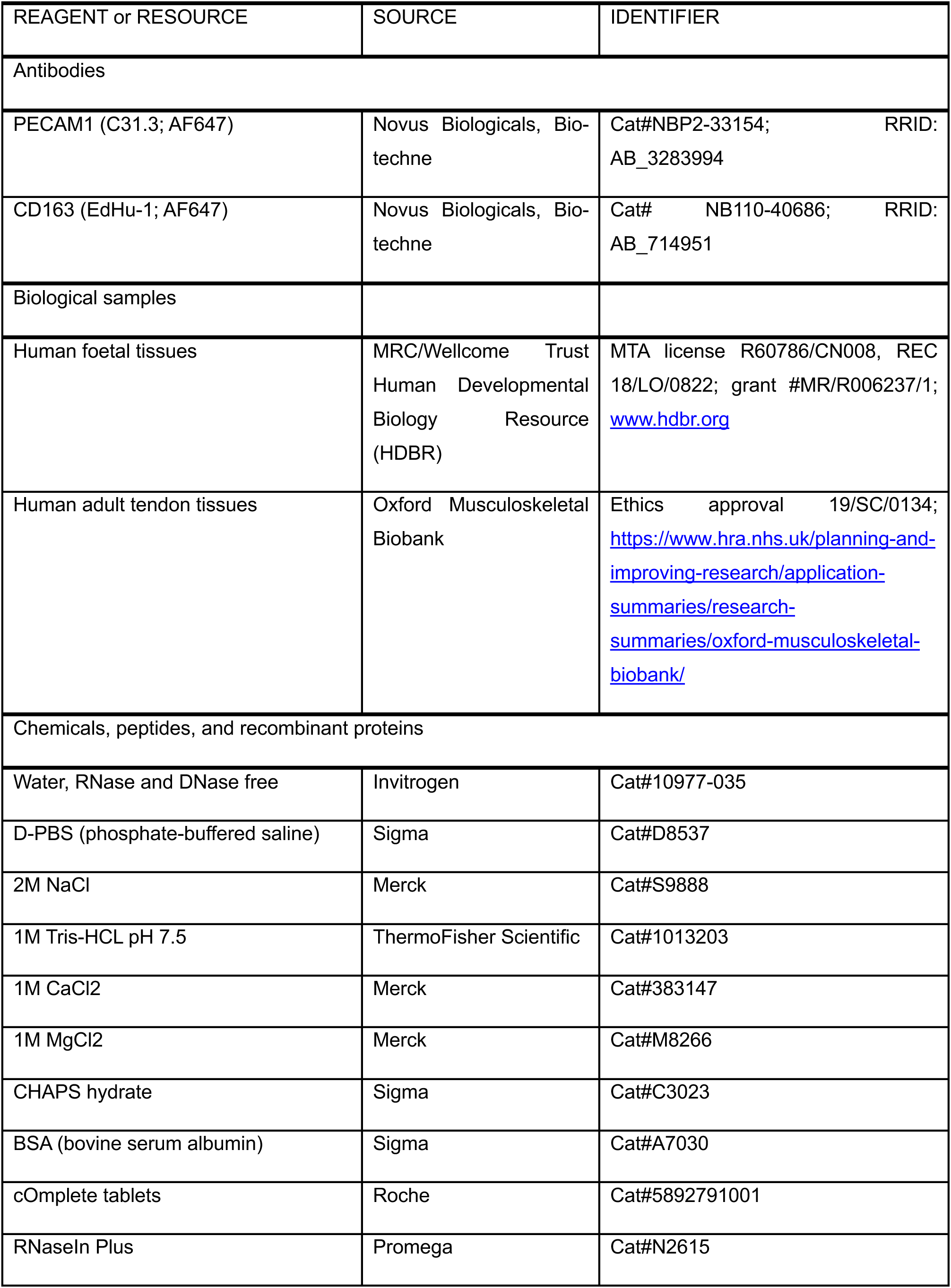
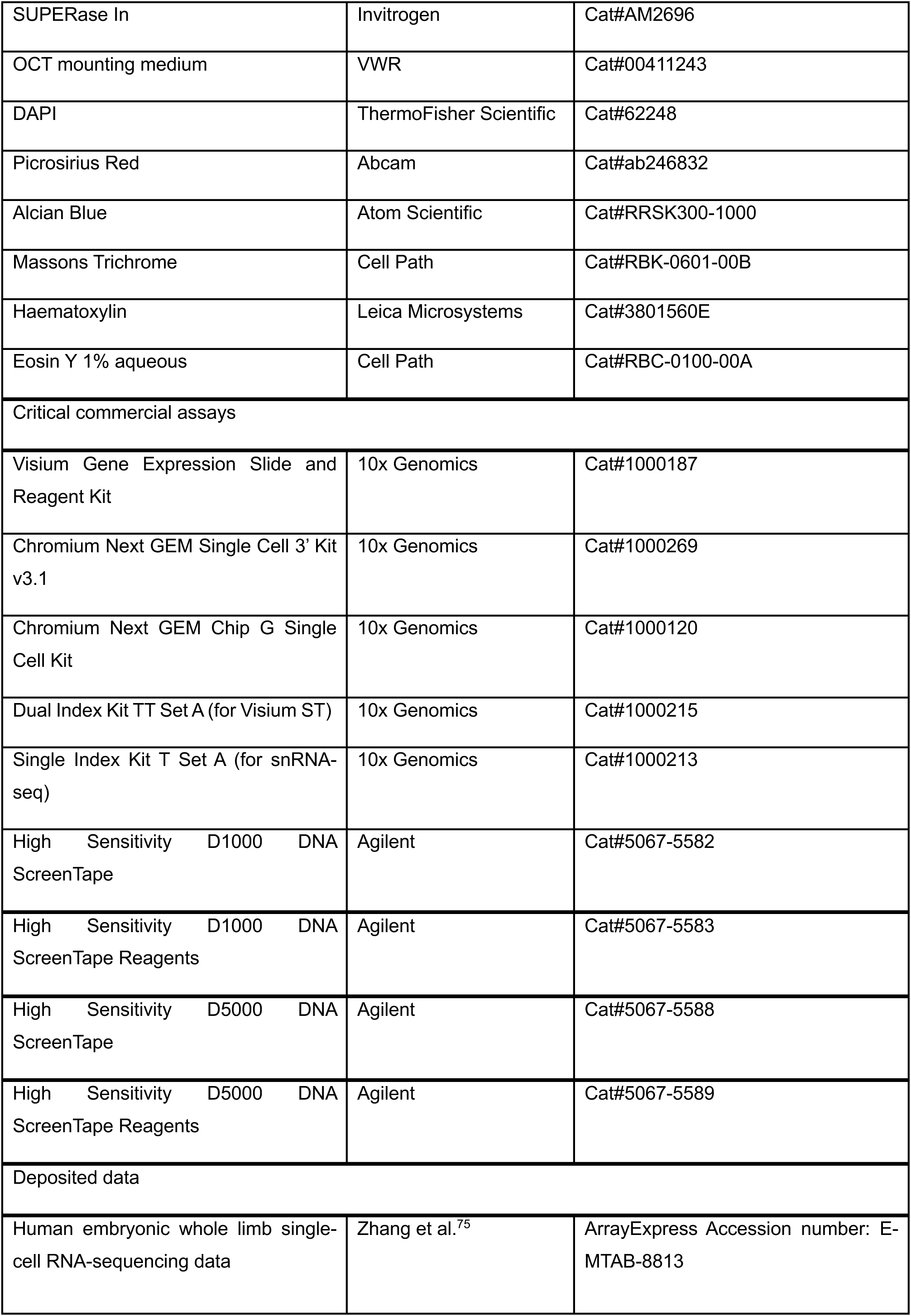
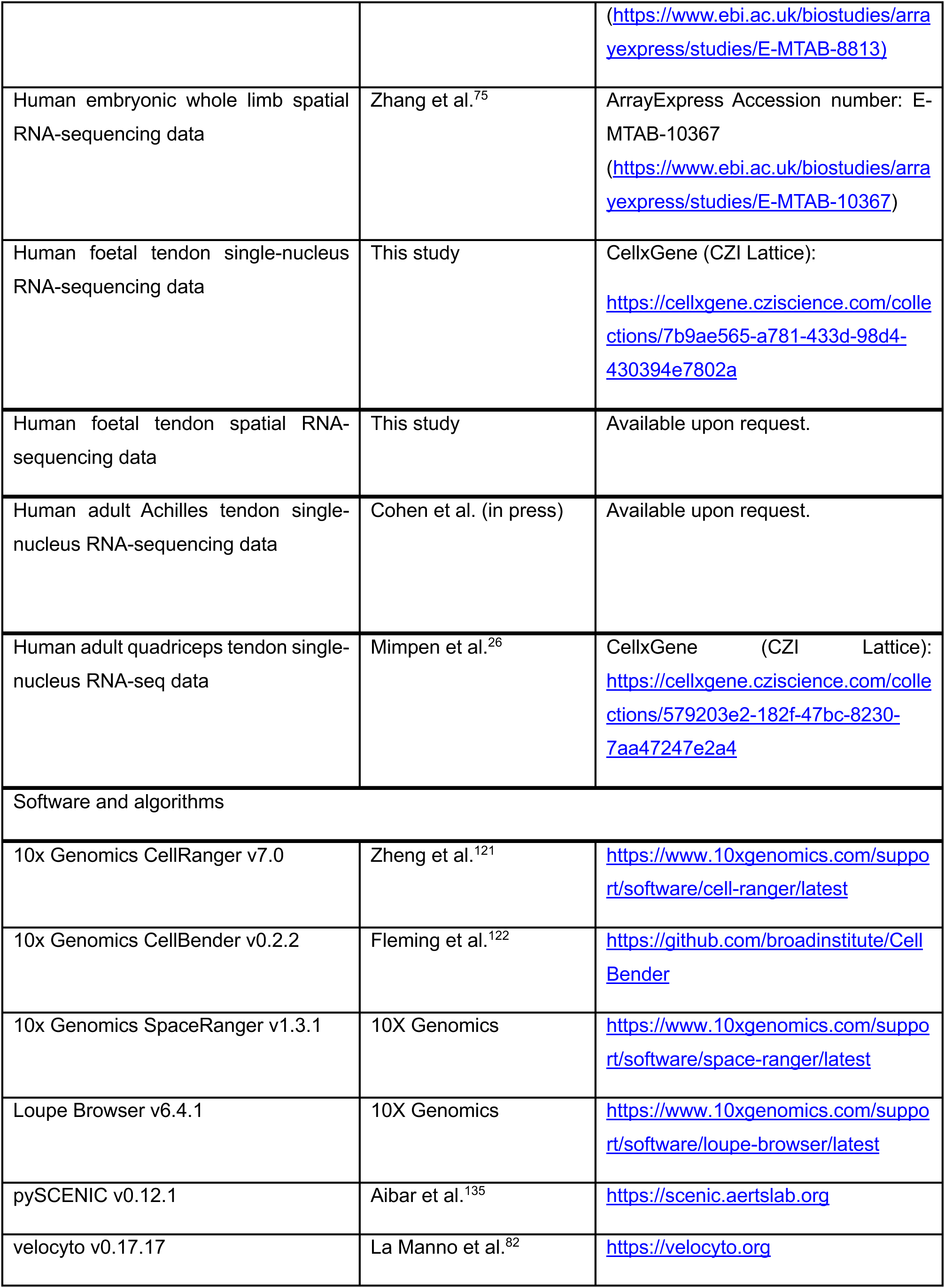
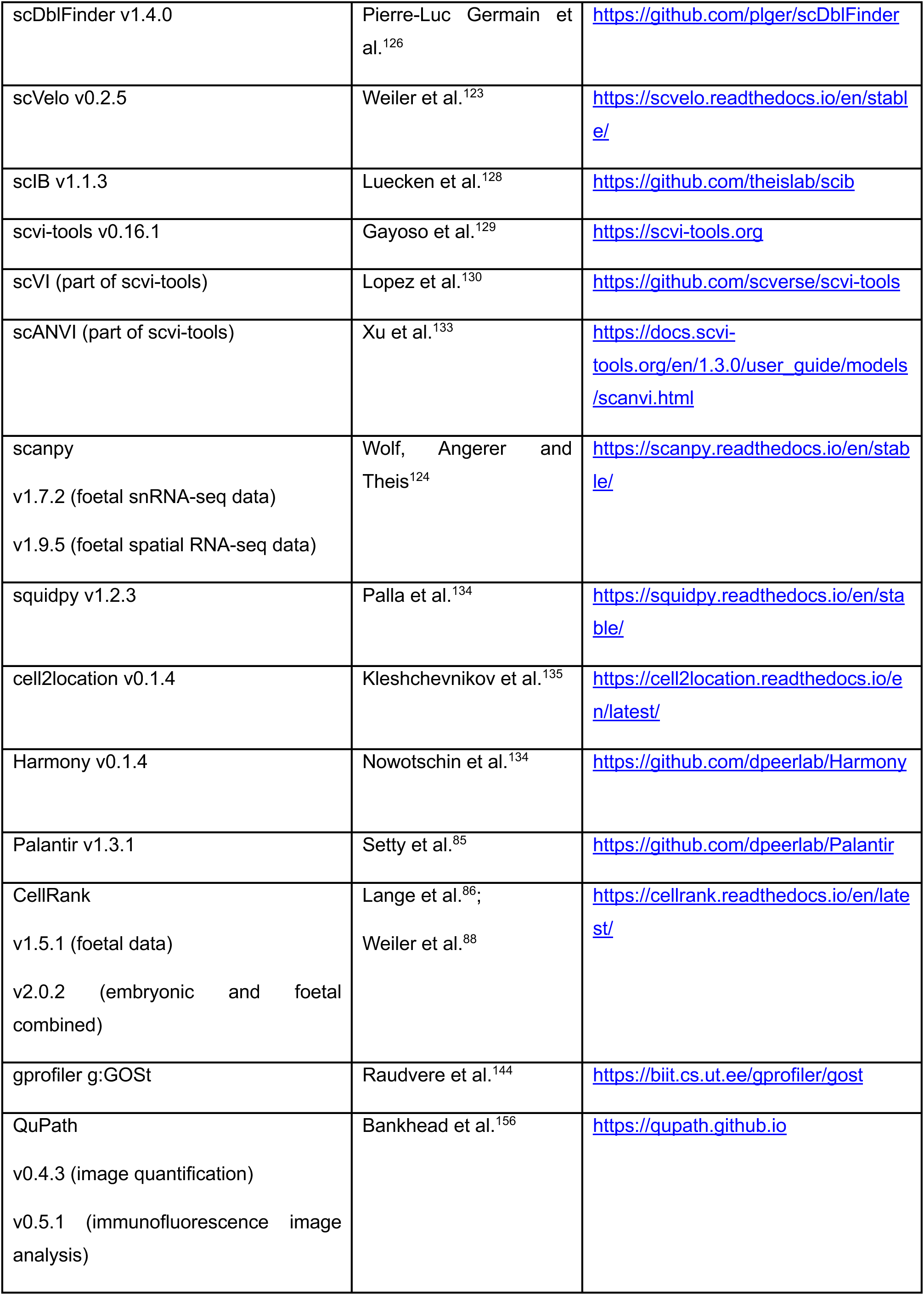
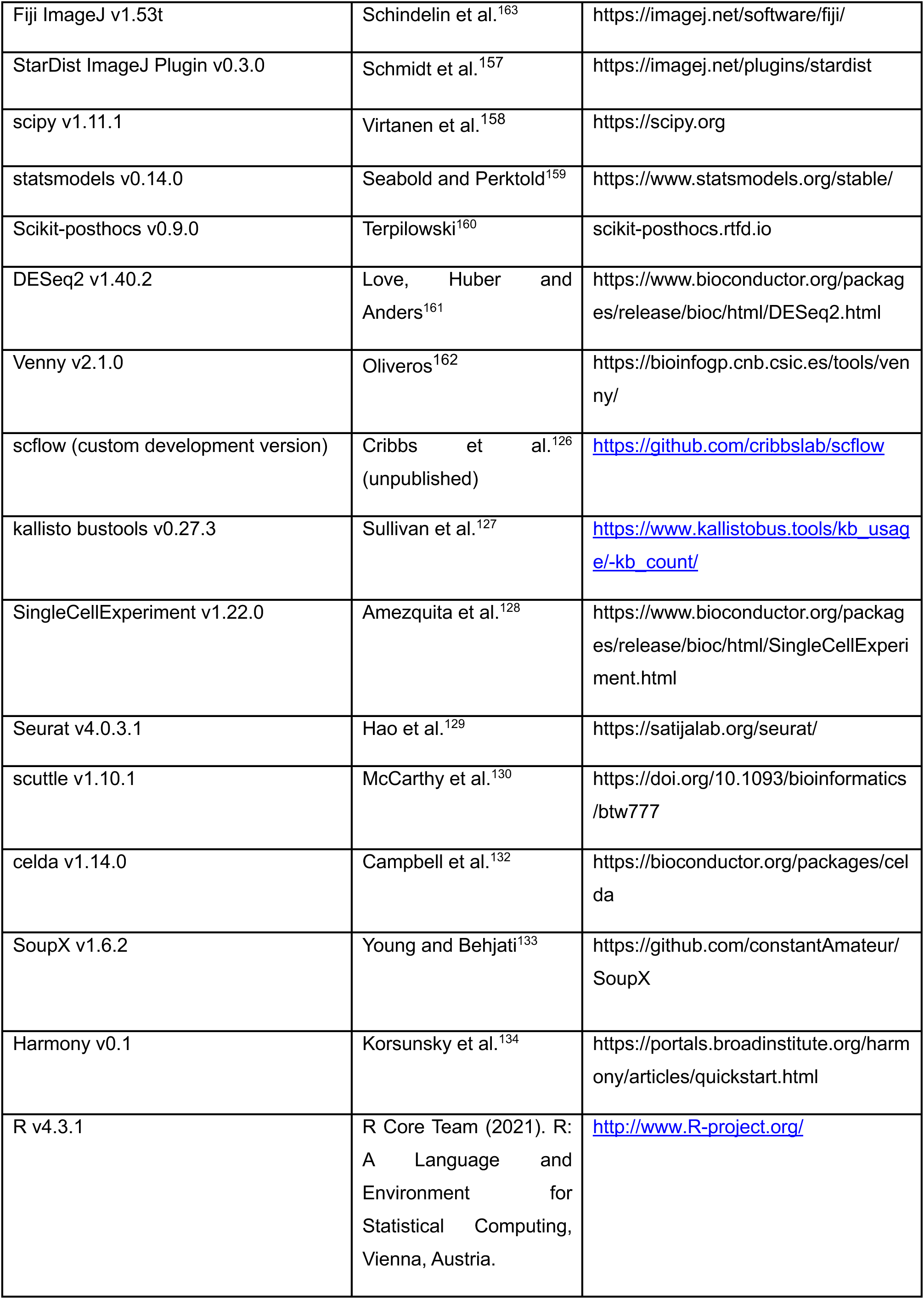
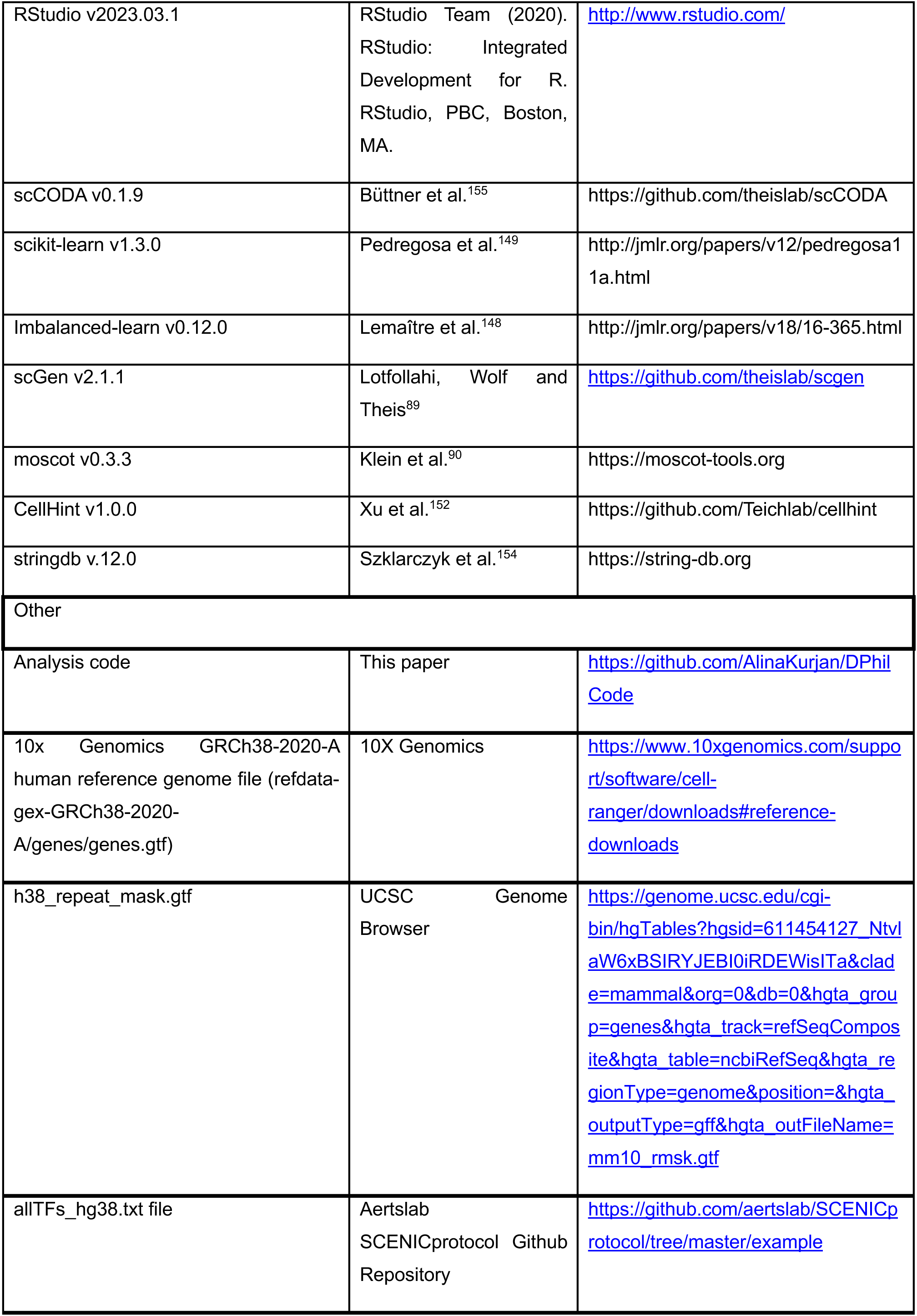
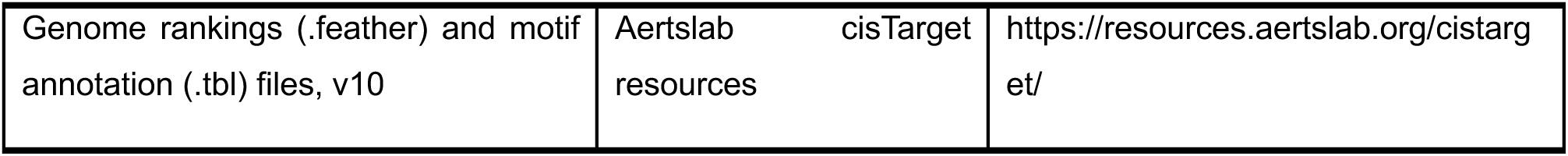
KEY RESOURCES TABLE

## REFERENCES

1. Magnusson, S. P., Langberg, H. & Kjaer, M. The pathogenesis of tendinopathy: balancing the response to loading. Nat. Rev. Rheumatol. 6, 262–268 (2010).

2. Malliaras, P., Barton, C. J., Reeves, N. D. & Langberg, H. Achilles and Patellar Tendinopathy Loading Programmes. Sports Med. 43, 267–286 (2013).

3. Simpson, M., Rio, E. & Cook, J. At What Age Do Children and Adolescents Develop Lower Limb Tendon Pathology or Tendinopathy? A Systematic Review and Meta-analysis. Sports Med. 46, 545–557 (2016).

4. Millar, N. L. et al. Tendinopathy. Nat. Rev. Dis. Primer 7, 1–21 (2021).

5. Ansorge, H. L., Adams, S., Birk, D. E. & Soslowsky, L. J. Mechanical, Compositional, and Structural Properties of the Post-natal Mouse Achilles Tendon. Ann. Biomed. Eng. 39, 1904–1913 (2011).

6. Beredjiklian, P. K. et al. Regenerative versus reparative healing in tendon: A study of biomechanical and histological properties in fetal sheep. Ann. Biomed. Eng. 31, 1143–1152 (2003).

7. Chan, B. P., Fu, S. C., Qin, L., Rolf, C. & Chan, K. M. Pyridinoline in relation to ultimate stress of the patellar tendon during healing: An animal study. J. Orthop. Res. 16, 597–603 (1998).

8. Favata, M. et al. Regenerative properties of fetal sheep tendon are not adversely affected by transplantation into an adult environment. J. Orthop. Res. 24, 2124– 2132 (2006).

9. Howell, K. et al. Novel Model of Tendon Regeneration Reveals Distinct Cell Mechanisms Underlying Regenerative and Fibrotic Tendon Healing. Sci. Rep. 7, (2017).

10. Lui, P. P. Y., Cheuk, Y. C., Lee, Y. W. & Chan, K. M. Ectopic chondro-ossification and erroneous extracellular matrix deposition in a tendon window injury model. J. Orthop. Res. 30, 37–46 (2012).

11. Garner, W. L., McDonald, J. A., Koo, M., Kuhn, C. & Weeks, P. M. Identification of the collagen-producing cells in healing flexor tendons. Plast. Reconstr. Surg. 83, 875–879 (1989).

12. Grinstein, M. et al. A quiescent resident progenitor pool is the central organizer of tendon healing. *bioRxiv* 2022.02.02.478533 (2022) doi:10.1101/2022.02.02.478533.

13. Harvey, T., Flamenco, S. & Fan, C. M. A Tppp3+Pdgfra+ tendon stem cell population contributes to regeneration and reveals a shared role for PDGF signalling in regeneration and fibrosis. Nat. Cell Biol. 21, 1490 (2019).

14. Pakshir, P. & Hinz, B. The big five in fibrosis: Macrophages, myofibroblasts, matrix, mechanics, and miscommunication. Matrix Biol. 68–69, 81–93 (2018).

15. Snedeker, J. G. & Foolen, J. Tendon injury and repair – A perspective on the basic mechanisms of tendon disease and future clinical therapy. Acta Biomater. 63, 18– 36 (2017).

16. Voleti, P. B., Buckley, M. R. & Soslowsky, L. J. Tendon Healing: Repair and Regeneration. Annu. Rev. Biomed. Eng. 14, 47–71 (2012).

17. Walia, B., Li, T. M., Crosio, G., Montero, A. M. & Huang, A. H. Axin2-lineage cells contribute to neonatal tendon regeneration. Httpsezproxy-Prdbodleianoxacuk21021010800300820720222036732 1–14 (2022) doi:10.1080/03008207.2022.2036732.

18. Yin, Z. et al. Single-cell analysis reveals a nestin+ tendon stem/progenitor cell population with strong tenogenic potentiality. Sci. Adv. 2, 1600874 (2016).

19. De Micheli, A. J. et al. Single-cell transcriptomic analysis identifies extensive heterogeneity in the cellular composition of mouse Achilles tendons. Am. J. Physiol. - Cell Physiol. 319, C885–C894 (2020).

20. Kendal, A. R. et al. Multi-omic single cell analysis resolves novel stromal cell populations in healthy and diseased human tendon. Sci. Rep. 10, 1–14 (2020).

21. Akbar, M. et al. Single cell and spatial transcriptomics in human tendon disease indicate dysregulated immune homeostasis. Ann. Rheum. Dis. 80, 1494–1497 (2021).

22. Steffen, D., Mienaltowski, M. & Baar, K. Spatial gene expression in the adult rat patellar tendon. Matrix Biol. Plus 19–20, 100138 (2023).

23. Ackerman, J. E. et al. Defining the spatial-molecular map of fibrotic tendon healing and the drivers of Scleraxis-lineage cell fate and function. Cell Rep. 41, 111706 (2022).

24. Mimpen, J. Y. et al. Single nucleus and spatial transcriptomic profiling of human healthy hamstring tendon. 2022.12.19.521110 Preprint at 10.1101/2022.12.19.521110 (2022).

25. Fu, W., Yang, R. & Li, J. Single-cell and spatial transcriptomics reveal changes in cell heterogeneity during progression of human tendinopathy. BMC Biol. 21, 132 (2023).

26. Mimpen, J. Y. et al. Exploring cellular changes in ruptured human quadriceps tendons at single-cell resolution. J. Physiol. **n/a**, (2025).

27. McNeilly, C. M., Banes, A. J., Benjamin, M. & Ralphs, J. R. Tendon cells in vivo form a three dimensional network of cell processes linked by gap junctions. J. Anat. 189 (Pt 3), 593–600 (1996).

28. Kannus, P. Structure of the tendon connective tissue. Scand. J. Med. Sci. Sports 10, 312–320 (2000).

29. Franchi, M., Trirè, A., Quaranta, M., Orsini, E. & Ottani, V. Collagen Structure of Tendon Relates to Function. Sci. World J. 7, 404–420 (2007).

30. Kalson, N. S., Lu, Y., Taylor, S. H., Holmes, D. F. & Kadler, K. E. A structure-based extracellular matrix expansion mechanism of fibrous tissue growth. eLife 4, 1–22 (2015).

31. Steffen, D., Avey, A., Mienaltowski, M. J. & Baar, K. The rat Achilles and patellar tendons have similar increases in mechanical properties but become transcriptionally divergent during postnatal development. J. Physiol. 601, 3869– 3884 (2023).

32. Bi, Y. et al. Identification of tendon stem/progenitor cells and the role of the extracellular matrix in their niche. Nat. Med. 13, 1219–1227 (2007).

33. Lui, P. P. Y. & Chan, K. M. Tendon-Derived Stem Cells (TDSCs): From Basic Science to Potential Roles in Tendon Pathology and Tissue Engineering Applications. Stem Cell Rev. Rep. 7, 883–897 (2011).

34. Staverosky, J. A., Pryce, B. A., Watson, S. S. & Schweitzer, R. Tubulin polymerization-promoting protein family member 3, Tppp3, is a specific marker of the differentiating tendon sheath and synovial joints. Dev. Dyn. 238, 685–692 (2009).

35. Sakabe, T. et al. Transcription factor scleraxis vitally contributes to progenitor lineage direction in wound healing of adult tendon in mice. J. Biol. Chem. 293, 5766–5780 (2018).

36. Dyment, N. A. et al. The Paratenon Contributes to Scleraxis-Expressing Cells during Patellar Tendon Healing. PLOS ONE 8, e59944 (2013).

37. Maeda, T. et al. Conversion of Mechanical Force into TGF-β-Mediated Biochemical Signals. Curr. Biol. 21, 933–941 (2011).

38. Ansorge, H. L. et al. Recapitulation of the Achilles tendon mechanical properties during neonatal development: a study of differential healing during two stages of development in a mouse model. J. Orthop. Res. 30, 448–456 (2012).

39. Nichols, A. E. C., Wagner, N. W., Ketonis, C. & Loiselle, A. E. Epitenon-derived cells comprise a distinct progenitor population that contributes to both tendon fibrosis and regeneration following acute injury. *bioRxiv* 2023.01.30.526242 (2023) doi:10.1101/2023.01.30.526242.

40. Howell, K. L. et al. Macrophage depletion impairs neonatal tendon regeneration. FASEB J. Off. Publ. Fed. Am. Soc. Exp. Biol. 35, e21618 (2021).

41. Vinestock, R. C. et al. Neonatal Enthesis Healing Involves Noninflammatory Acellular Scar Formation through Extracellular Matrix Secretion by Resident Cells. Am. J. Pathol. 192, 1122–1135 (2022).

42. Moser, H. L. et al. Genetic Lineage Tracing of Targeted Cell Populations During Enthesis Healing. J. Orthop. Res. Off. Publ. Orthop. Res. Soc. 36, 3275–3284 (2018).

43. Moser, H. L. et al. Cell lineage tracing and functional assessment of supraspinatus tendon healing in an acute repair murine model. J. Orthop. Res. 39, 1789–1799 (2021).

44. Chen, C. H. et al. Tendon Healing In Vivo: Gene Expression and Production of Multiple Growth Factors in Early Tendon Healing Period. *J*. Hand Surg. 33, 1834– 1842 (2008).

45. Kaji, D. A., Howell, K. L., Balic, Z., Hubmacher, D. & Huang, A. H. Tgfβ signaling is required for tenocyte recruitment and functional neonatal tendon regeneration. eLife 9, 1–19 (2020).

46. Lopez, R., Regier, J., Cole, M. B., Jordan, M. I. & Yosef, N. Deep generative modeling for single-cell transcriptomics. Nat. Methods 15, 1053–1058 (2018).

47. Kleshchevnikov, V. et al. Cell2location maps fine-grained cell types in spatial transcriptomics. Nat. Biotechnol. 40, 661–671 (2022).

48. Birk, D. E. & Brückner, P. Collagens, Suprastructures, and Collagen Fibril Assembly. in The Extracellular Matrix: an Overview (ed. Mecham, R. P.) 77–115 (Springer, Berlin, Heidelberg, 2011). doi:10.1007/978-3-642-16555-9_3.

49. Docheva, D., Hunziker, E. B., Fässler, R. & Brandau, O. Tenomodulin Is Necessary for Tenocyte Proliferation and Tendon Maturation. Mol. Cell. Biol. 25, 699–705 (2005).

50. Dex, S., Lin, D., Shukunami, C. & Docheva, D. TENOgenic MODULating INsider factor: systematic assessment on the functions of tenomodulin gene. Gene 587, 1–17 (2016).

51. Shukunami, C. et al. Scleraxis is a transcriptional activator that regulates the expression of Tenomodulin, a marker of mature tenocytes and ligamentocytes. Sci. Rep. 8, 3155 (2018).

52. Shukunami, C., Takimoto, A., Oro, M. & Hiraki, Y. Scleraxis positively regulates the expression of tenomodulin, a differentiation marker of tenocytes. Dev. Biol. 298, 234–247 (2006).

53. Anderson, D. M. et al. Mohawk is a novel homeobox gene expressed in the developing mouse embryo. Dev. Dyn. 235, 792–801 (2006).

54. Ito, Y. et al. The Mohawk homeobox gene is a critical regulator of tendon differentiation. Proc. Natl. Acad. Sci. U. S. A. 107, 10538–10542 (2010).

55. Liu, W. et al. The Atypical Homeodomain Transcription Factor Mohawk Controls Tendon Morphogenesis. Mol. Cell. Biol. 30, 4797–4807 (2010).

56. Wang, W. et al. Collagen XXIV (Col24α1) Promotes Osteoblastic Differentiation and Mineralization through TGF-β/Smads Signaling Pathway. Int. J. Biol. Sci. 8, 1310–1322 (2012).

57. Takahata, Y. et al. Smoc1 and Smoc2 regulate bone formation as downstream molecules of Runx2. *Commun*. Biol. 4, 1–11 (2021).

58. He, Z. et al. Enpp1 mutations promote upregulation of hedgehog signaling in heterotopic ossification with aging. J. Bone Miner. Metab. 42, 681–698 (2024).

59. Jenkins, E., Moss, J. B., Pace, J. M. & Bridgewater, L. C. The new collagen gene COL27A1 contains SOX9-responsive enhancer elements. Matrix Biol. J. Int. Soc. Matrix Biol. 24, 177–184 (2005).

60. Alcaide-Ruggiero, L., Molina-Hernández, V., Granados, M. M. & Domínguez, J. M. Main and Minor Types of Collagens in the Articular Cartilage: The Role of Collagens in Repair Tissue Evaluation in Chondral Defects. Int. J. Mol. Sci. 22, 13329 (2021).

61. de Castro, L. F. et al. Secreted frizzled related-protein 2 (Sfrp2) deficiency decreases adult skeletal stem cell function in mice. Bone Res. 9, 1–12 (2021).

62. Bastepe, M. GNAS mutations and heterotopic ossification. Bone 109, 80–85 (2018).

63. Kalamajski, S., Aspberg, A., Lindblom, K., Heinegård, D. & Oldberg, Å. Asporin competes with decorin for collagen binding, binds calcium and promotes osteoblast collagen mineralization. Biochem. J. 423, 53–59 (2009).

64. Fan, R., Yan, X. & Zhang, W. Relationship between asporin and extracellular matrix behavior: A literature review. Medicine (Baltimore) 101, e32490 (2022).

65. Maccarana, M. et al. Asporin-deficient mice have tougher skin and altered skin glycosaminoglycan content and structure. PLOS ONE 12, e0184028 (2017).

66. Charvet, B. et al. Knockdown of col22a1 gene in zebrafish induces a muscular dystrophy by disruption of the myotendinous junction. Development 140, 4602– 4613 (2013).

67. Koch, M. et al. A novel marker of tissue junctions, collagen XXII. J. Biol. Chem. 279, 22514–22521 (2004).

68. Zhang, C.-H. et al. Creb5 coordinates synovial joint formation with the genesis of articular cartilage. Nat. Commun. 13, 7295 (2022).

69. Zhang, C.-H. et al. Creb5 establishes the competence for Prg4 expression in articular cartilage. Commun. Biol. 4, 1–17 (2021).

70. Hayashi, M. et al. The Effect of Lubricin on the Gliding Resistance of Mouse Intrasynovial Tendon. PLOS ONE 8, e83836 (2013).

71. Thornton, G. M. et al. Aging affects mechanical properties and lubricin/PRG4 gene expression in normal ligaments. J. Biomech. 48, 3306–3311 (2015).

72. Kohrs, R. T. et al. Tendon fascicle gliding in wild type, heterozygous, and lubricin knockout mice. J. Orthop. Res. 29, 384–389 (2011).

73. Sun, Y.-L. et al. Lubricin in human achilles tendon: The evidence of intratendinous sliding motion and shear force in achilles tendon. J. Orthop. Res. 33, 932–937 (2015).

74. Sun, Y. et al. Expression and mapping of lubricin in canine flexor tendon. J. Orthop. Res. 24, 1861–1868 (2006).

75. Reuvers, J. et al. The mechanical properties of tail tendon fascicles from lubricin knockout, wild type and heterozygous mice. J. Struct. Biol. 176, 41–45 (2011).

76. Ostner, J., Carcy, S. & Müller, C. L. tascCODA: Bayesian Tree-Aggregated Analysis of Compositional Amplicon and Single-Cell Data. Front. Genet. 12, (2021).

77. Zhang, B. et al. A human embryonic limb cell atlas resolved in space and time. Nature 1–11 (2023) doi:10.1038/s41586-023-06806-x.

78. Grimaldi, A., Comai, G., Mella, S. & Tajbakhsh, S. Identification of bipotent progenitors that give rise to myogenic and connective tissues in mouse. eLife 11, e70235 (2022).

79. Yamamoto, S. et al. Hoxa13 regulates expression of common Hox target genes involved in cartilage development to coordinate the expansion of the autopodal anlage. Dev. Growth Differ. 61, 228–251 (2019).

80. Bonnin, M.-A. et al. Six1 is not involved in limb tendon development, but is expressed in limb connective tissue under Shh regulation. Mech. Dev. 122, 573– 585 (2005).

81. Wu, W. et al. The Role of Six1 in the Genesis of Muscle Cell and Skeletal Muscle Development. Int. J. Biol. Sci. 10, 983–989 (2014).

82. Nassari, S. et al. The chemokines CXCL12 and CXCL14 differentially regulate connective tissue markers during limb development. Sci. Rep. 7, 17279 (2017).

83. Stricker, S. et al. Odd-skipped related genes regulate differentiation of embryonic limb mesenchyme and bone marrow mesenchymal stromal cells. Stem Cells Dev. 21, 623–633 (2012).

84. La Manno, G. et al. RNA velocity of single cells. Nature 560, 494–498 (2018).

85. Setty, M. et al. Characterization of cell fate probabilities in single-cell data with Palantir. Nat. Biotechnol. 37, 451–460 (2019).

86. Lange, M. et al. CellRank for directed single-cell fate mapping. Nat. Methods 19, 159–170 (2022).

87. Reuter, B., Klein, M. & Lange, M. pyGPCCA - python GPCCA: Generalized Perron Cluster Cluster Analysis package to coarse-grain reversible and non-reversible Markov state models. Zenodo 10.5281/zenodo.6914001 (2022).

88. Weiler, P., Lange, M., Klein, M., Pe’er, D. & Theis, F. CellRank 2: unified fate mapping in multiview single-cell data. Nat. Methods 21, 1196–1205 (2024).

89. Lotfollahi, M., Wolf, F. A. & Theis, F. J. scGen predicts single-cell perturbation responses. Nat. Methods 16, 715–721 (2019).

90. Klein, D. et al. Mapping cells through time and space with moscot. Nature 638, 1065–1075 (2025).

91. Fearon, A., Dahlstrom, J. E., Twin, J., Cook, J. & Scott, A. The Bonar score revisited: Region of evaluation significantly influences the standardized assessment of tendon degeneration. J. Sci. Med. Sport 17, 346–350 (2014).

92. Khan, K. M., Cook, J. L., Bonar, F., Harcourt, P. & Åstrom, M. Histopathology of Common Tendinopathies. Sports Med. 27, 393–408 (1999).

93. Maffulli, N., Longo, U. G., Franceschi, F., Rabitti, C. & Denaro, V. Movin and Bonar Scores Assess the Same Characteristics of Tendon Histology. Clin. Orthop. 466, 1605–1611 (2008).

94. Broner, E. C. et al. Doublecortin-Like Kinase 1 (DCLK1) Is a Novel NOTCH Pathway Signaling Regulator in Head and Neck Squamous Cell Carcinoma. Front. Oncol. 11, 677051 (2021).

95. Vijai, M., Baba, M., Ramalingam, S. & Thiyagaraj, A. DCLK1 and its interaction partners: An effective therapeutic target for colorectal cancer. Oncol. Lett. 22, 850 (2021).

96. Kong, D. et al. Platelet-Derived Growth Factor-D Overexpression Contributes to Epithelial-Mesenchymal Transition of PC3 Prostate Cancer Cells. Stem Cells 26, 1425–1435 (2008).

97. Folestad, E., Kunath, A. & Wågsäter, D. PDGF-C and PDGF-D signaling in vascular diseases and animal models. Mol. Aspects Med. 62, 1–11 (2018).

98. Al-Hattab, D. S. et al. Scleraxis regulates Twist1 and Snai1 expression in the epithelial-to-mesenchymal transition. Am. J. Physiol.-Heart Circ. Physiol. 315, H658–H668 (2018).

99. Pryce, B. A., Brent, A. E., Murchison, N. D., Tabin, C. J. & Schweitzer, R. Generation of transgenic tendon reporters, ScxGFP and ScxAP, using regulatory elements of the scleraxis gene. Dev. Dyn. 236, 1677–1682 (2007).

100. Schweitzer, R. et al. Analysis of the tendon cell fate using Scleraxis, a specific marker for tendons and ligaments. Dev. Camb. Engl. 128, 3855–3866 (2001).

101. Mezu-Ndubuisi, O. J. & Maheshwari, A. Role of macrophages in fetal development and perinatal disorders. Pediatr. Res. 90, 513–523 (2021).

102. Epelman, S., Lavine, K. J. & Randolph, G. J. Origin and Functions of Tissue Macrophages. Immunity 41, 21–35 (2014).

103. Muscat, S., Nichols, A. E. C., Gira, E. & Loiselle, A. E. CCR2 is expressed by tendon resident macrophage and T cells, while CCR2 deficiency impairs tendon healing via blunted involvement of tendon-resident and circulating monocytes/macrophages. FASEB J. 36, e22607 (2022).

104. Wynn, T. A., Chawla, A. & Pollard, J. W. Origins and Hallmarks of Macrophages: Development, Homeostasis, and Disease. Nature 496, 445–455 (2013).

105. Bielefeld, K. A., Amini-Nik, S. & Alman, B. A. Cutaneous wound healing: recruiting developmental pathways for regeneration. Cell. Mol. Life Sci. 70, 2059– 2081 (2013).

106. Little, M. H. & Kairath, P. Does Renal Repair Recapitulate Kidney Development? J. Am. Soc. Nephrol. 28, 34 (2017).

107. Matsubayashi, Y. & Millard, T. H. Developmental Models for Wound Healing. In *eLS* (John Wiley & Sons, Ltd, 2013). doi:10.1002/9780470015902.a0021306.

108. Almekinders, L. C. & Deol, G. The effects of aging, antiinflammatory drugs, and ultrasound on the in vitro response of tendon tissue. Am. J. Sports Med. 27, 417– 421 (1999).

109. Delabastita, T., Bogaerts, S. & Vanwanseele, B. Age-related changes in achilles tendon stiffness and impact on functional activities: A systematic review and meta-analysis. J. Aging Phys. Act. 27, 116–127 (2019).

110. Ippolito, E., Natali, P. G., Postacchini, F., Accinni, L. & De Martino, C. Morphological, immunochemical, and biochemical study of rabbit Achilles tendon at various ages. J. Bone Jt. Surg. - Ser. A 62, 583–598 (1980).

111. Kannus, P., Paavola, M. & Józsa, L. Aging and Degeneration of Tendons. in Tendon Injuries: Basic Science and Clinical Medicine (eds. Maffulli, N., Renström, P. & Leadbetter, W. B.) 25 (Springer, 2005). doi:10.1007/1-84628-050-8_4.

112. Nakagawa, Y., Majima, T. & Nagashima, K. Effect of ageing on ultrastructure of slow and fast skeletal muscle tendon in rabbit Achilles tendons. Acta Physiol. Scand. 152, 307–313 (1994).

113. Strocchi, R. et al. Human achilles tendon: Morphological and morphometric variations as a function of age. Foot Ankle 12, 100–104 (1991).

114. Birk, D. E. & Trelstad, R. L. Extracellular compartments in tendon morphogenesis: collagen fibril, bundle, and macroaggregate formation. J. Cell Biol. 103, 231–240 (1986).

115. Humphries, S. M., Lu, Y., Canty, E. G. & Kadler, K. E. Active Negative Control of Collagen Fibrillogenesis in Vivo: INTRACELLULAR CLEAVAGE OF THE TYPE I PROCOLLAGEN PROPEPTIDES IN TENDON FIBROBLASTS WITHOUT INTRACELLULAR FIBRILS *. J. Biol. Chem. 283, 12129–12135 (2008).

116. Heinemeier, K. M., Schjerling, P., Heinemeier, J., Magnusson, S. P. & Kjaer, M. Lack of tissue renewal in human adult Achilles tendon is revealed by nuclear bomb 14C. FASEB J. 27, 2074–2079 (2013).

117. Heinemeier, K. M. et al. Carbon-14 bomb pulse dating shows that tendinopathy is preceded by years of abnormally high collagen turnover. FASEB J. 32, 4763– 4775 (2018).

118. Kalluri, R. & Weinberg, R. A. The basics of epithelial-mesenchymal transition. J. Clin. Invest. 119, 1420–1428 (2009).

119. Alhajj, M. & Goyal, A. Physiology, Granulation Tissue. in StatPearls (StatPearls Publishing, Treasure Island (FL), 2024).

120. Hope, M. & Saxby, T. S. Tendon Healing. Foot Ankle Clin. 12, 553–567 (2007).

121. Bruns, J., Kampen, J., Kahrs, J. & Plitz, W. Achilles tendon rupture: Experimental results on spontaneous repair in a sheep-model. Knee Surg. Sports Traumatol. Arthrosc. 8, 364–369 (2000).

122. Grinstein, M. et al. A distinct transition from cell growth to physiological homeostasis in the tendon. eLife 8, (2019).

123. Mimpen, J., Paul, C., Network, T. S., Cribbs, A. & Snelling, S. Nuclei isolation from snap-frozen tendon tissue for single nucleus RNA Sequencing. (2021).

124. Baldwin, M. J., Cribbs, A. P., Guilak, F. & Snelling, S. J. B. Mapping the musculoskeletal system one cell at a time. Nat. Rev. Rheumatol. 17, 247–248 (2021).

125. Baldwin, M. et al. A roadmap for delivering a human musculoskeletal cell atlas. Nat. Rev. Rheumatol. 19, 738–752 (2023).

126. Cribbs, A. P. cribbslab/scflow. cribbslab (2024).

127. Sullivan, D. K. et al. kallisto, bustools and kb-python for quantifying bulk, single-cell and single-nucleus RNA-seq. Nat. Protoc. 20, 587–607 (2025).

128. Amezquita, R. A. et al. Orchestrating single-cell analysis with Bioconductor. Nat. Methods 17, 137–145 (2020).

129. Hao, Y. et al. Integrated analysis of multimodal single-cell data. Cell 184, 3573–3587.e29 (2021).

130. McCarthy, D. J., Campbell, K. R., Lun, A. T. L. & Wills, Q. F. Scater: pre-processing, quality control, normalization and visualization of single-cell RNA-seq data in R. Bioinformatics 33, 1179–1186 (2017).

131. Germain, P.-L., Lun, A., Meixide, C. G., Macnair, W. & Robinson, M. D. Doublet identification in single-cell sequencing data using *scDblFinder*. Preprint at 10.12688/f1000research.73600.2 (2022).

132. Campbell, J., Yang, S., Wang, Z., Corbett, S. & Koga, Y. celda: CEllular Latent Dirichlet. 10.18129/B9.bioc.celda (2025).

133. Young, M. D. & Behjati, S. SoupX removes ambient RNA contamination from droplet-based single-cell RNA sequencing data. GigaScience 9, giaa151 (2020).

134. Korsunsky, I. et al. Fast, sensitive and accurate integration of single-cell data with Harmony. Nat. Methods 16, 1289–1296 (2019).

135. Zheng, G. X. Y. et al. Massively parallel digital transcriptional profiling of single cells. Nat. Commun. 8, 14049 (2017).

136. Fleming, S. J. et al. Unsupervised removal of systematic background noise from droplet-based single-cell experiments using CellBender. Nat. Methods 20, 1323– 1335 (2023).

137. Bergen, V., Lange, M., Peidli, S., Wolf, F. A. & Theis, F. J. Generalizing RNA velocity to transient cell states through dynamical modeling. Nat. Biotechnol. 38, 1408–1414 (2020).

138. Wolf, F. A., Angerer, P. & Theis, F. J. SCANPY: large-scale single-cell gene expression data analysis. Genome Biol. 19, 15 (2018).

139. Heumos, L. et al. Best practices for single-cell analysis across modalities. Nat. Rev. Genet. 24, 550–572 (2023).

140. Ahlmann-Eltze, C. & Huber, W. Comparison of transformations for single-cell RNA-seq data. Nat. Methods 20, 665–672 (2023).

141. Luecken, M. D. et al. Benchmarking atlas-level data integration in single-cell genomics. Nat. Methods 19, 41–50 (2022).

142. Gayoso, A. et al. A Python library for probabilistic analysis of single-cell omics data. Nat. Biotechnol. 40, 163–166 (2022).

143. Mao, S., Zhang, Y., Seelig, G. & Kannan, S. CellMeSH: probabilistic cell-type identification using indexed literature. Bioinformatics 38, 1393–1402 (2022).

144. Raudvere, U. et al. g:Profiler: a web server for functional enrichment analysis and conversions of gene lists (2019 update). Nucleic Acids Res. 47, W191–W198 (2019).

145. Xu, C. et al. Probabilistic harmonization and annotation of single-cell transcriptomics data with deep generative models. Mol. Syst. Biol. 17, e9620 (2021).

146. Palla, G. et al. Squidpy: a scalable framework for spatial omics analysis. Nat. Methods 19, 171–178 (2022).

147. Heumos, L., Litinetskaya, A. & Hediyeh-Zadeh, S. 16. Differential gene expression analysis. in.

148. Lemaître, G., Nogueira, F. & Aridas, C. K. Imbalanced-learn: A Python Toolbox to Tackle the Curse of Imbalanced Datasets in Machine Learning. J. Mach. Learn. Res. 18, 1–5 (2017).

149. Pedregosa, F. et al. Scikit-learn: Machine Learning in Python. J. Mach. Learn. Res. 12, 2825–2830 (2011).

150. CZI Single-Cell Biology Program et al. CZ CELL×GENE Discover: A single-cell data platform for scalable exploration, analysis and modeling of aggregated data. 2023.10.30.563174 Preprint at 10.1101/2023.10.30.563174 (2023).

151. Nowotschin, S. et al. The emergent landscape of the mouse gut endoderm at single-cell resolution. Nature 569, 361–367 (2019).

152. Xu, C. et al. Automatic cell-type harmonization and integration across Human Cell Atlas datasets. Cell 186, 5876–5891.e20 (2023).

153. Aibar, S. et al. SCENIC: single-cell regulatory network inference and clustering. Nat. Methods 14, 1083–1086 (2017).

154. Szklarczyk, D. et al. The STRING database in 2023: protein–protein association networks and functional enrichment analyses for any sequenced genome of interest. Nucleic Acids Res. 51, D638–D646 (2023).

155. Büttner, M., Ostner, J., Müller, C. L., Theis, F. J. & Schubert, B. scCODA is a Bayesian model for compositional single-cell data analysis. Nat. Commun. 12, 6876 (2021).

156. Bankhead, P. et al. QuPath: Open source software for digital pathology image analysis. Sci. Rep. 7, 16878 (2017).

157. Schmidt, U., Weigert, M., Broaddus, C. & Myers, G. Cell Detection with Star-convex Polygons. in vol. 11071 265–273 (2018).

158. Virtanen, P. et al. SciPy 1.0: fundamental algorithms for scientific computing in Python. Nat. Methods 17, 261–272 (2020).

159. Seabold, S. & Perktold, J. Statsmodels: Econometric and Statistical Modeling with Python. scipy (2010) doi:10.25080/Majora-92bf1922-011.

160. Terpilowski, M. A. scikit-posthocs: Pairwise multiple comparison tests in Python. J. Open Source Softw. 4, 1169 (2019).

161. Love, M. I., Huber, W. & Anders, S. Moderated estimation of fold change and dispersion for RNA-seq data with DESeq2. Genome Biol. 15, 1–21 (2014).

162. Oliveros, J. C. Venny. An interactive tool for comparing lists with Venn’s diagrams. BioinfoGP Service (2015).

163. Schindelin, J., et al. Fiji: an open-source platform for biological-image analysis. Nat. Methods 9, 676–682 (2012).

