## Supplementary figures and images for "Cellular and molecular landscapes of human tendons across the lifespan revealed by spatial and single-cell transcriptomics"

### Figure S1-1

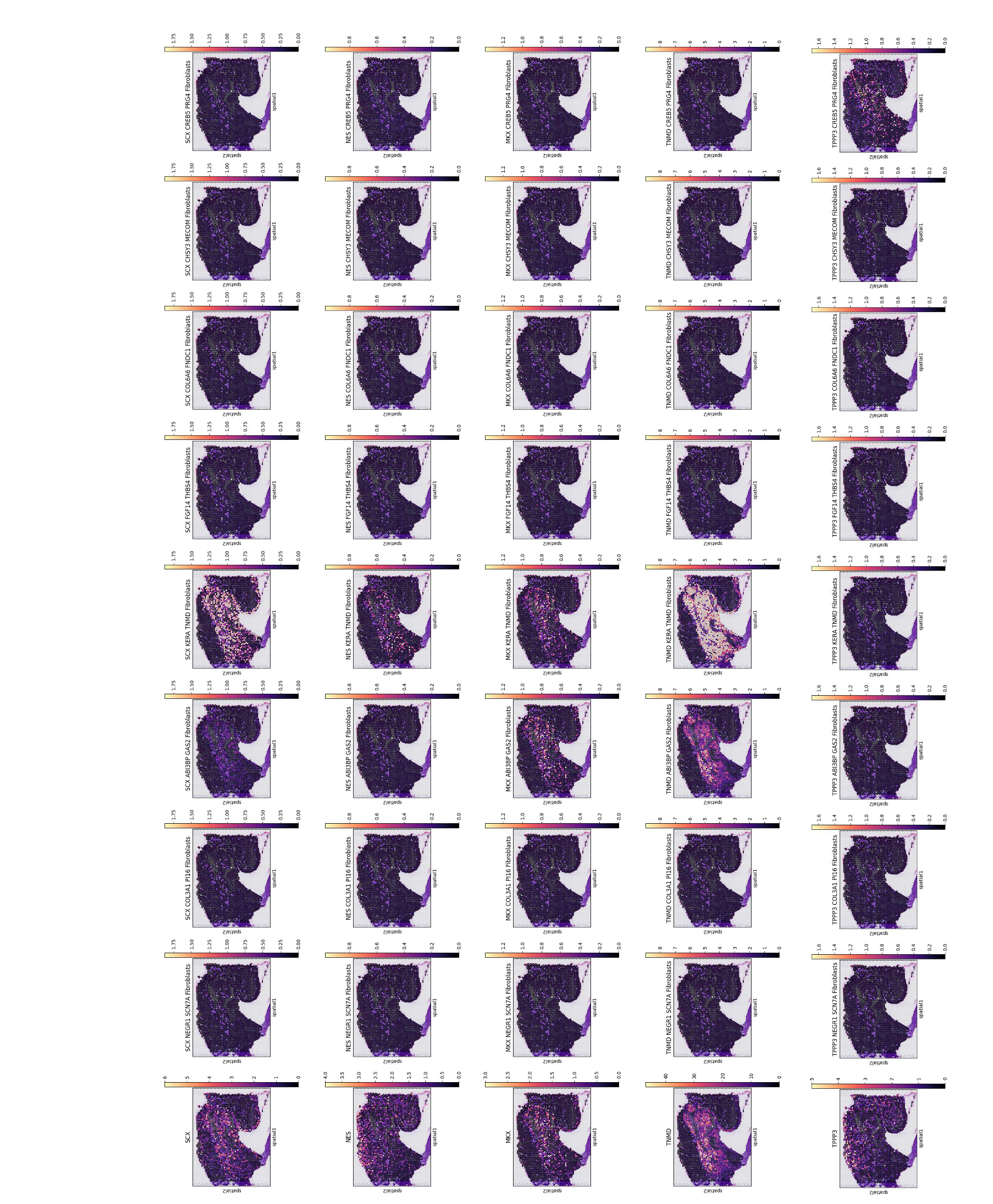

### Figure S1-2

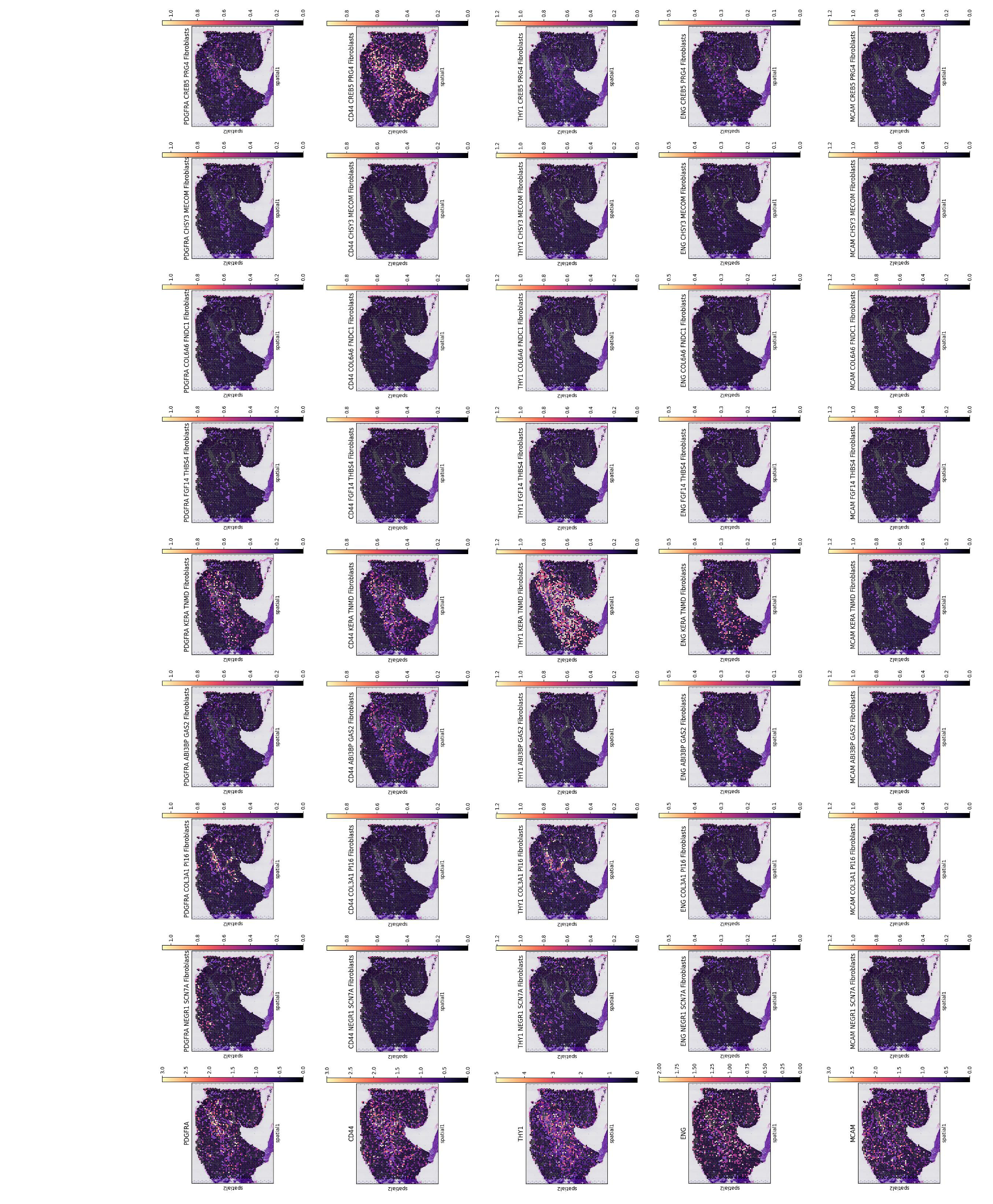

### Figure S2

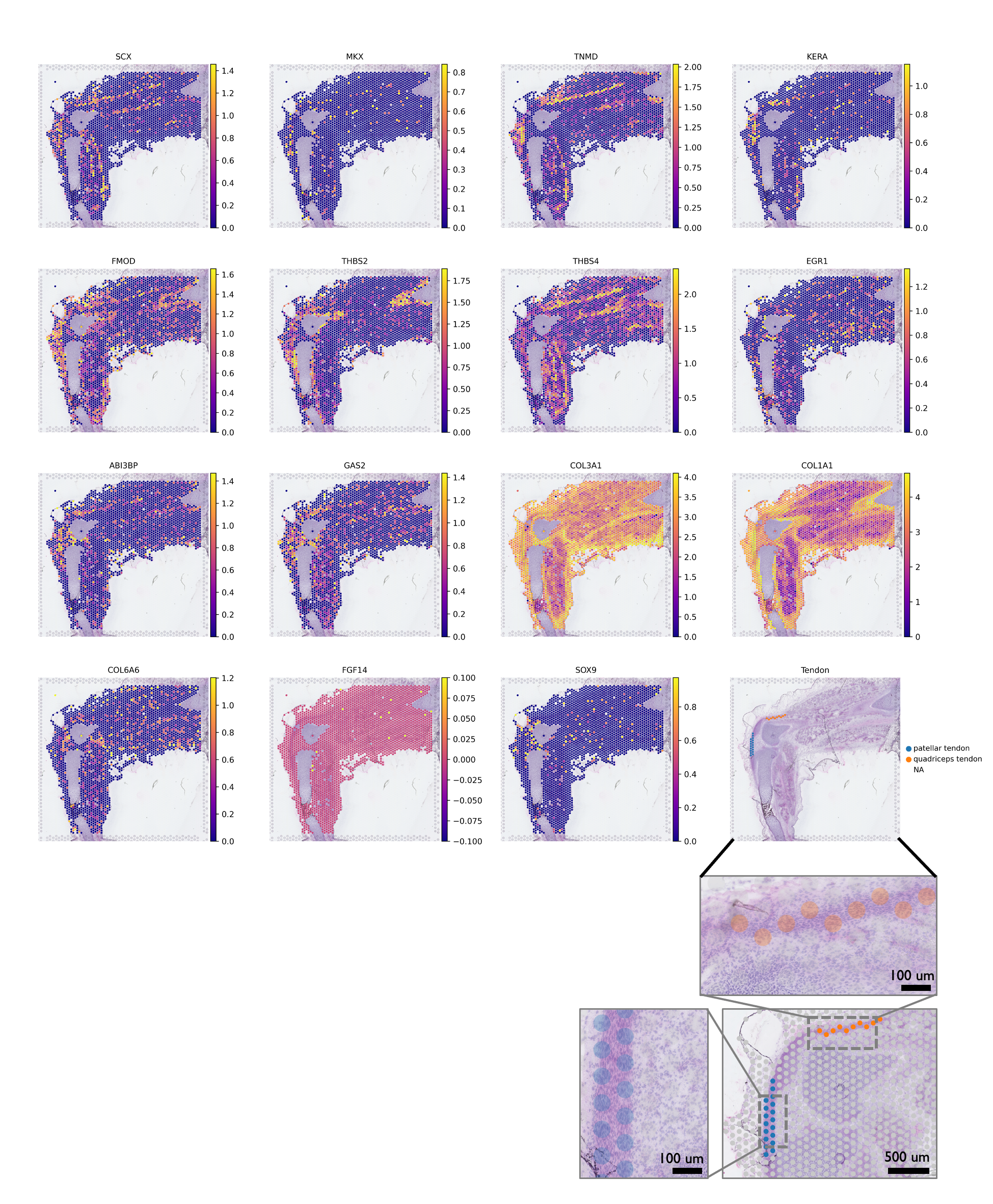

### Figure S3

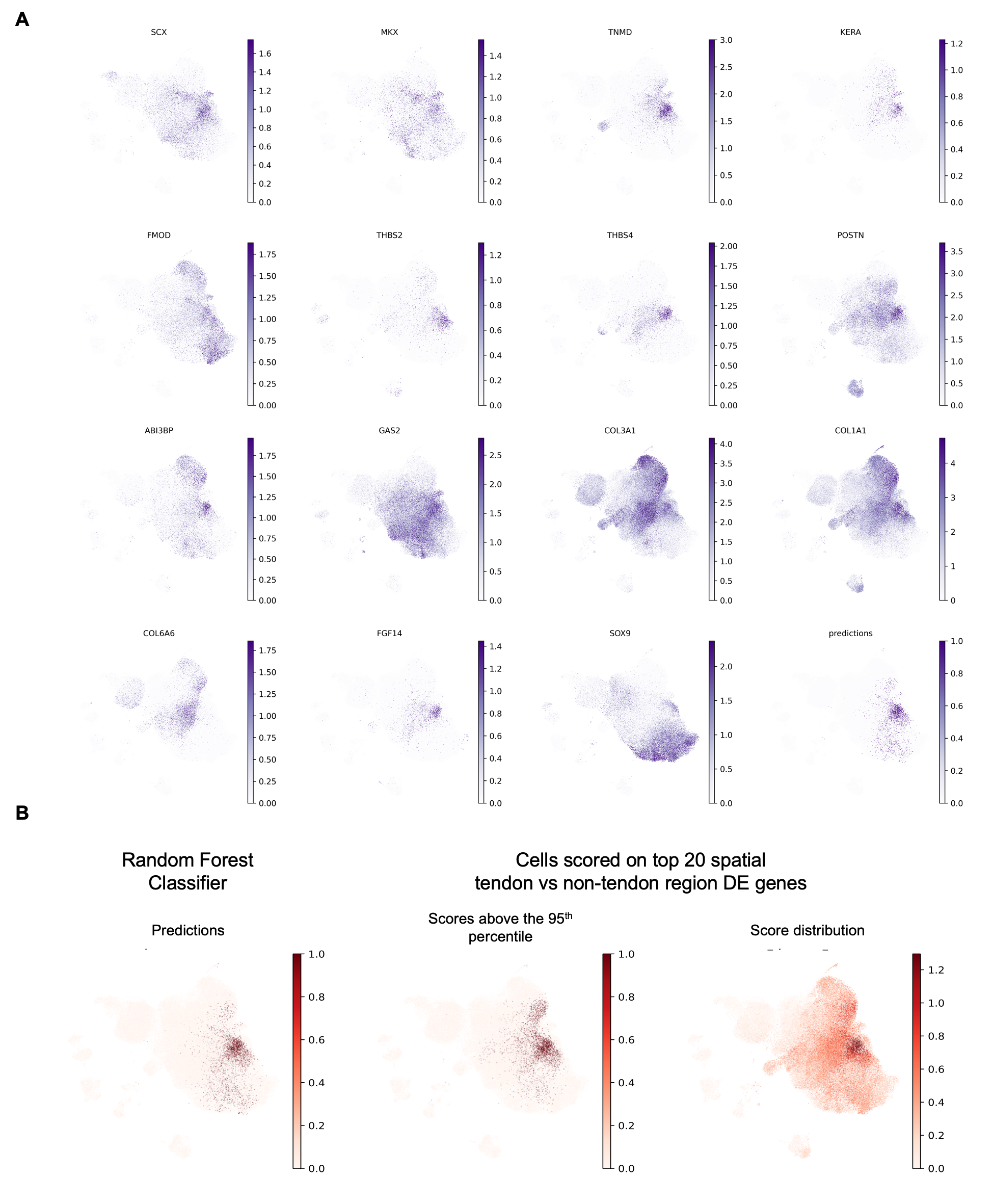

### Figure S4

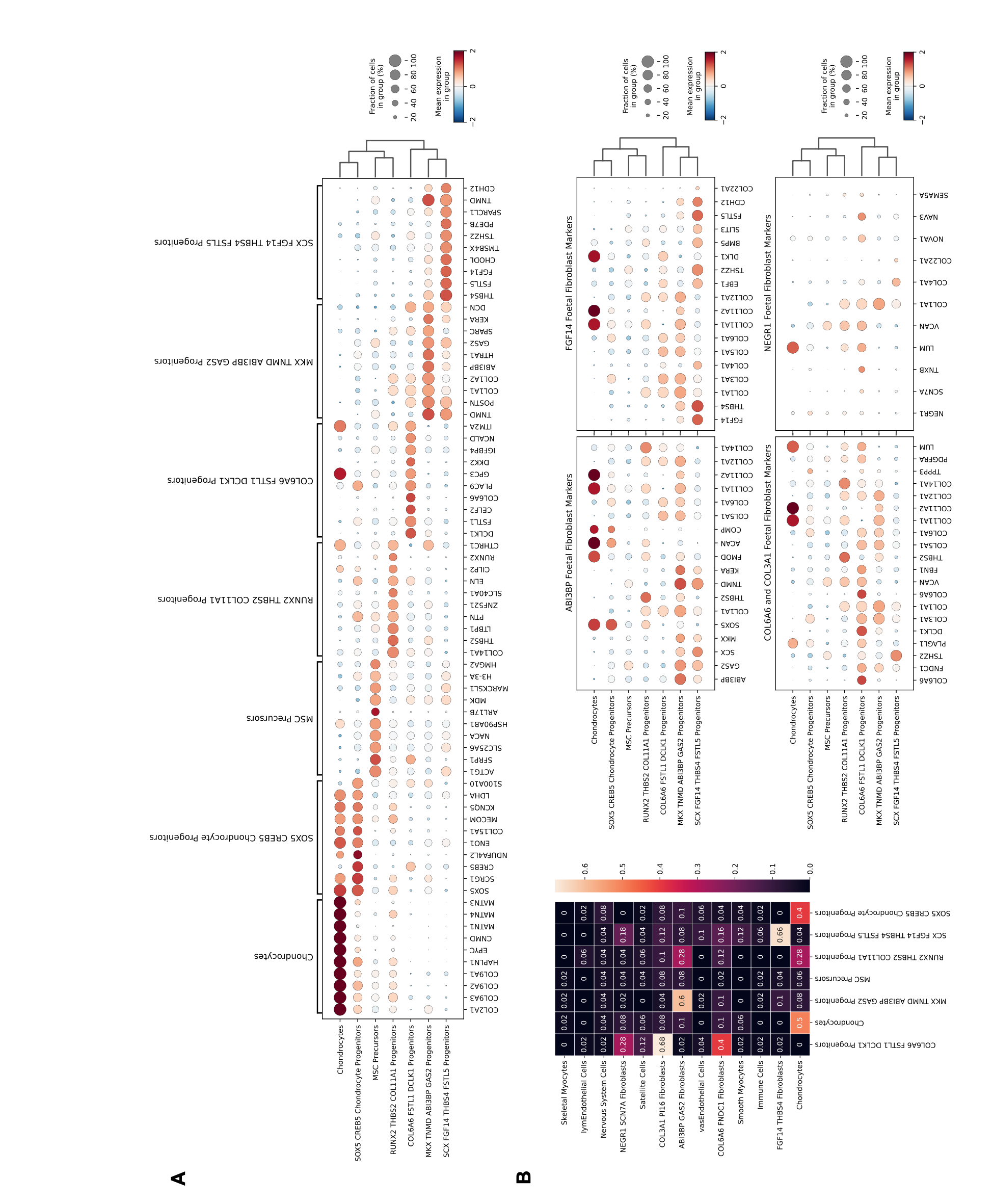

### Figure S5 (1)

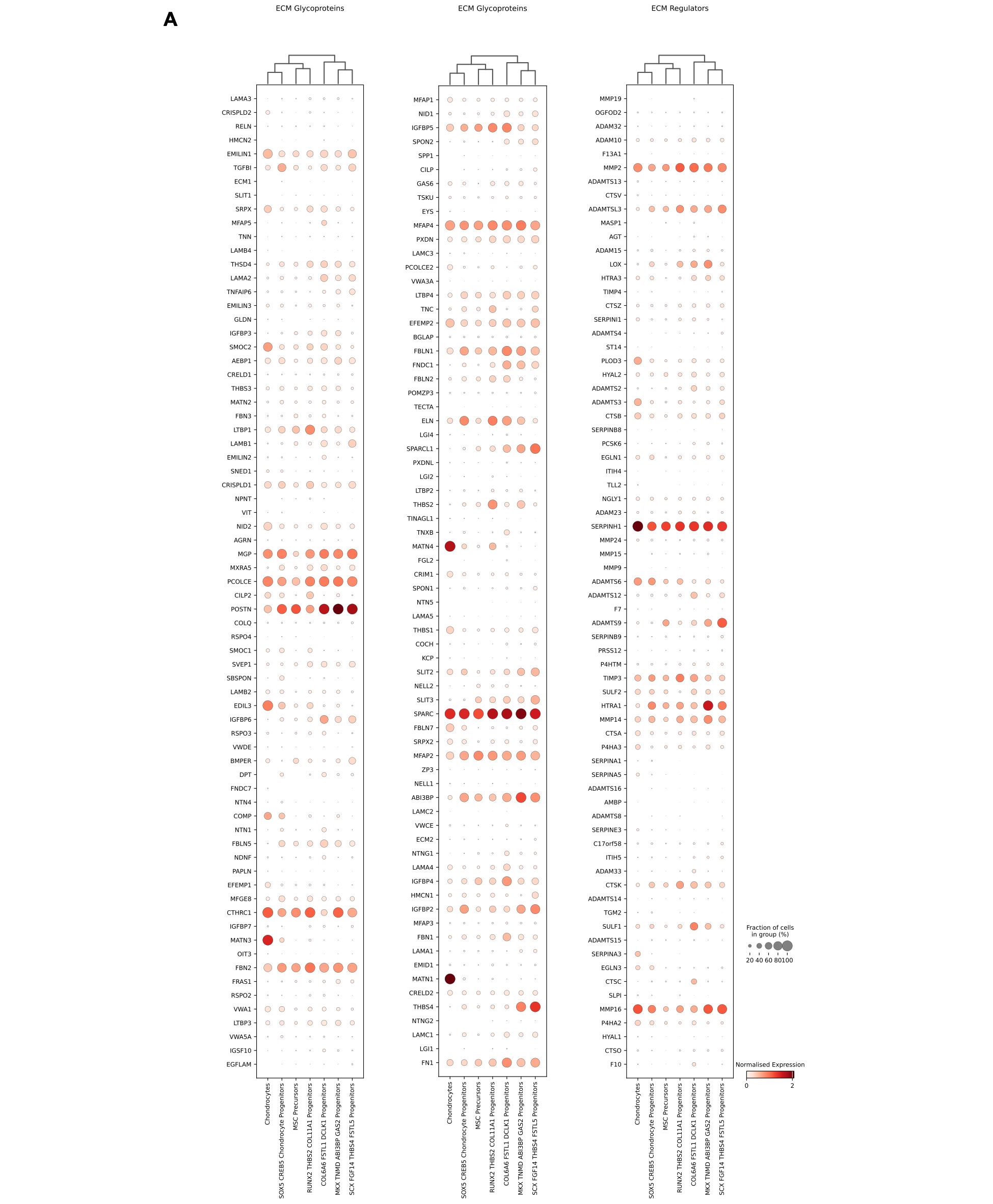

### Figure S5 (2)

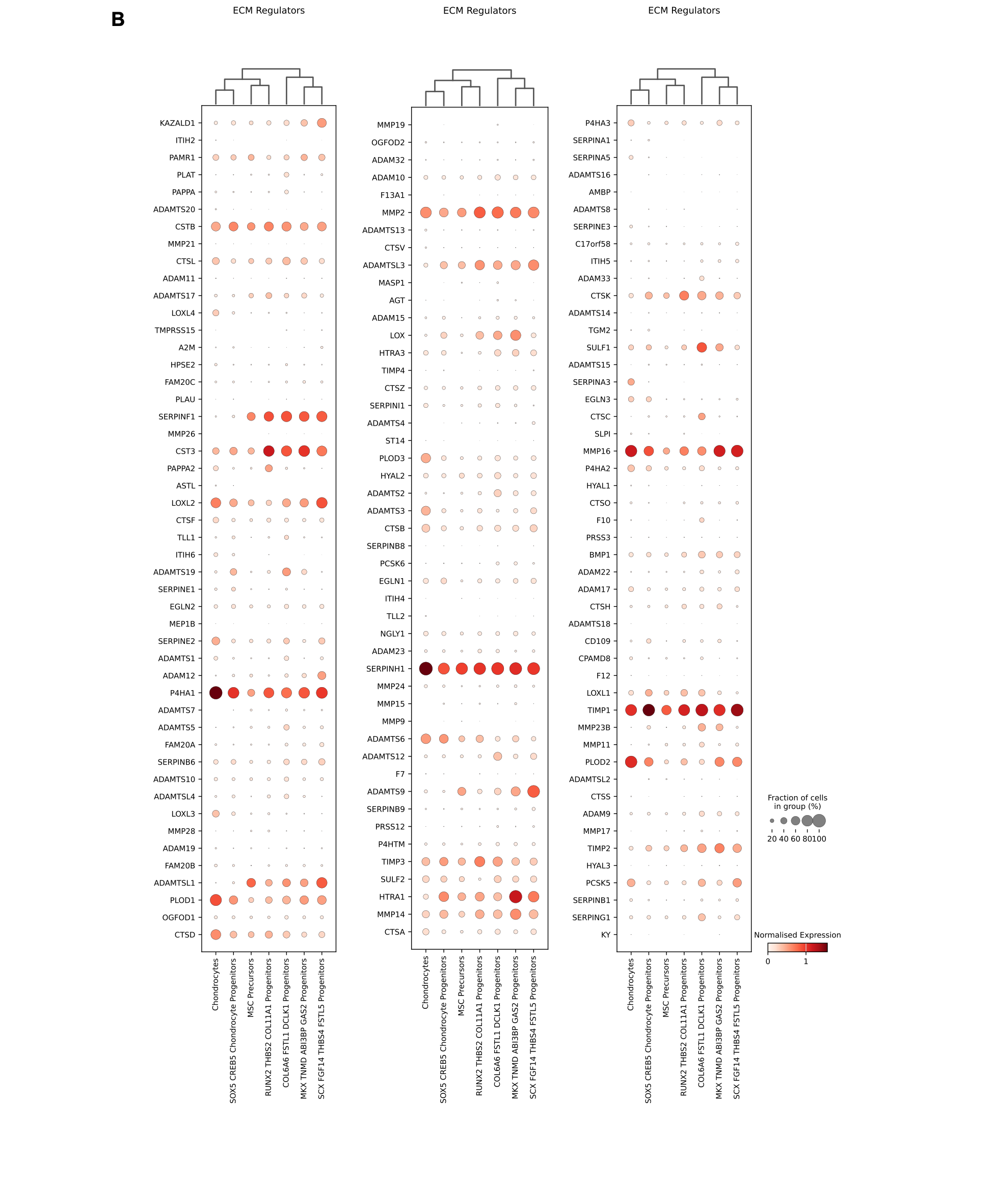

### Figure S5 (3)

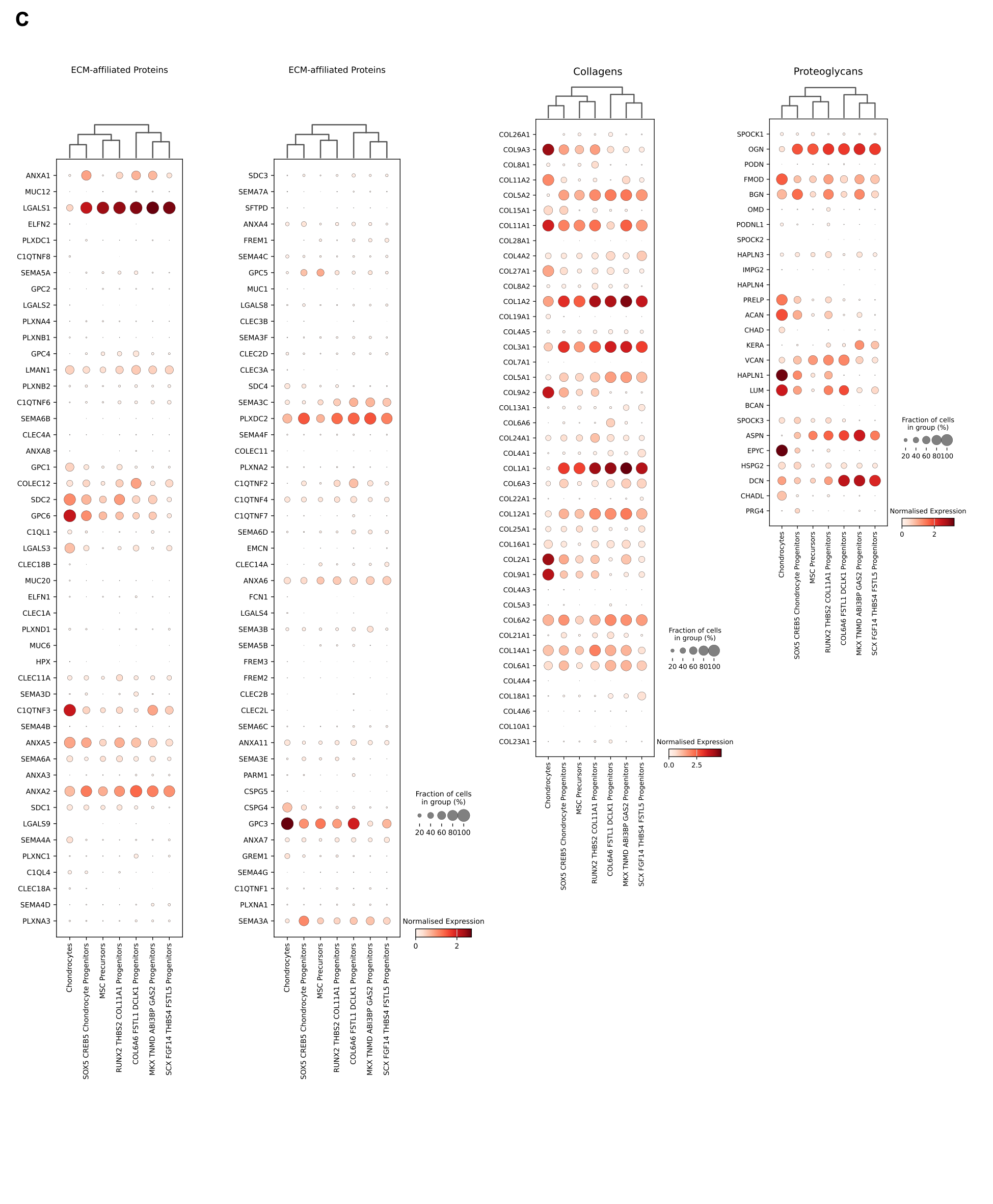

### Figure S6

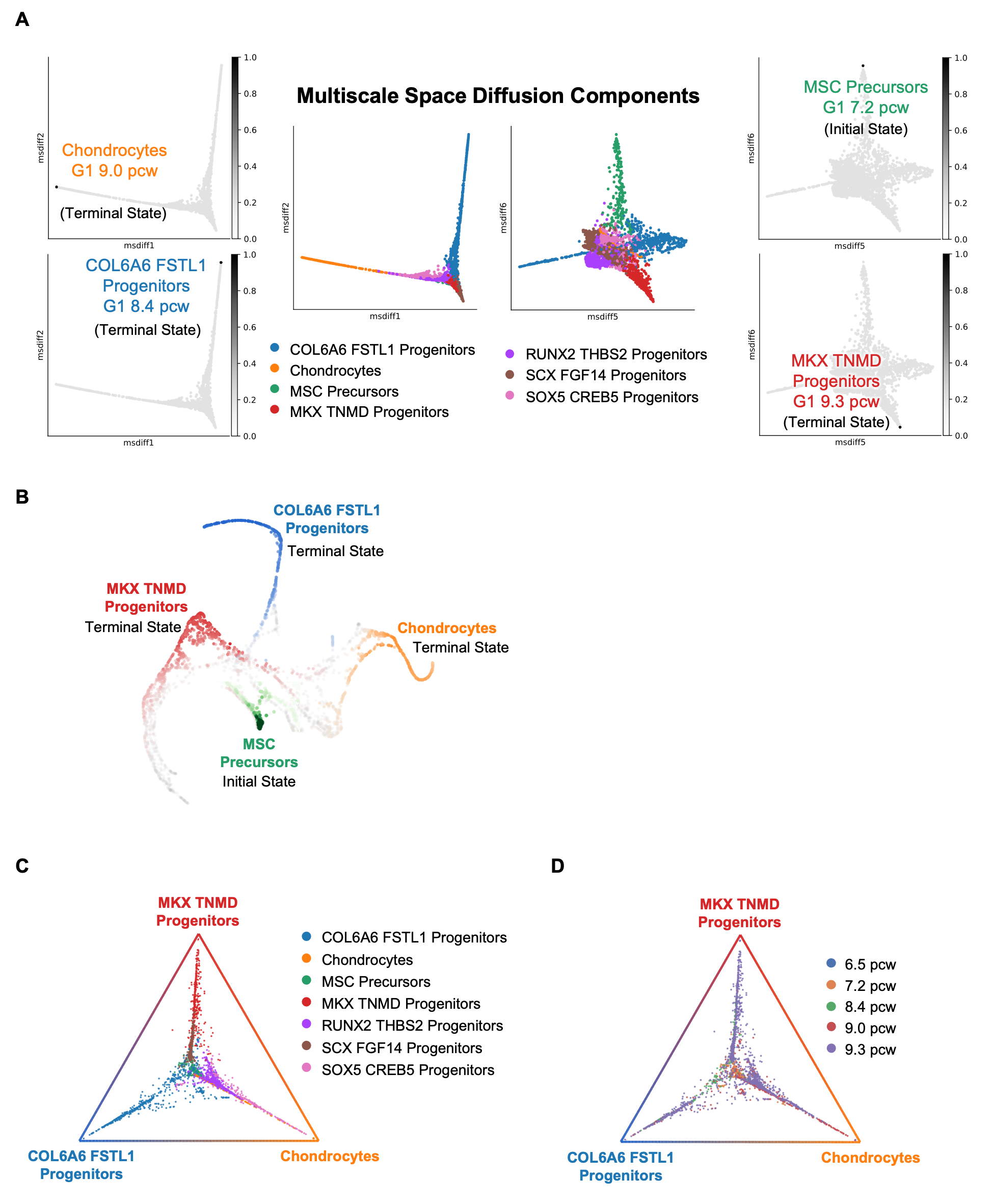

### Figure S7

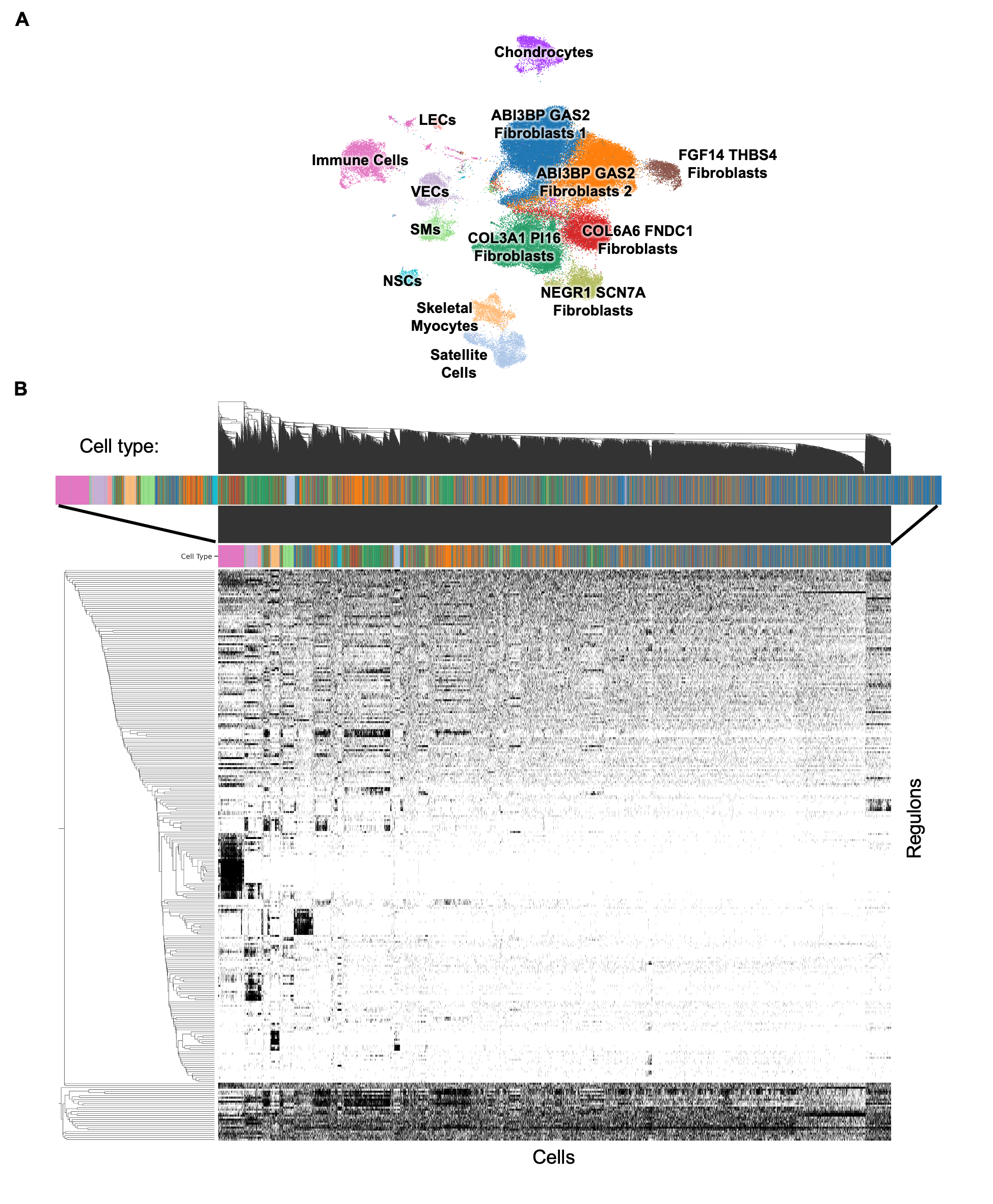

### Figure S8

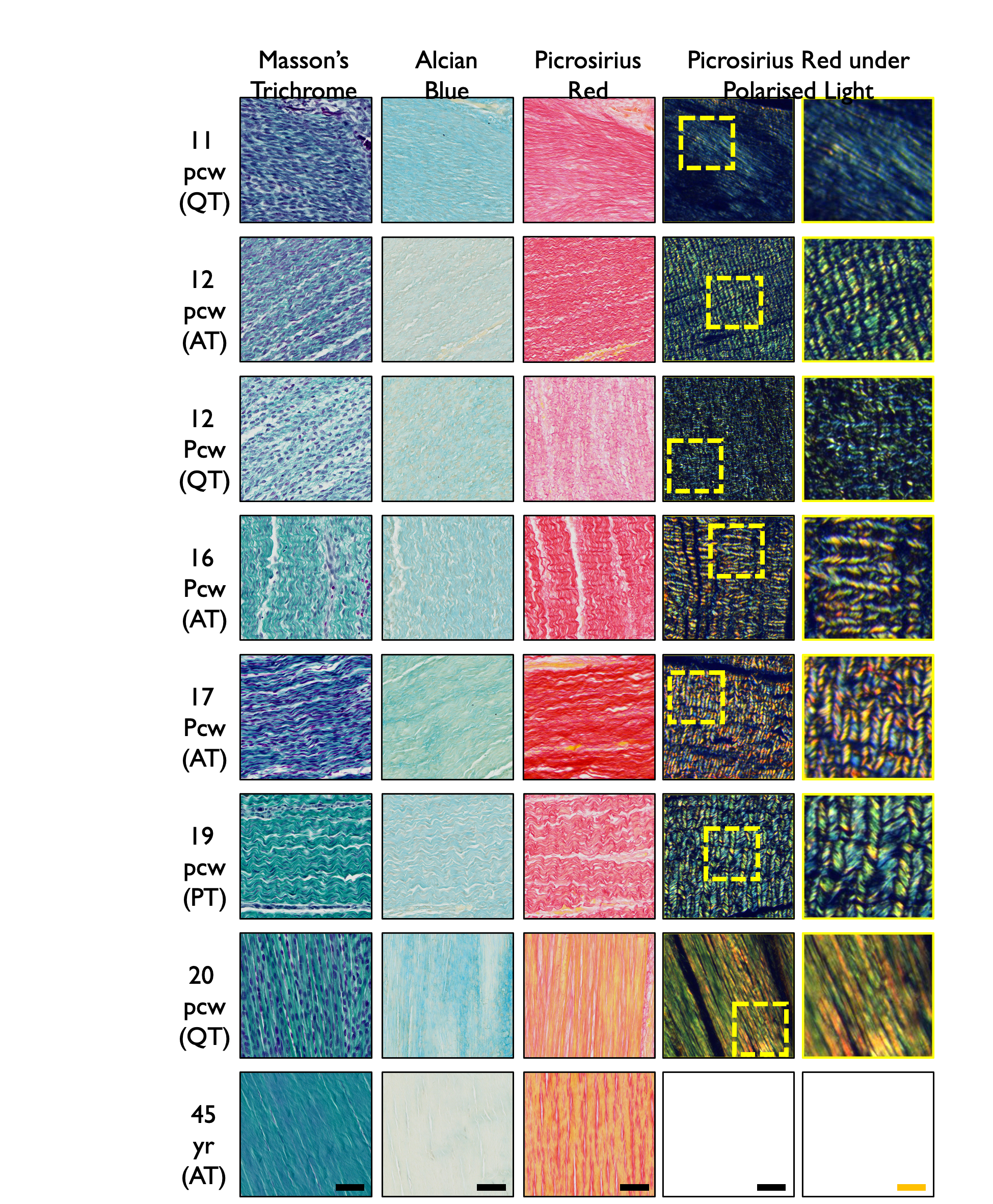

### Figure S9

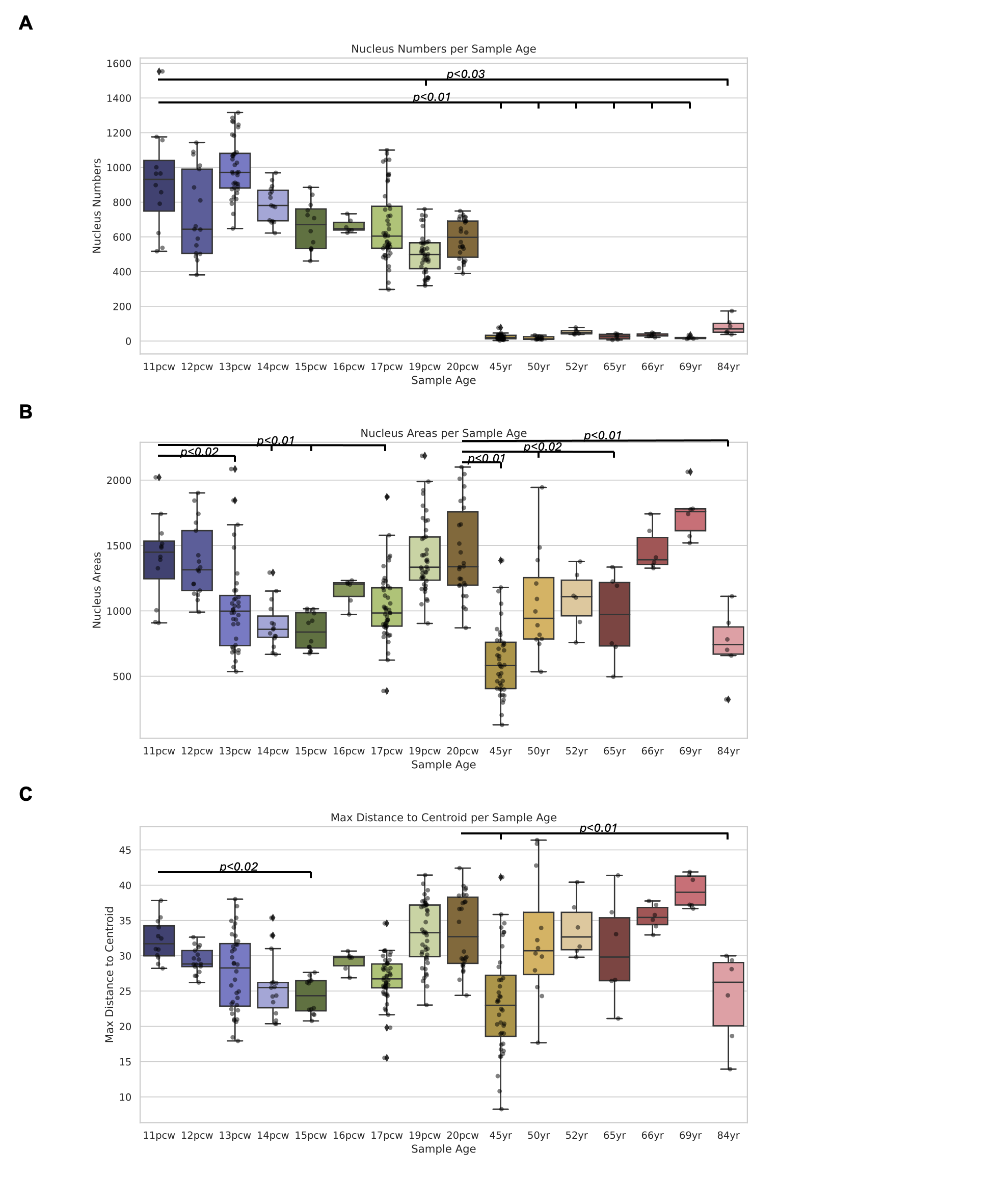

### Figure S10

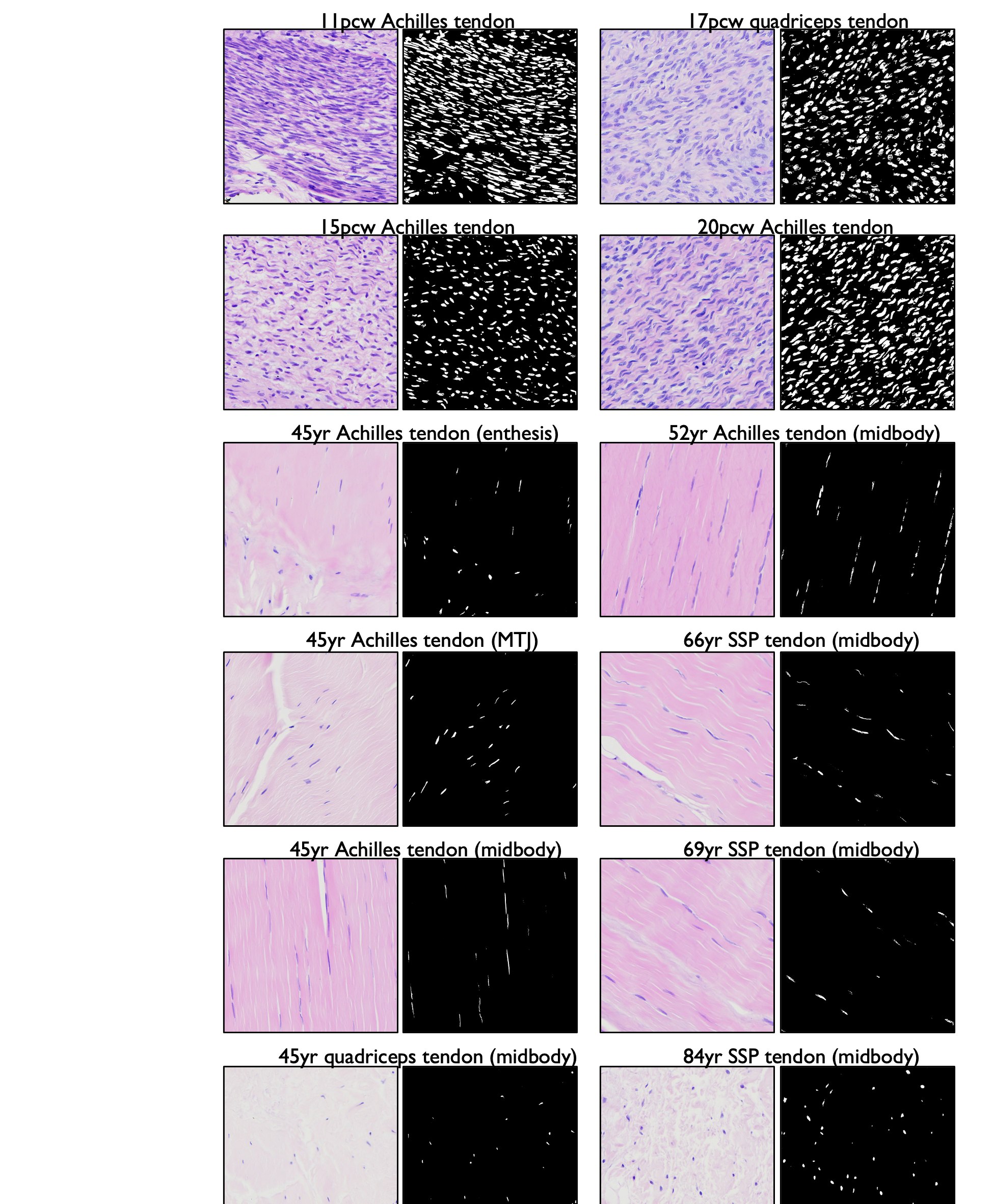

### Figure S11

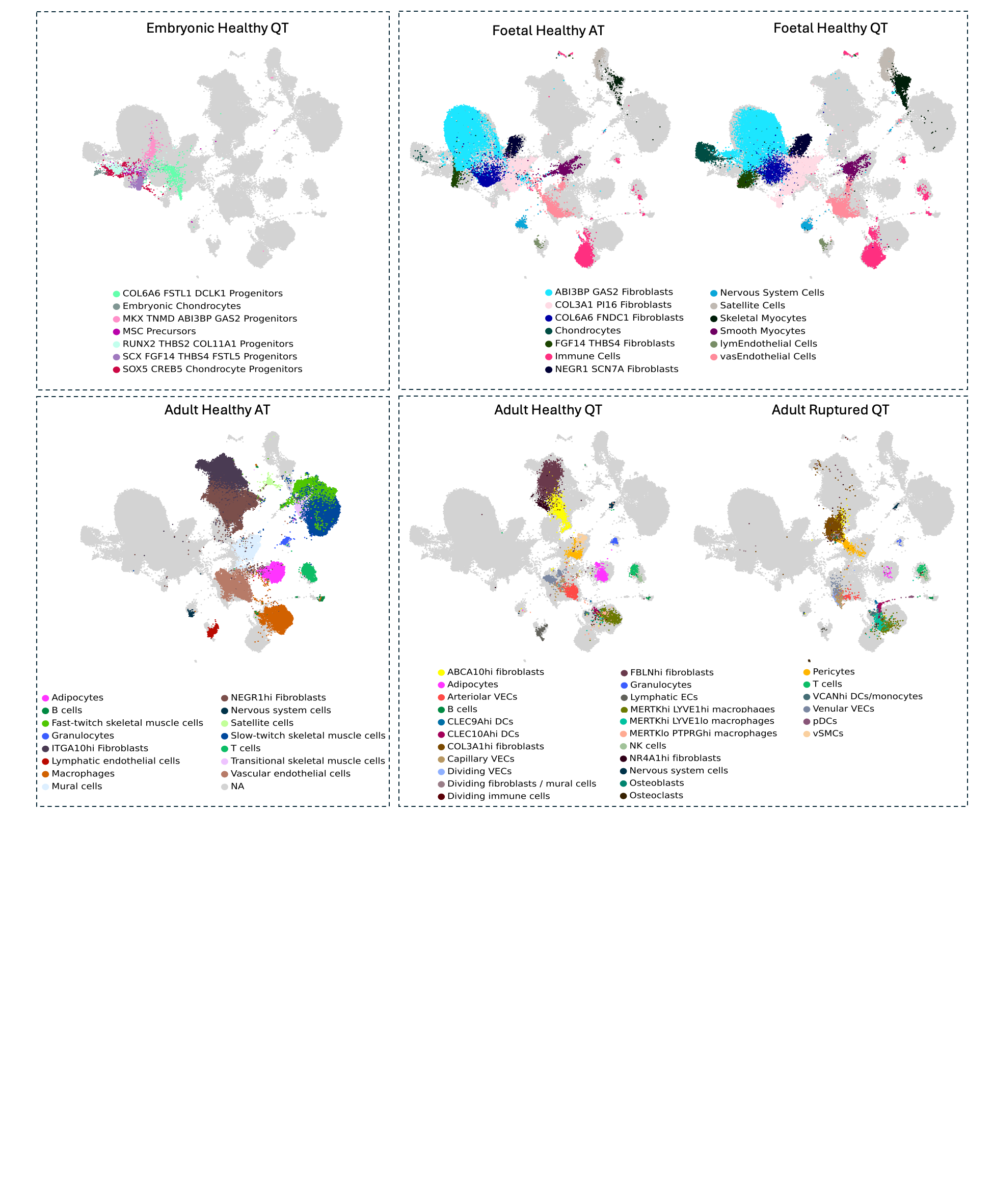

### Figure S12

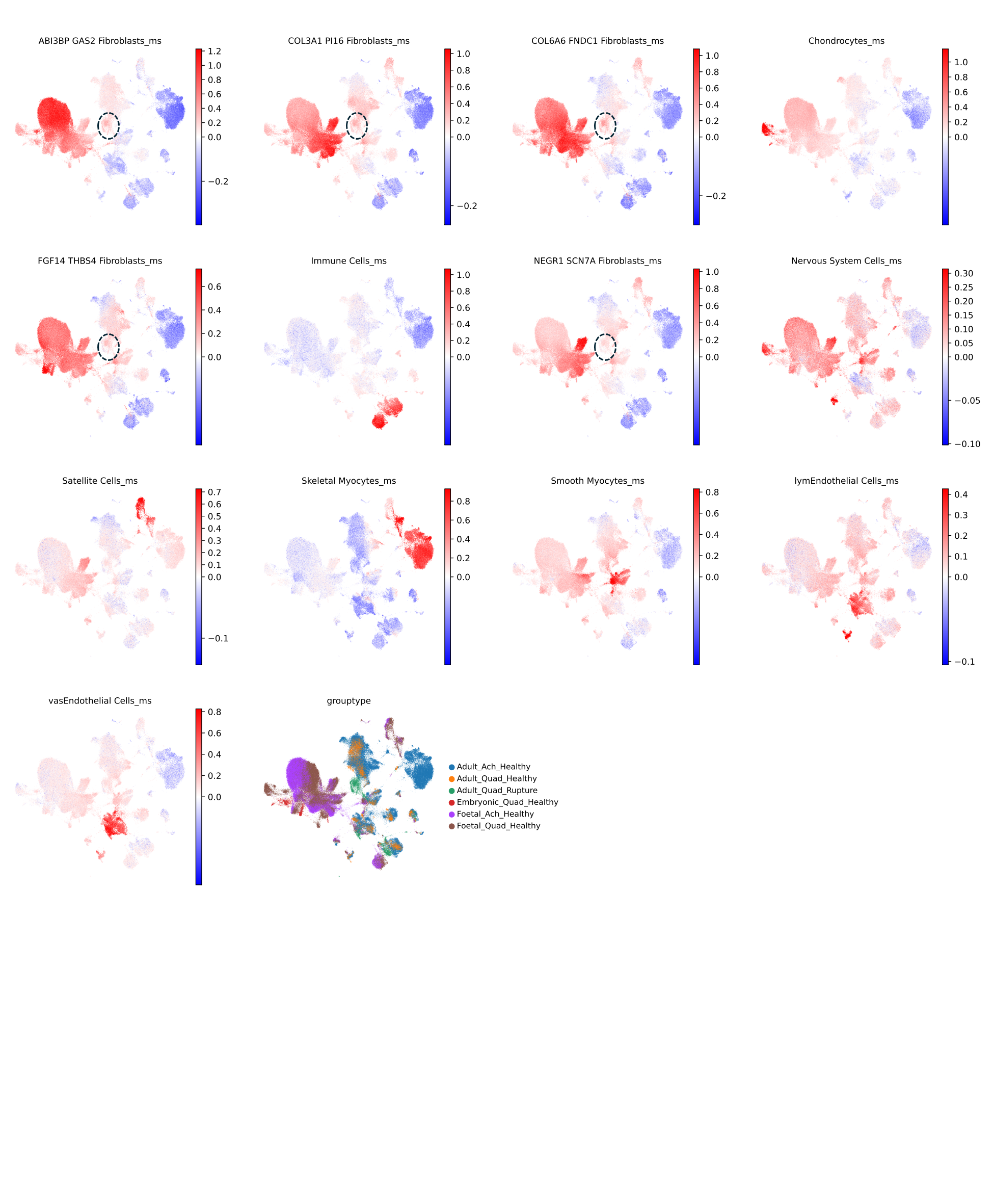

### Figure S13

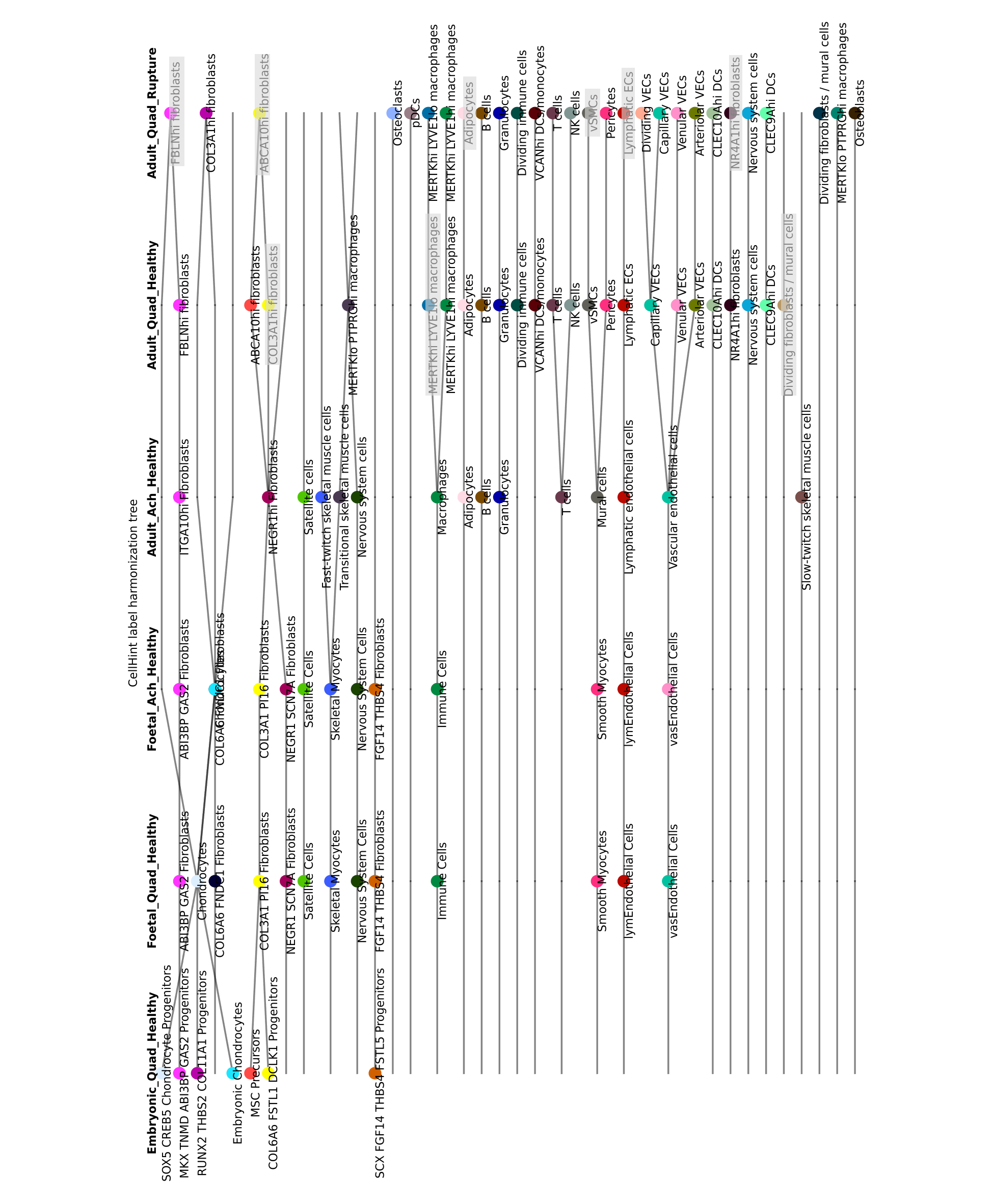

### Figure S14

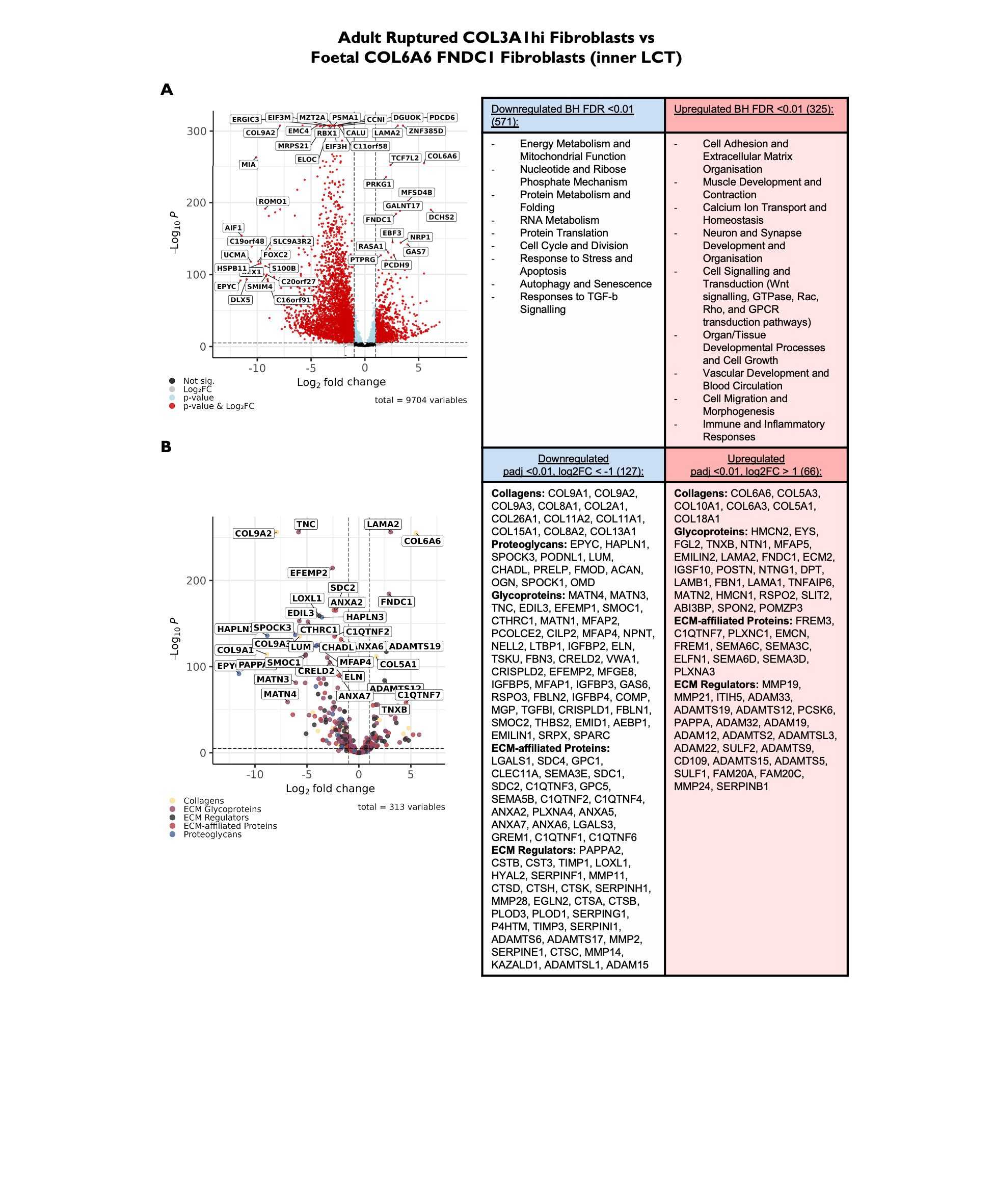

### Figure S15

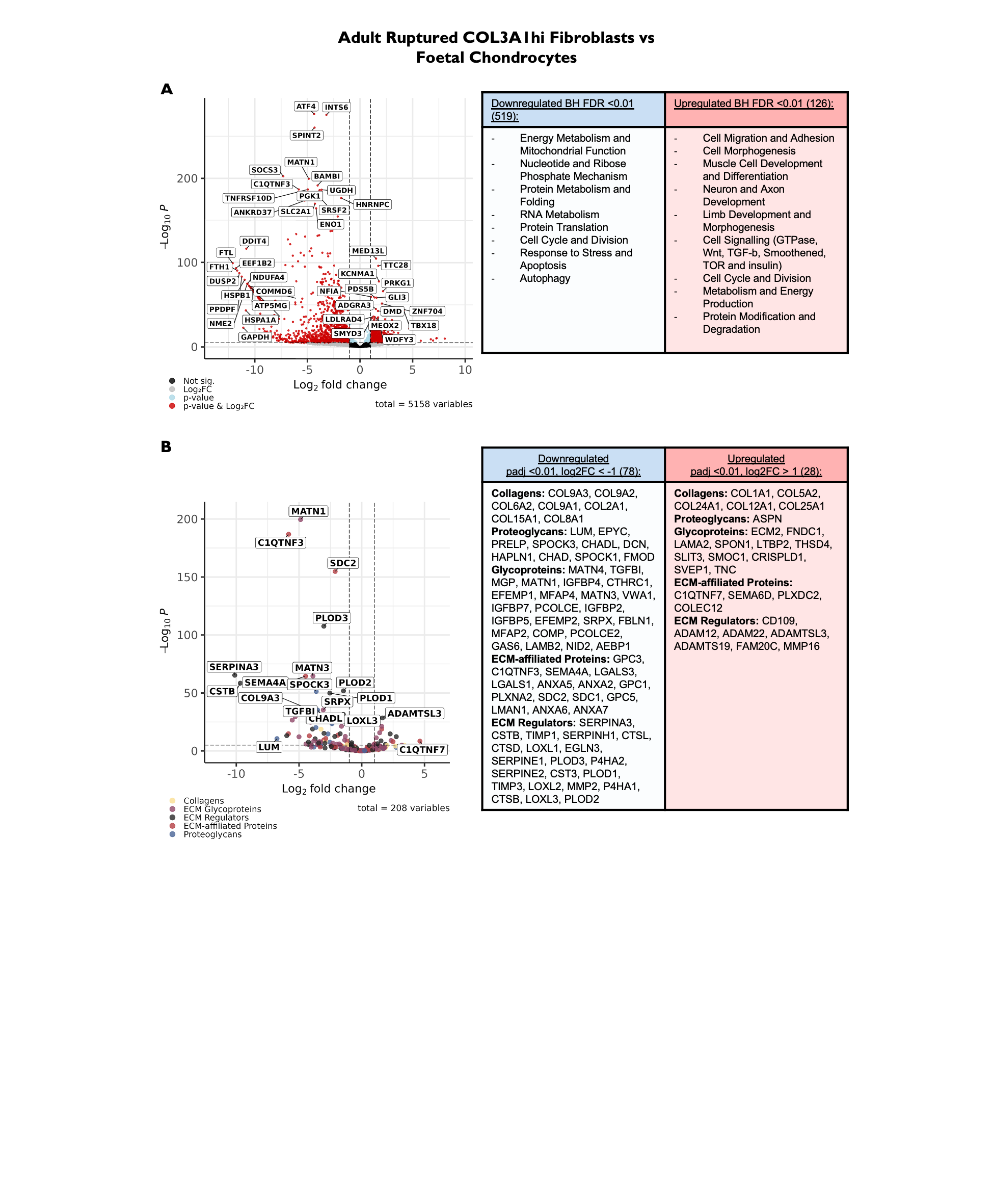

### Figure S16

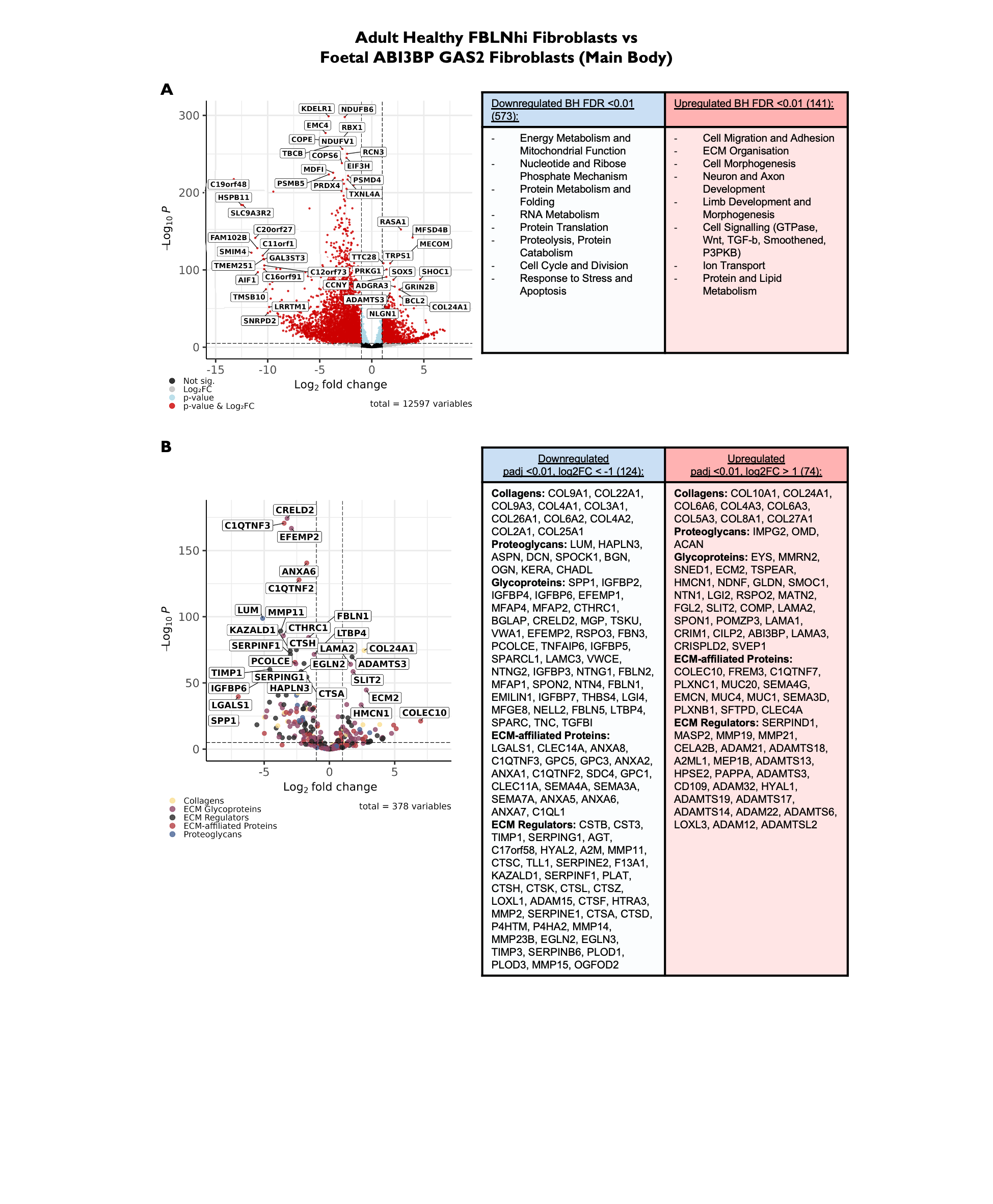

### Figure S17

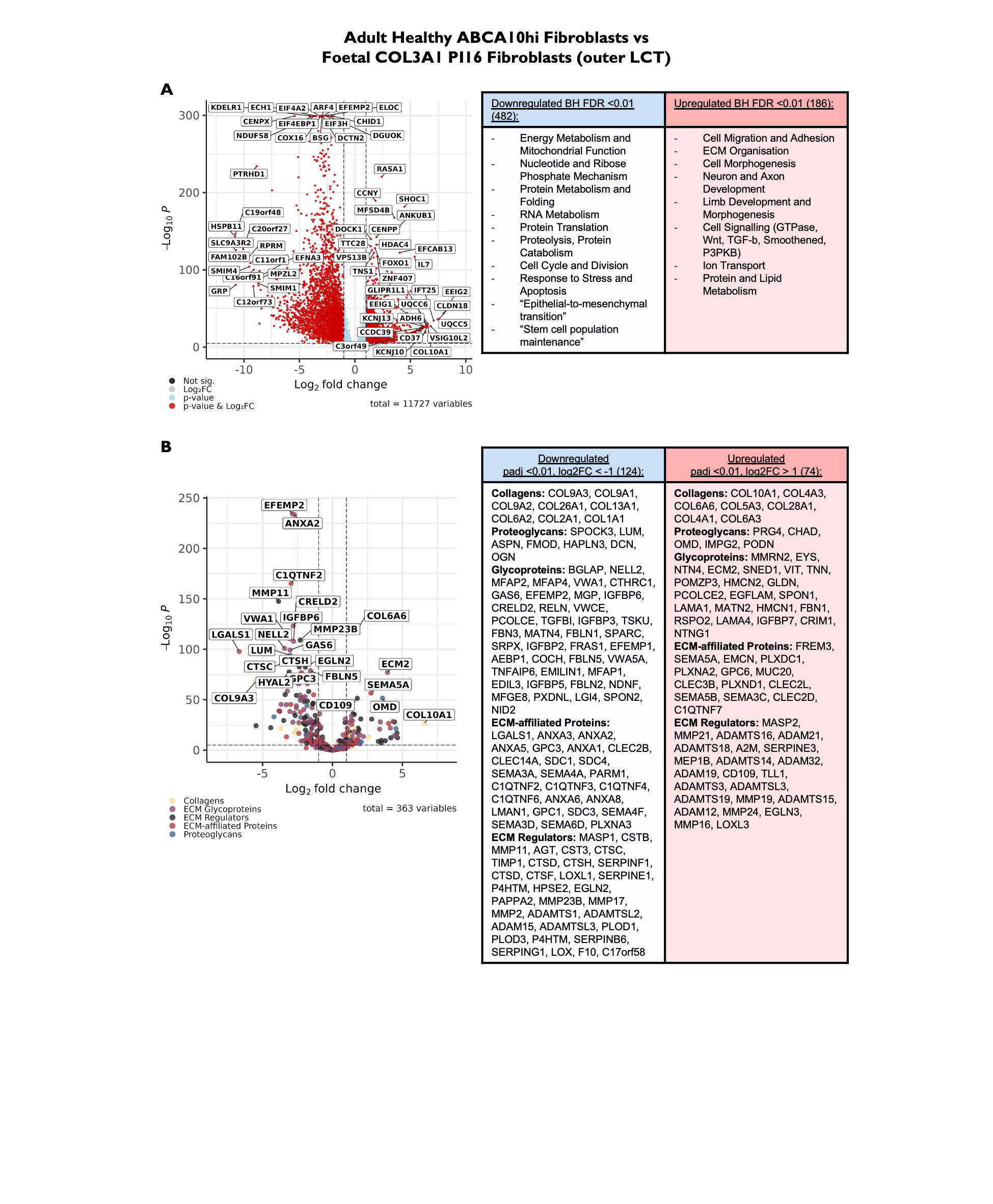
